# Resource diversity begets stability in complex ecosystems

**DOI:** 10.1101/2025.10.20.683391

**Authors:** Jamila Rowland-Chandler, Wenying Shou, Akshit Goyal

## Abstract

A fundamental paradox in ecology is the relationship between species diversity and ecosystem stability: May’s stability condition predicts that species diversity destabilises communities, yet many diverse ecosystems in nature are stable. Here, we show that this paradox can be resolved by explicitly considering resources, which May neglects. Specifically, May’s framework and the competitive exclusion principle jointly predict that resource diversity, which promotes species diversity, should destabilise communities. However, from computer simulations and analytical calculations using the finite-size cavity method, we find the opposite: resource diversity consistently generates stable, species-rich communities. Importantly, this stabilising effect disappears when resource dynamics is neglected (set to steady state). We also show that, contrary to the prevailing belief that interaction heterogeneity is always destabilising, different biological sources of heterogeneity have opposing effects on stability. Our work provides a solution to May’s paradox and demonstrates that resource dynamics are not just negligible background but are central drivers of ecosystem stability.

## Introduction

Ecological communities exhibit diverse dynamical behaviours, ranging from perturbation-resistant mono-stable states [1, 2] to multi-stability [3–5] to persistent fluctuations [6–12]. The dynamical behaviour of a community is critical for its functioning. For example, stability the ability of a community to return to the original steady state after a small perturbation can preserve the healthy metabolism of gut microbiomes, while pathogen invasion can drive the emergence of low-diversity, fluctuating diseased states [13, 14]. In contrast, fluctuations and turnover of soil communities can promote nutrient cycling [15–17] critical for crop and ecosystem productivity. Given the importance of ecological dynamics, ecologists have sought to understand how they arise from interactions between community members and their resource environment, such as competition for shared resources or cross-feeding [18–21].

Theoretical ecologists have attempted to understand the mechanisms driving stability and other dynamics. Traditionally, these studies used models where species interact directly (e.g., the generalised Lotka-Volterra model, or gLV), without accounting for the resource dynamics that mediate these interactions [22–27]. Using these “resource-implicit models”, Robert May and others suggested that species-rich communities with heterogeneous inter-species interactions are inherently unstable — known as “May’s stability condition” (Fig. 1 B, [8, 25–29] although exceptions exist [22]). However, these predictions often conflict with empirical evidence where species diversity can stabilise community biomass [30, 31] and species abundances [32, 33], and can increase resilience to perturbations [34]. This discrepancy between theory and observation may occur because resource-implicit models ignore the resource dynamics that mediate many species interactions [18, 21, 35–48]. Thus, it remains unclear whether resource-explicit models would reveal alternative stability relationships.

**Figure 1:**
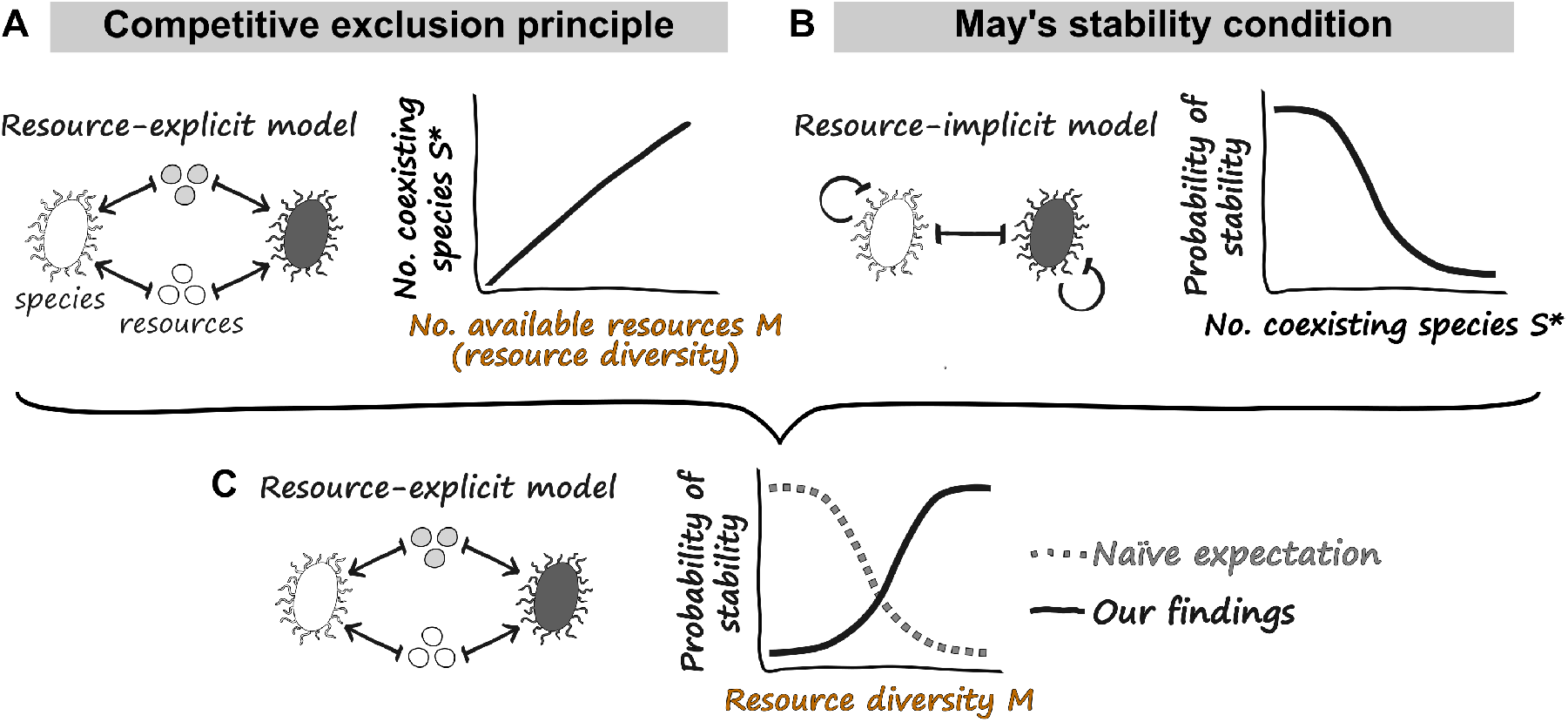
Does increasing the diversity of available resources destabilise community dynamics? **(A)** Competitive exclusion principle (derived from resource-explicit models): Increasing the number of available resources can support more co-existing species. **(B)** May’s stability condition (derived from resource-implicit models): Increasing the number of coexisting species should destabilise community dynamics. **(C)** Combining these principles, we naïvely expected that increasing the number of available resources would destabilise community dynamics by increasing species diversity (grey dotted line). However, we found the opposite (black line).

Here, we test a prediction about how resource diversity might affect stability: the competitive exclusion principle states that higher resource diversity can support more coexisting species [49–51] (Fig. 1 A). According to May’s stability condition, this higher species diversity should destabilise communities (Fig. 1 C). Specifically, we tested whether increasing the diversity of available resources, measured by resource pool size (*M*), destabilises community dynamics (Fig. 1 C) using a resource-explicit consumer-resource model. Contrary to expectations (Fig. 1 C), increasing the resource pool size increased species richness and stabilised community dynamics. This positive diversity-stability relationship was driven by resource dynamics themselves, because the stabilising effect disappeared when we reduced the consumer-resource model to a resource-implicit species-only model. Given this surprising result, we investigated another prevailing belief: that heterogeneity in species interactions always destabilises communities [25–27, 29, 52–55]. Our model couples resource consumption to species growth via a biologically intuitive yield conversion factor, which allowed us to generate interaction heterogeneity through two distinct biological mechanisms: variance in consumption 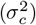 or variance in the yield conversion factor 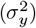 Again contrary to expectation, different sources of heterogeneity induced opposing stability transitions: although high variance in yield conversion 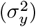 destabilised communities, high variance in consumption 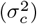 stabilised communities.

To understand these phenomena, we analytically derived the model’s stability condition, which requires the correlation between growth and consumption coefficients to exceed the square root of the species packing ratio (the ratio between the number of surviving species and the number of surviving resources) [52, 56, 57]. When the resource pool size *M* was small, reciprocity was smaller than 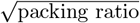, making communities unstable. Increasing the resource pool size increased both quantities, but reciprocity grew faster and ultimately exceeded 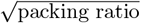, generating stable and species-rich communities. Increasing variance in consumption 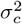 stabilised communities through a similar mechanism. In contrast, increasing variance in yield 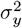 reduced reciprocity faster than reducing 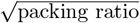, leading to instability. Together, our results demonstrate that resource dynamics can fundamentally alter the relationship between ecological diversity, interaction heterogeneity, and stability, and help reconcile theoretical predictions with empirical observations.

### The model

In our consumer-resource model, a pool of *S* consumer species grows by consuming a pool of *M* resources. The dynamics of resource *α* (*R*_*α*_) and consumer *i* (*N*_*i*_) are described by the following set of ordinary differential equations:

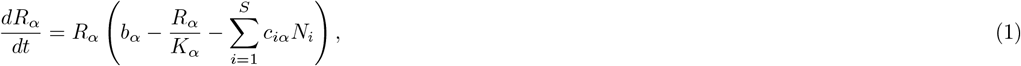

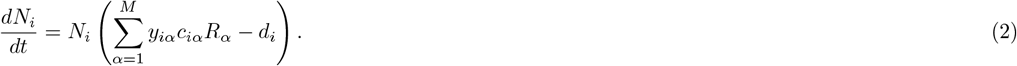

Here, resources (e.g., phytoplankton) grows logistically with *b*_*α*_ being the intrinsic growth rate and *K*_*α*_ being the carrying capacity of resource *α* (both set to 1 without loss of generality). Consumers (e.g., heterotrophic microbes) deplete resources, where *c*_*iα*_ is the “consumption coefficient” of resource *α* by consumer *i* (i.e., consumption rate per capita consumer per unit resource). Note that both resources and consumers can go extinct, and we use *M* ^∗^ and *S*^∗^ to represent the numbers of surviving resources and consumers, respectively. We assumed that all resources are substitutable (e.g., different types of carbon instead of carbon versus nitrogen), so they contribute additively to consumer growth rates (for a full list of assumptions and model details, see SI Appendix B). Consumer species *i* dies with a death rate of *d*_*i*_, and grows by converting consumed resources into biomass via *y*_*iα*_, which is the yield conversion factor for species *i* on resource *α*. The resulting per capita growth rate of species *i* from consuming resource *α* is thus given by *y*_*iα*_*c*_*iα*_*R*_*α*_, where we denote *y*_*iα*_*c*_*iα*_ as the “growth coefficient” (i.e., per capita growth rate per unit resource). Because yield conversion is consumer-specific, growth and consumption coefficients are *non-reciprocal*, meaning they are not perfectly correlated. This non-reciprocity is crucial for communities to display both stable and unstable dynamics in different parameter regimes [52, 57].

To understand how the resource pool size affects the diversity and stability of complex communities, we followed a long tradition of treating the large number of model parameters as random variables [24, 25, 27–29, 36, 55, 58–64]. Because the total resource consumption rate cannot continue to increase arbitrarily with resource pool size *M*, we assumed that the consumption coefficient for each resource decreases with resource pool size *M* (*c*_*iα*_ ∝ 1*/M*). Thus, we sampled each *c*_*iα*_ from a normal distribution with mean *µ*_*c*_*/M* and variance 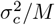 (Fig. 2 A). We sampled yield conversion factors *y*_*iα*_ from a normal distribution with mean *µ*_*y*_ and variance 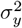 (Fig. 2 A). For complete details of the model parametrisation and simulations, see SI Appendices B and D. Our central results are robust to many details of parameter choices (e.g., the choice of distribution and values of consumption coefficients and yield conversion factors) as well as some details of the formulation of the model (SI Appendices E and G).

**Figure 2:**
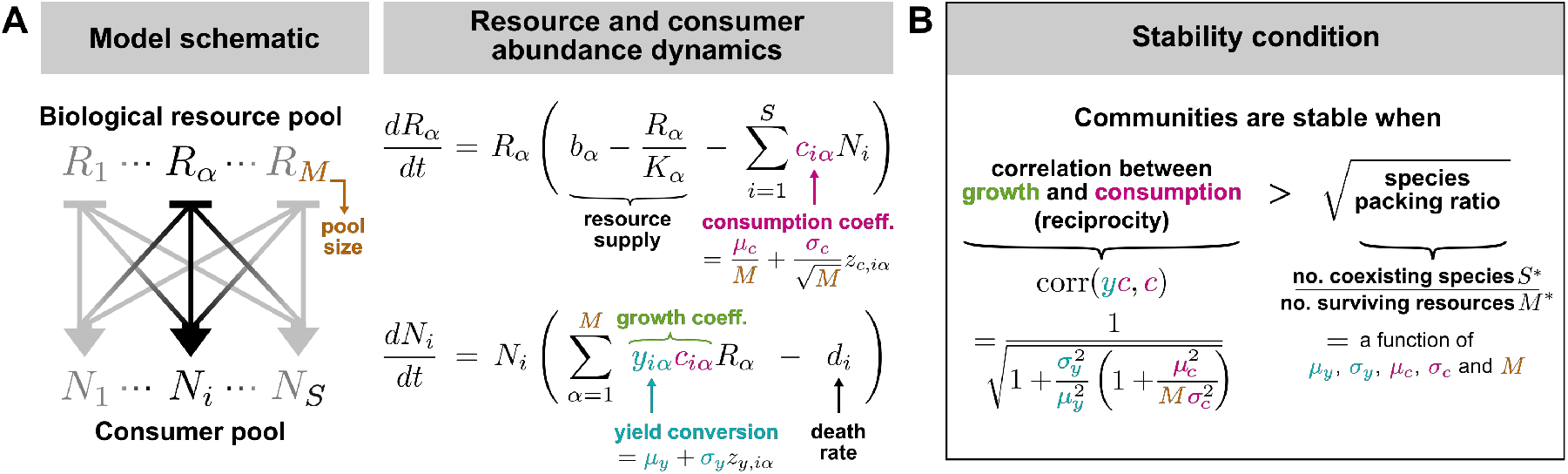
Our consumer-resource model and its stability condition. **(A)** Model schematic and consumer-resource dynamics. We sample yield conversion factors from a normal distribution with mean *µ*_*y*_ and variance 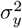. We sample consumption coefficients from a normal distribution with mean *µ*_*c*_*/M* and variance of 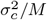, because resources are substitutable and thus high resource diversity leads to less consumption per resource. **(B)** The stability condition, derived using the cavity method. Note that throughout the manuscript, we focus on the effect of changing resource pool size (*M*) **not** the number of surviving resources (*M* ^∗^).

## Results

### A large resource pool size stabilises species-rich communities

The competitive exclusion principle states that increasing the diversity of available resources can support more species, which, according to May’s theory, should destabilise community dynamics (Fig. 1). We tested this hypothesis by simulating community dynamics across different resource pool sizes *M* (our measure of resource diversity). As expected, larger resource pool sizes supported greater species diversity (Fig. 3 A), but contrary to our prediction, these diverse communities were also more stable (Fig. 3 B and C). Communities with smaller resource pools, in contrast, exhibited persistent fluctuations characteristic of unstable dynamics (Fig. 3 D, compare top and bottom panels). These results demonstrate that resource-explicit dynamics can generate a positive relationship between species diversity and community stability, in stark contrast to the negative relationship from resource-implicit models.

**Figure 3:**
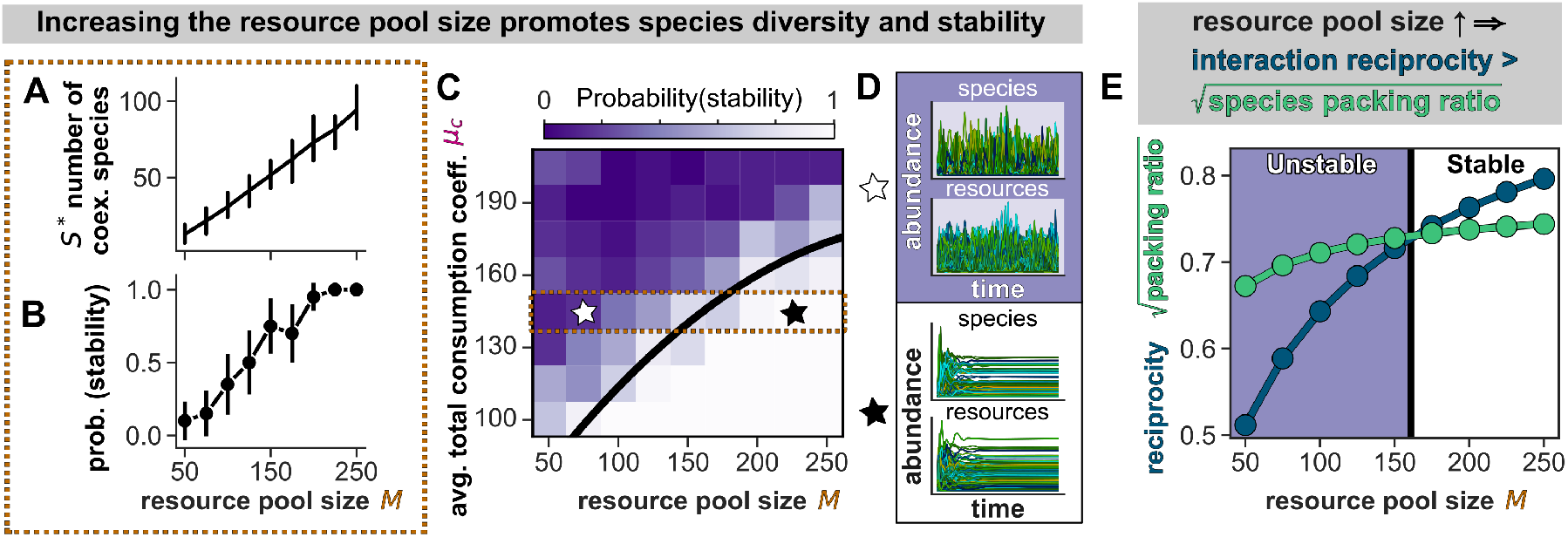
A large resource pool stabilises communities with diverse species. Increasing the resource pool size promotes species diversity **(A)** and community stability **(B)**, in direct opposition of our 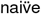 expectation in fig. 1 C. Shown are simulations for *µ*_*c*_ = 145, corresponding to the outlined area in panel (C). **(C)** Stability diagram as a function of resource pool size (*M*) and average total consumption coefficient *µ*_*c*_. Each cell shows results from 20 simulated communities, with darker shades indicating a higher proportion of simulations showing unstable, chaotic dynamics (with maximum Lyapunov exponent *>* 0). Purple: unstable region; white: stable region. The black line shows the analytically-derived stability boundary (SI Appendix C). **(D)** Representative examples of community dynamics from the unstable region with chaotic abundance fluctuations (white star, top) and from the stable region (black star, bottom). Each curve in each plot represents a single species or resource, as labeled. **(E)** Increasing the resource pool size stabilises communities by increasing the interaction reciprocity faster than 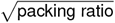. Shown are analytically-derived values of the reciprocity (dark blue), 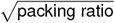 (green) and the stability boundary (black) for *µ*_*c*_ = 145.

### Large resource pools stabilise diverse communities by increasing interaction reciprocity faster than 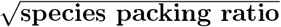

To understand how resource dynamics can invert the negative diversity-stability relationship predicted by resource-implicit models, we analytically derived our model’s stability condition using the cavity method (SI). This is a statistical physics approach that explains how community-level properties arise from ecological interactions [27, 52, 58, 59, 65–67]. Consistent with previous studies in resource-explicit models [52, 56, 57], we found that communities are stable when the correlation between growth and consumption exceeds the square root of the species packing ratio 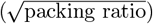 (Fig. 2 B).

High reciprocity between growth and consumption (≈ 1) indicates that consumers are “well-adapted” to their metabolic niches [57]: given they have a fixed total consumption rate, high reciprocity means they predominantly consume resources that promote rapid growth. In contrast, low reciprocity means consumers are less selective about resource quality, so a greater fraction of their consumption is spent on resources that do not promote growth. The packing ratio — the ratio of surviving species to surviving resources (Fig. 2 B) — quantifies how many species coexist per resource. Due to competitive exclusion, this ratio is generically less than 1, except in “supersaturated” communities with extremely fine-tuned consumption patterns [68, 69]. Therefore, we can interpret the stability condition as the following: reciprocity sets an upper bound on the number of species that can coexist on the same set of resources([57], SI Appendix F). For example, high reciprocity allows consumers to exploit shared resources in different ways (i.e., they occupy distinct niches), enabling them to densely pack into communities without encroaching on each other’s niches; therefore, consumers stably coexist. Otherwise, consumers can continuously encroach on each other’s niches, making community dynamics unstable.

Unlike previous studies, in our model both reciprocity and the species packing ratio emerge naturally from the mechanisms underpinning consumer-resource interactions rather than being imposed *a priori* [52, 56, 57]. This allowed us to determine how both reciprocity and packing ratio change with resource pool size (Fig. 2 B), as well as how other interaction parameters (e.g. the mean and variance in consumption and the yield coefficients) affect community stability (which we explore later). We found that small resource pools produced unstable communities because in this case, 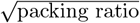 was much higher than reciprocity. Increasing the resource pool size raised both reciprocity and 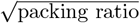, but reciprocity rose much faster, eventually leading to stability (Fig. 3 E, black transition). Although the exact transition point is not fixed (e.g. can depend on a consumer’s average consumption of all resources (*µ*_*c*_); Fig. 3C), the qualitative picture is robust: increasing resource pool size generates species-rich and stable communities by increasing reciprocity faster than 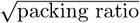, which prevents densely-packed consumers from encroaching on each other’s niches.

### Resource dynamics are critical for large resource pools to generate stable, species-rich communities

Are resource dynamics important in driving the observed positive diversity-stability relationships? To address this question, we simulated the effect of resource pool size on stability when resource dynamics are neglected - i.e. set to pseudo-steady state which occurs, for example, when resource dynamics are fast relative to consumer dynamics. Under this scenario, the consumer-resource model reduces to a model where species compete directly, which we call an “effective Lotka-Volterra model” (eLV) (Fig. 4 A). In eLV, species’ growth rates and competition coefficients are determined by their consumption and their competitors’ consumption of shared resources, and not by resource abundances (Fig. 4 B). Unlike in CRM where increasing resource diversity improved stability, in eLV stability is largely unaffected by resource diversity (Fig. 4 C). Indeed, eLV’s stability condition is fundamentally different to that of CRM across all resource pool sizes (SI Appendix E). Thus, resource dynamics themselves, not merely resource-mediated interactions, drive the positive relationship between resource diversity and stability.

**Figure 4:**
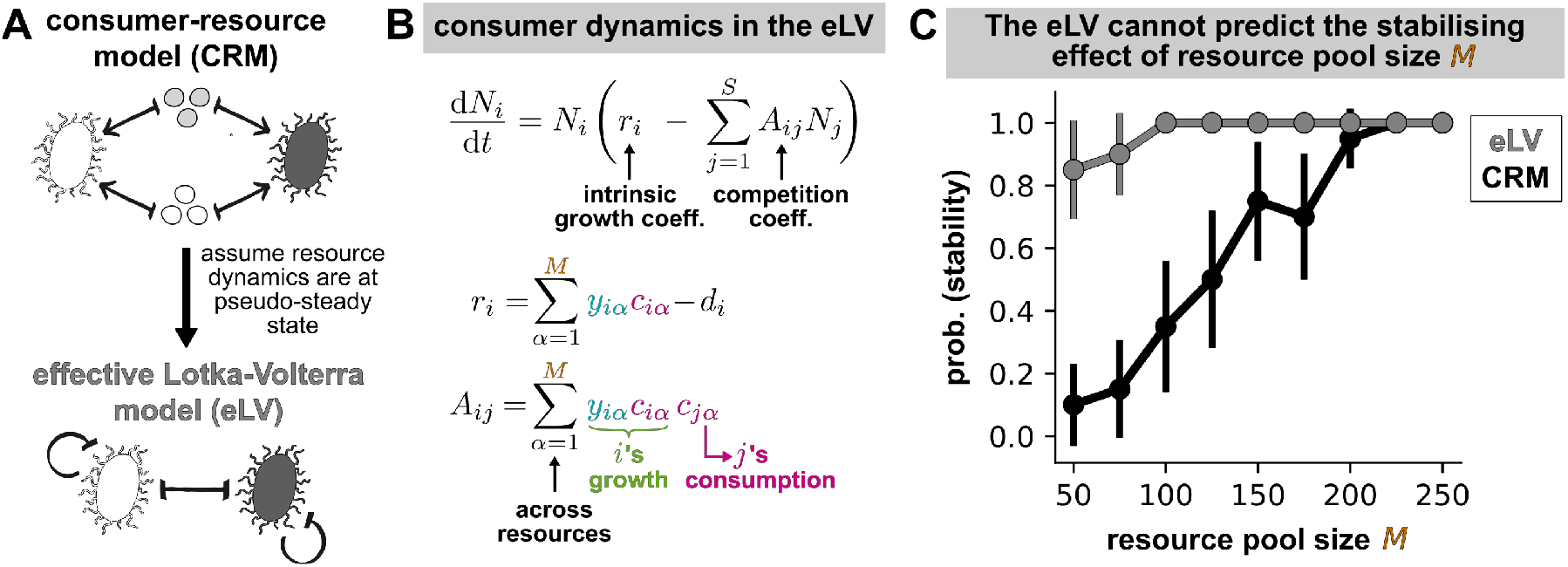
Resource-implicit models cannot capture the stabilising effect of resource diversity. **(A)** Assuming resource dynamics reach pseudo-steady state (e.g., under fast resource dynamics), the resource-explicit consumer-resource model (CRM) reduces to the resource-implicit effective Lotka-Volterra model (eLV). **(B)** In eLV, species competition is direct, or resource-implicit. However, the eLV is parametrised by CRM: for example, the competition coefficient of species *j* on *i* depends on *i*’s growth and *j*’s consumption on shared resources. We assume that all resources survive because extinctions cannot be determined without explicitly modelling resource dynamics; results are unchanged when only surviving resources are included (SI Appendix E). **(C)** Increasing resource pool size *M* stabilises community dynamics in CRM (black line), but has little effect in eLV (grey line). Shown are simulations for *µ*_*c*_ = 145. Growth and competition coefficients in eLV are parametrised from existing CRMs (SI Appendix E)

### Increasing heterogeneity in consumer-resource interactions through different biological mechanisms induces opposing stability transitions

Having shown the positive diversity-stability relationship, we tested whether increasing heterogeneity in interaction strength destabilises communities. This prediction comes from both resource-implicit [25–27, 29] and -explicit models [52]. Interaction heterogeneity can be achieved through increasing heterogeneity in underlying consumer-resource interactions, which is easily observable for the special case where consumer-resource model is reduced to eLV [50, 52, 62] (SI Appendix E).

Contrary to expectation, increasing the variance in consumption coefficient 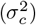 and the variance in the yield conversion factor 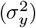 induced opposing stability transitions. While high variance in yield conversion destabilized communities (Fig. 5 C), high variance in consumption coefficient surprisingly stabilised community dynamics (Fig. 5 A).

**Figure 5:**
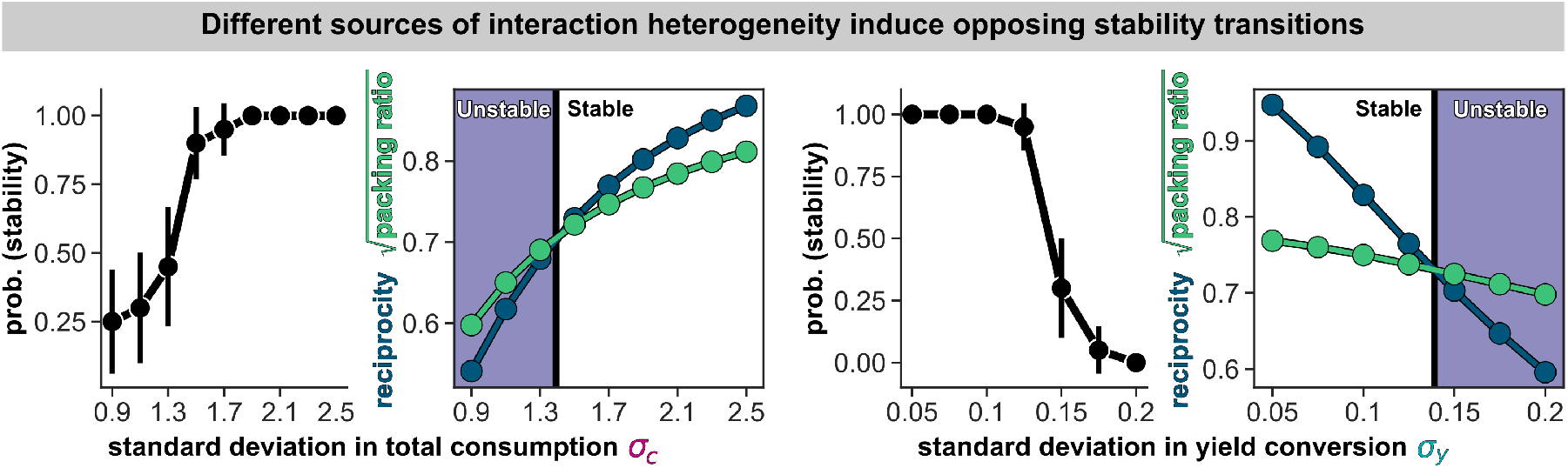
Different sources of interaction heterogeneity induce opposing stability transitions. **(A)** Increasing the heterogeneity in the total consumption coefficient stabilises community dynamics. **(B)** Increasing *σ*_*c*_ stabilises communities by increasing the growth-consumption correlation faster than the 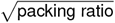. The numerically-solved values of the analytically-derived correlation (dark blue), packing ratio (green) and stability boundary (black). **(C)** Increasing the heterogeneity in the yield conversion factor destabilises community dynamics. **(D)** Increasing *σ*_*y*_ destabilises communities by decreasing the growth-consumption correlation faster than 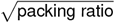. For each point, 20 communities were sampled from the same parameter distributions and simulated. The probability communities exhibited stable dynamics was determined by the proportion of communities with numerically-estimated maximum Lyapunov exponent less than 0. All parameters were the same as fig. 3, except *M* = 200 and *µ*_*c*_ = 160.

To understand why variance in consumption and yield conversion had opposing effects on stability, we again turned to our stability condition. At low consumption variance, 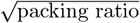 was larger than reciprocity, placing communities in the unstable region (Fig. 5 B). Increasing 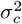 increased both reciprocity and 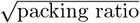, but reciprocity increased faster, which ultimately stabilised communities (Fig. 5 B). Increasing yield variance 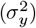 showed the opposite effect: When 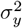 was low, reciprocity exceeded 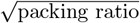, making communities stable. Increasing 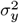 decreased both quantities, but reciprocity declined faster until it became smaller than 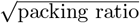, destabilising community dynamics (Fig. 5 D). Together, the *source* of heterogeneity in consumer-resource interactions — not the presence of variation alone — determines community stability.

## Discussion

Robert May’s stability condition[25, 26, 28] and the competitive exclusion principle [49–51] together predict that increasing resource diversity should destabilise communities (Fig. 1 C). We tested this prediction in a consumer-resource model where species growth and resource consumption are coupled through a yield conversion factor. Contrary to the expectation, large resource pools generated stable, species-rich communities (Fig.), and this stabilisation is driven by resource dynamics (Fig.). We also found that different sources of heterogeneity induced opposing stability transitions: While increasing variance in yield indeed destabilised community dynamics, increasing variance in consumption stabilised communities (Fig.). Overall, resource dynamics and the biological mechanisms coupling them to species dynamics can invert classical relationships between species diversity or interaction heterogeneity and stability.

Our results differ from May’s predictions because resource-implicit and explicit models have different stability criteria. In resource-implicit models, communities become unstable when the total strength of inter-species interactions exceeds self-inhibition. This occurs when species diversity and interaction heterogeneity are high [25–27, 29], when the average inter-species interaction strength is strong, and when interaction networks are highly connected [23–26, 29]. Although resource-explicit models can be reduced to resource-implicit models when, for example, resource dynamics are much faster than species dynamics and are thus “driven” by species dynamics [50, 62, 70–72], this equivalence does not hold in our work. When we assumed fast resource-dynamics in the consumer-resource model, the effective Lokta-Volterra model could not produce the stability transitions of the resource-explicit model (Fig. 4). Therefore, the positive diversity-stability relationship, as well as other stability transitions, were actively driven by resource dynamics. In this regime, communities become unstable when the correlation between growth and consumption is lower than 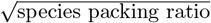, meaning that consumers encroach on each other’s niches [52, 56, 57]. Given that most ecological communities interact strongly with their resource environment, we expect that using resource-explicit models is essential for making meaningful predictions about community stability [48].

To our knowledge, the only other mechanism that generates positive diversity-stability relationships in the resource-implicit gLV model is when species exhibit sublinear growth, rather than logistic growth as is widely assumed [22]. However, as far as we are aware, sublinear growth has only been observed indirectly in ecological time series data, not directly in controlled experiments [73]. In addition, it lacks a clear mechanistic basis. For example, most resource-explicit models such as the MacArthur or Monod consumer-resource model are well-approximated by logistic [64] or superlinear, not sublinear, growth [74]. In contrast, our model is more biologically grounded. For example, consumed resources are converted into consumer biomass through yield conversion factors, a simple yet natural representation of consumer-specific metabolism. It will be worthwhile to experimentally test our diversity-stability relationship: by increasing the resource pool size, one can test whether the frequency of observing stable, species-rich communities increases.

Our study demonstrates that resource diversity is a biologically grounded mechanism for generating stable, diverse communities. But how broadly does this mechanism apply to natural ecosystems? One limitation of our study is that we only model competitive unstructured interactions, excluding the functional groups and cross-feeding interactions common in natural and experimental communities [36, 40, 75]. However, we expect that including these features would maintain or even strengthen the diversity-stability relationship. Adding functional groups to our model would maintain the relationship within groups (as our model effectively has one functional group) but decrease niche encroachment between groups, potentially accelerating the onset of stability. Cross-feeding expands the effective resource pool as species diversity increases, which our results suggest should stabilise communities. In the future, it will be useful to understand more broadly how the plethora of resource-mediated community interactions impact the stability of complex ecosystems.

## Acknowledgements

J.RC. and W.S. acknowledge support from the Royal Society Wolfson Fellowship. J.RC. would also like to thank the Shou group lab members for their feedback on the project. W.S. is additionally supported by an Academy of Medical Sciences Professorship. A.G. acknowledges support from the Ashok and Gita Vaish Junior Researcher Award, the DAE, Govt. of India under project number RTI4001, as well as the Ramanujan Fellowship. Finally, we thank National Institute for Theory and Mathematics in Biology (NITMB, USA) where we presented this work and received valuable feedback.

## Notes

https://github.com/JamilaRowlandChandler/CRM-Resource-diversity-vs-Stability

### Competing Interest Statement

The authors have declared no competing interest.

### Summary of Updates

New results section on how resource dynamics drive the resource diversity-stability relationship (the relationship is lost in a resource-mediated by implicit model). More details added to the SI and extended information. Slight revision to methods and figures.

## References

[1] Jeremiah J. Faith et al. “The Long-Term Stability of the Human Gut Microbiota”. en. In: Science 341.6141 (July 2013), p. 1237439. doi: 10.1126/science.1237439.

[2] Catherine A. Lozupone et al. “Diversity, stability and resilience of the human gut microbiota”. en. In: Nature 489.7415 (Sept. 2012), pp. 220–230. doi: 10.1038/nature11550.

[3] Didier Gonze et al. “Multi-stability and the origin of microbial community types”. en. In: The ISME Journal 11.10 (Oct. 2017), pp. 2159–2166. doi: 10.1038/ismej.2017.60.

[4] Tahmineh Khazaei et al. “Metabolic multistability and hysteresis in a model aerobe-anaerobe microbiome community”. en. In: Science Advances 6.33 (Aug. 2020), eaba0353. doi: 10.1126/sciadv.aba0353.

[5] William Lopes, Daniel R. Amor, and Jeff Gore. “Cooperative growth in microbial communities is a driver of multistability”. en. In: Nature Communications 15.1 (June 2024), p. 4709. doi: 10.1038/s41467-024-48521-9.

[6] Ana Fernandez et al. “How Stable Is Stable? Function versus Community Composition”. en. In: Applied and Environmental Microbiology 65.8 (Aug. 1999), pp. 3697–3704. doi: 10.1128/AEM.65.8.3697-3704.1999.

[7] Nuria Fernandez-Gonzalez, Julie A. Huber, and Joseph J. Vallino. “Microbial Communities Are Well Adapted to Disturbances in Energy Input”. en. In: mSystems 1.5 (Oct. 2016). Ed. by Haiyan Chu, e00117–16. doi: 10.1128/mSystems.00117-16.

[8] Jiliang Hu et al. “Emergent phases of ecological diversity and dynamics mapped in microcosms”. en. In: Science 378.6615 (Oct. 2022), pp. 85–89. doi: 10.1126/science.abm7841.

[9] Christoph Ratzke, Julien Barrere, and Jeff Gore. “Strength of species interactions determines biodiversity and stability in microbial communities”. en. In: Nature Ecology & Evolution 4.3 (Feb. 2020), pp. 376–383. doi: 10.1038/s41559-020-1099-4.

[10] Antonio M. Martin-Platero et al. “High resolution time series reveals cohesive but short-lived communities in coastal plankton”. en. In: Nature Communications 9.1 (Jan. 2018), p. 266. doi: 10.1038/s41467-017-02571-4.

[11] Elisa Beninca et al. “Chaos in a long-term experiment with a plankton community”. en. In: Nature 451.7180 (Feb. 2008), pp. 822–825. doi: 10.1038/nature06512.

[12] Elisa Beninca et al. “Species fluctuations sustained by a cyclic succession at the edge of chaos”. en. In: Proceedings of the National Academy of Sciences 112.20 (May 2015), pp. 6389–6394. doi: 10.1073/pnas.1421968112.

[13] JuÂ Young Chang et al. “Decreased Diversity of the Fecal Microbiome in Recurrent Clostridium difficile âAssociated Diarrhea”. en. In: The Journal of Infectious Diseases 197.3 (Feb. 2008), pp. 435–438. doi: 10.1086/525047.

[14] Jonas Halfvarson et al. “Dynamics of the human gut microbiome in inflammatory bowel disease”. en. In: Nature Microbiology 2.5 (Feb. 2017), p. 17004. doi: 10.1038/nmicrobiol.2017.4.

[15] H.W. Paerl and J.L. Pinckney. “A mini-review of microbial consortia: Their roles in aquatic production and biogeochemical cycling”. en. In: Microbial Ecology 31.3 (1996). Publisher: Springer Science and Business Media LLC. doi: 10.1007/bf00171569.

[16] Judith Prommer et al. “Increased microbial growth, biomass, and turnover drive soil organic carbon accumulation at higher plant diversity”. en. In: Global Change Biology 26.2 (Feb. 2020), pp. 669–681. doi: 10.1111/gcb.14777.

[17] Xujun Liu et al. “Plant diversity and species turnover co-regulate soil nitrogen and phosphorus availability in Dinghushan forests, southern China”. en. In: Plant and Soil 464. 1-2 (July 2021), pp. 257–272. doi: 10.1007/s11104-021-04940-x.

[18] Tim A. Hoek et al. “Resource Availability Modulates the Cooperative and Competitive Nature of a Microbial Cross-Feeding Mutualism”. en. In: PLOS Biology 14.8 (Aug. 2016). Ed. by Nathalie Balaban, e1002540. doi: 10.1371/journal.pbio.1002540.

[19] Wenying Shou, Sri Ram, and Jose M. G. Vilar. “Synthetic cooperation in engineered yeast populations”. en. In: Proceedings of the National Academy of Sciences 104.6 (Feb. 2007), pp. 1877–1882. doi: 10.1073/pnas.0610575104.

[20] Cyrille Violle et al. “Phylogenetic limiting similarity and competitive exclusion: Phylogenetic relatedness and competition”. en. In: Ecology Letters 14.8 (Aug. 2011), pp. 782–787. doi: 10.1111/j.1461-0248.2011.01644.x.

[21] Tyler D. Ross, Christopher A. Klausmeier, and Ophelia S. Venturelli. Metabolic interplay drives population cycles in a cross-feeding microbial community. en. Oct. 2024. doi: 10.1101/2024.10.14.618235.

[22] Ian A. Hatton et al. “Diversity begets stability: Sublinear growth and competitive coexistence across ecosystems”. en. In: Science 383.6688 (Mar. 2024), eadg8488. doi: 10.1126/science.adg8488.

[23] Jacopo Grilli, Tim Rogers, and Stefano Allesina. “Modularity and stability in ecological communities”. en. In: Nature Communications 7.1 (June 2016), p. 12031. doi: 10.1038/ncomms12031.

[24] Stefano Allesina et al. “Predicting the stability of large structured food webs”. en. In: Nature Communications 6.1 (July 2015), p. 7842. doi: 10.1038/ncomms8842.

[25] Robert M. May. “Will a Large Complex System be Stable?” en. In: Nature 238.5364 (Aug. 1972), pp. 413–414. doi: 10.1038/238413a0.

[26] Robert M. May. “Qualitative Stability in Model Ecosystems”. en. In: Ecology 54.3 (May 1973), pp. 638–641. doi: 10.2307/1935352.

[27] Guy Bunin. “Ecological communities with Lotka-Volterra dynamics”. en. In: Physical Review E 95.4 (Apr. 2017), p. 042414. doi: 10.1103/PhysRevE.95.042414.

[28] Stefano Allesina and Si Tang. “Stability criteria for complex ecosystems”. en. In: Nature 483.7388 (Mar. 2012), pp. 205–208. doi: 10.1038/nature10832.

[29] Emil Mallmin, Arne Traulsen, and Silvia De Monte. “Chaotic turnover of rare and abundant species in a strongly interacting model community”. en. In: Proceedings of the National Academy of Sciences 121.11 (Mar. 2024), e2312822121. doi: 10.1073/pnas.2312822121.

[30] David Tilman, Peter B. Reich, and Johannes M. H. Knops. “Biodiversity and ecosystem stability in a decade-long grassland experiment”. en. In: Nature 441.7093 (June 2006), pp. 629–632. doi: 10.1038/nature04742.

[31] Cameron Wagg et al. “Biodiversityâstability relationships strengthen over time in a long-term grassland experiment”. en. In: Nature Communications 13.1 (Dec. 2022), p. 7752. doi: 10.1038/s41467-022-35189-2.

[32] Amy L. Downing, Bryan L. Brown, and Mathew A. Leibold. “Multiple diversityâstability mechanisms enhance population and community stability in aquatic food webs”. en. In: Ecology 95.1 (Jan. 2014), pp. 173–184. doi: 10.1890/12-1406.1.

[33] Jasper Van Ruijven and Frank Berendse. “Contrasting effects of diversity on the temporal stability of plant populations”. en. In: Oikos 116.8 (Aug. 2007), pp. 1323–1330. doi: 10.1111/j.0030-1299.2007.16005.x.

[34] David Tilman and John A. Downing. “Biodiversity and stability in grasslands”. en. In: Nature 367.6461 (Jan. 1994), pp. 363–365. doi: 10.1038/367363a0.

[35] Lori Niehaus et al. “Microbial coexistence through chemical-mediated interactions”. en. In: Nature Communications 10.1 (May 2019), p. 2052. doi: 10.1038/s41467-019-10062-x.

[36] Elizabeth J. Culp and Andrew L. Goodman. “Cross-feeding in the gut microbiome: Ecology and mechanisms”. en. In: Cell Host & Microbe 31.4 (Apr. 2023). Publisher: Elsevier BV, pp. 485–499. doi: 10.1016/j.chom.2023.03.016.

[37] Martina Dal Bello et al. “Resourceâdiversity relationships in bacterial communities reflect the network structure of microbial metabolism”. en. In: Nature Ecology & Evolution 5.10 (Aug. 2021), pp. 1424–1434. doi: 10.1038/s41559-021-01535-8.

[38] Sylvie Estrela et al. “Nutrient dominance governs the assembly of microbial communities in mixed nutrient environments”. en. In: eLife 10 (Apr. 2021), e65948. doi: 10.7554/eLife.65948.

[39] Harry J. Flint et al. “Interactions and competition within the microbial community of the human colon: links between diet and health”. en. In: Environmental Microbiology 9.5 (May 2007). Publisher: Wiley, pp. 1101–1111. doi: 10.1111/j.1462-2920.2007.01281.x.

[40] Joshua E. Goldford et al. “Emergent simplicity in microbial community assembly”. en. In: Science 361.6401 (Aug. 2018), pp. 469–474. doi: 10.1126/science.aat1168.

[41] Diane Lawrence et al. “Species Interactions Alter Evolutionary Responses to a Novel Environment”. en. In: PLoS Biology 10.5 (May 2012). Ed. by Stephen P. Ellner, e1001330. doi: 10.1371/journal.pbio.1001330.

[42] Mark E. Ritchie and David Tilman. “Predictions of species interactions from consumer-resource theory: experimental tests with grasshoppers and plants”. en. In: Oecologia 94.4 (July 1993), pp. 516–527. doi: 10.1007/BF00566967.

[43] Xin Sun et al. “Metabolic Plasticity Shapes Microbial Communities across a Temperature Gradient”. en. In: The American Naturalist 204.4 (Oct. 2024), pp. 381–399. doi: 10.1086/731997.

[44] Jason M Tylianakis et al. “Resource Heterogeneity Moderates the Biodiversity-Function Relationship in Real World Ecosystems”. en. In: PLoS Biology 6.5 (May 2008). Ed. by Michel Loreau, e122. doi: 10.1371/journal.pbio.0060122.

[45] Jeremy P. H. Wong et al. “Fluid flow structures gut microbiota biofilm communities by distributing public goods”. en. In: Proceedings of the National Academy of Sciences 120.25 (June 2023), e2217577120. doi: 10.1073/pnas.2217577120.

[46] Hyunseok Lee, Blox Bloxham, and Jeff Gore. “Resource competition can explain simplicity in microbial community assembly”. en. In: Proceedings of the National Academy of Sciences 120.35 (Aug. 2023), e2212113120. doi: 10.1073/pnas.2212113120.

[47] Michael Daniels, Simon Van Vliet, and Martin Ackermann. “Changes in interactions over ecological time scales influence single-cell growth dynamics in a metabolically coupled marine microbial community”. en. In: The ISME Journal 17.3 (Mar. 2023), pp. 406–416. doi: 10.1038/s41396-022-01312-w.

[48] Babak Momeni, Li Xie, and Wenying Shou. “Lotka-Volterra pairwise modeling fails to capture diverse pairwise microbial interactions”. en. In: eLife 6 (Mar. 2017), e25051. doi: 10.7554/eLife.25051.

[49] Robert MacArthur and Richard Levins. “COMPETITION, HABITAT SELECTION, AND CHARACTER DISPLACEMENT IN A PATCHY ENVIRONMENT”. en. In: Proceedings of the National Academy of Sciences 51.6 (June 1964), pp. 1207–1210. doi: 10.1073/pnas.51.6.1207.

[50] Robert MacArthur. “SPECIES PACKING, AND WHAT COMPETITION MINIMIZES”. en. In: Proceedings of the National Academy of Sciences 64.4 (Dec. 1969), pp. 1369–1371. doi: 10.1073/pnas.64.4.1369.

[51] David Tilman. Resource Competition and Community Structure. Princeton University Press, 1982. doi: 10.2307/j.ctvx5wb72.

[52] Emmy Blumenthal, Jason W. Rocks, and Pankaj Mehta. “Phase Transition to Chaos in Complex Ecosystems with Nonreciprocal Species-Resource Interactions”. en. In: Physical Review Letters 132.12 (Mar. 2024), p. 127401. doi: 10.1103/PhysRevLett.132.127401.

[53] Vincent A. A. Jansen and Giorgos D. Kokkoris. “Complexity and stability revisited”. en. In: Ecology Letters 6.6 (June 2003), pp. 498–502. doi: 10.1046/j.1461-0248.2003.00464.x.

[54] Lyle Poley, Joseph W. Baron, and Tobias Galla. “Generalized Lotka-Volterra model with hierarchical interactions”. en. In: Physical Review E 107.2 (Feb. 2023), p. 024313. doi: 10.1103/PhysRevE.107.024313.

[55] Laura Sidhom and Tobias Galla. “Ecological communities from random generalised Lotka-Volterra dynamics with non-linear feedback”. In: (2019). Publisher: arXiv Version Number: 2. doi: 10.48550/ARXIV.1909.05802.

[56] Yizhou Liu, Jiliang Hu, and Jeff Gore. Ecosystem stability relies on diversity difference between trophic levels. en. Aug. 2024. doi: 10.1101/2024.08.23.609466.

[57] Yizhou Liu et al. “Complex Ecosystems Lose Stability When Resource Consumption Is Out of Niche”. en. In: Physical Review X 15.1 (Jan. 2025), p. 011003. doi: 10.1103/PhysRevX.15.011003.

[58] Madhu Advani, Guy Bunin, and Pankaj Mehta. “Statistical physics of community ecology: a cavity solution to MacArthur’s consumer resource model”. eng. In: Journal of Statistical Mechanics (Online) 2018 (Mar. 2018), p. 033406. doi: 10.1088/1742-5468/aab04e.

[59] Wenping Cui, Robert Marsland, and Pankaj Mehta. “Diverse communities behave like typical random ecosystems”. en. In: Physical Review E 104.3 (Sept. 2021), p. 034416. doi: 10.1103/PhysRevE.104.034416.

[60] Imane Akjouj et al. “Complex systems in ecology: a guided tour with large LotkaâVolterra models and random matrices”. en. In: Proceedings of the Royal Society A: Mathematical, Physical and Engineering Sciences 480.2285 (Mar. 2024), p. 20230284. doi: 10.1098/rspa.2023.0284.

[61] Thibaut Arnoulx De Pirey and Guy Bunin. “Many-Species Ecological Fluctuations as a Jump Process from the Brink of Extinction”. en. In: Physical Review X 14.1 (Mar. 2024), p. 011037. doi: 10.1103/PhysRevX.14.011037.

[62] Itay Dalmedigos and Guy Bunin. “Dynamical persistence in high-diversity resource-consumer communities”. en. In: PLOS Computational Biology 16.10 (Oct. 2020). Ed. by James OâDwyer. Publisher: Public Library of Science (PLoS), e1008189. doi: 10.1371/journal.pcbi.1008189.

[63] F Roy et al. “Numerical implementation of dynamical mean field theory for disordered systems: application to the LotkaâVolterra model of ecosystems”. In: Journal of Physics A: Mathematical and Theoretical 52.48 (Nov. 2019), p. 484001. doi: 10.1088/1751-8121/ab1f32.

[64] Akshit Goyal, Jason W. Rocks, and Pankaj Mehta. “Universal Niche Geometry Governs the Response of Ecosystems to Environmental Perturbations”. en. In: PRX Life 3.1 (Feb. 2025), p. 013010. doi: 10.1103/PRXLife.3.013010.

[65] Wenping Cui, Robert Marsland, and Pankaj Mehta. “Effect of Resource Dynamics on Species Packing in Diverse Ecosystems”. en. In: Physical Review Letters 125.4 (July 2020), p. 048101. doi: 10.1103/PhysRevLett.125.048101.

[66] Pankaj Mehta et al. “Constrained optimization as ecological dynamics with applications to random quadratic programming in high dimensions”. en. In: Physical Review E 99.5 (May 2019), p. 052111. doi: 10.1103/PhysRevE.99.052111.

[67] Matthieu Barbier and Jean-Francois Arnoldi. The cavity method for community ecology. en. June 2017. doi: 10.1101/147728.

[68] Anna Posfai, Thibaud Taillefumier, and Ned S. Wingreen. “Metabolic Trade-Offs Promote Diversity in a Model Ecosystem”. en. In: Physical Review Letters 118.2 (Jan. 2017), p. 028103. doi: 10.1103/PhysRevLett.118.028103.

[69] Jef Huisman and Franz J. Weissing. “Biodiversity of plankton by species oscillations and chaos”. en. In: Nature 402.6760 (Nov. 1999), pp. 407–410. doi: 10.1038/46540.

[70] Peter Chesson. “MacArthur’s consumer-resource model”. en. In: Theoretical Population Biology 37.1 (Feb. 1990), pp. 26–38. doi: 10.1016/0040-5809(90)90025-Q.

[71] Robert Marsland, Wenping Cui, and Pankaj Mehta. “A minimal model for microbial biodiversity can reproduce experimentally observed ecological patterns”. en. In: Scientific Reports 10.1 (Feb. 2020), p. 3308. doi: 10.1038/s41598-020-60130-2.

[72] Stefan Vet et al. “Bistability in a system of two species interacting through mutualism as well as competition: Chemostat vs. Lotka-Volterra equations”. en. In: PLOS ONE 13.6 (June 2018). Ed. by Ramon Grima, e0197462. doi: 10.1371/journal.pone.0197462.

[73] Samuel F. M. Hart et al. “Highâthroughput quantification of microbial birth and death dynamics using fluorescence microscopy”. en. In: Quantitative Biology 7.1 (Mar. 2019), pp. 69–81. doi: 10.1007/s40484-018-0160-7.

[74] Kevin S. McCann and Gabriel Gellner, eds. Theoretical Ecology: concepts and applications. en. 1st ed. Oxford University Press, May 2020. doi: 10.1093/oso/9780198824282.001.0001.

[75] Sylvie Estrela et al. Diversity begets diversity under microbial niche construction. en. Feb. 2022. doi: 10.1101/2022.02.13.480281.

