## Supplementary material for "Resource diversity begets stability in complex ecosystems": SI

### Supplementary information for "Resource diversity begets stability in complex ecosystems"

### A Nomenclature

This table details the nomenclature used throughout the supplementary information

| Symbol | Description |
|---|---|
| **Model variables and parameters** | |
| $N_i$ | abundance of species $i$ |
| $R_\alpha$ | abundance of resource $\alpha$ |
| $c_{i\alpha}$ | consumption coefficient of resource $\alpha$ by species $i$ |
| $y_{i\alpha}$ | yield conversion factor — growth rate yield for species $i$ per unit consumption flux of resource $\alpha$ |
| $g_{i\alpha}$ | growth coefficient of species $i$ on resource $\alpha$; $g_{i\alpha} = y_{i\alpha} c_{i\alpha}$ |
| $d_i$ | death rate of species $i$ (in the absence of resources) |
| $b_\alpha$ | supply rate of resource $\alpha$ from a consumer perspective; also the intrinsic growth rate of resource $\alpha$ from a resource perspective |
| $K_\alpha$ | carrying capacity of resource $\alpha$ |
| **Statistical properties** | |
| $\mu_c$ | the mean consumption coefficient $c_{i\alpha}$ |
| $\sigma_c$ | the standard deviation of consumption coefficients $c_{i\alpha}$ |
| $\mu_y$ | the mean yield conversion factor $c_{i\alpha}$ |
| $\sigma_y$ | the standard deviation of the yield conversion factor $c_{i\alpha}$ |
| $\rho_{c,g}$ | the correlation between consumption and growth coefficients of a given species, or the interaction reciprocity; given by $\mathrm{Corr}(c_{i\alpha}, g_{i\alpha})$ |
| $z_x$ or $Z_x$ | a standard normal variable with mean 0 and variance 1 used to sample the variable $x$ (subscripts may change based on variable) |
| $\langle ... \rangle$ | the expected value, or average |
| **Self-consistency equations and other terms relevant to the cavity method** | |
| $\mu_{g_N}$ and $\mu_{g_R}$ | the mean net growth rate of a species and resource, respectively |
| $\sigma_{g_N}$ and $\sigma_{g_R}$ | the standard deviation in the net growth rate of a species and resource, respectively |
| $\triangle\, g_N$ and $\triangle\, g_R$ | the inverse coefficient of variation of the net growth rate of species and resources, respectively. $\triangle\, g_N = \mu_{g_N}/\sigma_{g_N}$, $\triangle\, g_R = \mu_{g_R}/\sigma_{g_R}$ |
| $\tilde{N}_i$ and $\tilde{R}_\alpha$ | The perturbed abundances of species $i$ and resource $\alpha$ after the cavity species and resource invade the community |
| $N_0$ and $R_0$ | The steady-state abundance of the cavity species and the cavity resource |
| $S^*$ and $M^*$ | The number of surviving/coexisting species and resources, respectively |
| $\phi_N$ and $\phi_R$ | the species survival probability and the resource survival probability, respectively |
| $\langle N \rangle$ and $\langle R \rangle$ | the average species abundance and average resource abundance, respectively |
| $\langle N^2 \rangle$ and $\langle R^2 \rangle$ | the second moment in the species abundance distribution and the resource abundance distribution, respectively |

| | |
|---|---|
| $v_{ij}^{(N)}$ and $v_{\alpha j}^{(R)}$ | the respective susceptibilities of the abundance of species $i$ and resource $\alpha$ to small perturbations in the death rate of species $j$, respectively. $v_{ij}^{(N)} = \partial N_i/\partial d_j$, $v_{N,\alpha j} = \partial R_\alpha/\partial d_j$ |
| $v^{(N)}$ | the average trace of the susceptibility matrix $v_{ij}^{(N)}$, given by $\langle \partial N_i/\partial d_i \rangle$ |
| $\chi_{i\beta}^{(N)}$ and $\chi_{\alpha\beta}^{(R)}$ | the respective susceptibilities of the abundance of species $i$ and resource $\alpha$ to small perturbations in the supply rate (or intrinsic growth rate) of resource $\beta$. $\chi_{i\beta}^{(N)} = \partial N_i/\partial b_\beta$, $\chi_{\alpha\beta}^{(R)} = \partial R_\alpha/\partial b_\beta$ |
| $\chi^{(R)}$ | the average trace of the susceptibility matrix $\chi_{\alpha\beta}$ given by $\langle \partial R_\alpha/\partial b_\alpha \rangle$ |
| **Other** | |
| $R_\alpha^{(ss)}$ | the pseudo-steady state abundance of resource $\alpha$ under fast resource dynamics |

### B Model setup and parametrisation

Throughout this paper we study a biologically-motivated variant of the classic MacArthur consumer-resource model, adapted for a large number of species and resources. We follow the dynamics of the abundances of $S \gg 1$ species (consumers) growing by collectively consuming $M \gg 1$ resources. We follow the variables $N_i$ (the abundance of species $i$) and $R_\alpha$ (the abundance of resource $\alpha$), whose dynamics are governed by the following equations:

$$\begin{aligned} \frac{dN_i}{dt} =& N_i \left( \sum_{\alpha=1}^{M} y_{i\alpha} c_{i\alpha} R_\alpha - d_i \right), \\ \frac{dR_\alpha}{dt} =& R_\alpha \left( b_\alpha - \frac{R_\alpha}{K_\alpha} - \sum_{i=1}^{S} c_{i\alpha} N_i \right). \end{aligned} \tag{1}$$

where

- $c_{i\alpha}$ is the consumption or uptake coefficient of resource $\alpha$ by species $i$ (per capita uptake rate per unit biomass per unit resource abundance).
- $y_{i\alpha}$ is the yield or conversion efficiency of consumed resource $\alpha$ by species $i$ to its per capita growth rate
- $d_i$ is the per capita death rate of species $i$ (in the absence of resources) or maintenance rate of species $i$.
- $b_\alpha$ is the intrinsic supply rate of resource $\alpha$.
- $K_\alpha$ is the carrying capacity of resource $\alpha$
- $M$ is the resource pool size or the number of available resources, $S$ is the species pool size or the number of starting species.
- $\gamma$ is the ratio of resource pool size to species pool size, i.e., $\gamma = M/S$. Usually we will use $\gamma^{-1} = S/M$ in the calculations in this document.

In this model, species grow by consuming resources, with growth rates mediated not just by consumption but also by a yield conversion factor. Each species' net growth rate depends on the total flux of resources it consumes, its yields on those resources, and its intrinsic death rate or maintenance cost. Species reach a steady-state abundance and survive if after a long time, their net growth rate from consuming resources balances their death rate. Species that cannot achieve this balance continue to die and eventually go extinct at steady state. As species consume resources, resources get depleted. Resources reach steady state when their net supply rate balances total consumption by other species. We assume that resources in this model are self-renewing, which can be seen by noting that their dynamics follow logistic growth. Thus our model has the interpretation of a two-layer trophic ecosystem with resources serving as the bottom trophic layer of other biological organisms (e.g., grasses or phytoplankton) while consumers (what we often call "species") serving as the top trophic layer. Thus if a resource is depleted rapidly by other species and cannot be replenished or supplied fast enough, it might eventually go extinct at steady-state.

#### B.1 Parametrisation

Since we model complex ecosystems with a large number of species $S$ and resources $M$, we follow in a long tradition of theoretical ecology by assuming that model parameters are random variables drawn from appropriate distributions (e.g., [1–9]). We then study the general statistical properties of communities assembled under this parametrisation. For concreteness, in our simulations, we sample all parameters from Gaussian distributions, but note that our results are "universal" and apply when parameters are sampled from *any* distribution, as long as they have finite mean and variance and all parameters are sampled independently. This allows us to write all the model parameters in terms of a small number of underlying distributional parameters quantifying their means and variances. We write them as

$$y_{i\alpha} = \mu_y + \sigma_y z_{y,i\alpha}, \tag{2}$$

$$c_{i\alpha} = \frac{\mu_c}{M} + \frac{\sigma_c}{\sqrt{M}} z_{c,i\alpha}, \tag{3}$$

$$b_\alpha = \mu_b + \sigma_b z_{b,\alpha}, \tag{4}$$

$$d_i = \mu_d + \sigma_d z_{d,i}, \tag{5}$$

$$K_\alpha = \mu_K + \sigma_K z_{k,\alpha}, \tag{6}$$

where $z_{y,i\alpha}$, $z_{c,i\alpha}, z_{b,\alpha}, z_{d,i}$ and $z_{k,\alpha}$ are standard uncorrelated random variables (with a mean of 0 and a variance of 1). Standard variables obey the following rules:

$$\langle z_x \rangle = 0, \quad \langle z_x^2 \rangle = 1, \quad \langle z_x^n z_{y \neq x}^n \rangle = \langle z_x^n \rangle \langle z_{y \neq x}^n \rangle. \tag{7}$$

For simplicity we might assume that they are standard normal variables.

Throughout the paper, we will assume (without loss of generality) that $\mu_K = 1$ and $\sigma_K = 0$, resulting in

$$K_\alpha = 1. \tag{8}$$

Note that this is done only for minor simplicity and does not affect our results.

##### B.1.1 Useful mathematical properties based on parametrisation

For reference, below we list some useful mathematical properties that follow from our parametrisation above. Note that we denote the growth coefficient of consumer $i$ on resource $\alpha$ as $g_{i\alpha} = y_{i\alpha} c_{i\alpha}$.

The mean and variance of the consumption and growth coefficients are

| | **Consumption** ($c_{i\alpha}$) | **Growth** ($g_{i\alpha}$) | (9) |
|---|---|---|---|
| **Mean:** | $\frac{\mu_c}{M}$ | $\frac{\mu_y \mu_c}{M}$ | |
| **Variance:** | $\frac{\sigma_c^2}{M}$ | $\frac{(\mu_c \sigma_y)^2}{M^2} + \frac{\sigma_c^2(\mu_y^2 + \sigma_y^2)}{M}$ | (10) |

It is also useful to compute the correlation between per-species per-resource consumption coefficient ($c_{i,\alpha}$) and growth coefficient $g_{i\alpha}$, or the consumer-resource interaction reciprocity. This term is denoted by $\rho_{c,g}$. As we will show, this correlation is a critical determinant of the stability of community dynamics (Eq. 121).

**Growth-consumption correlation/interaction reciprocity:**

$$\rho_{c,g} = \frac{\mathrm{Cov}(c_{i\alpha}, g_{i\alpha})}{\sqrt{\mathrm{Var}(c_{i\alpha}) \cdot \mathrm{Var}(g_{i\alpha})}} = \frac{1}{\sqrt{1 + \frac{\sigma_y^2}{\mu_y^2}\left(1 + \frac{\mu_c^2}{M\sigma_c^2}\right)}} \tag{11}$$

Because of how we sample our model parameters, this is the only way growth and consumption rates can be correlated. For example, the growth of species $i$ on resource $\alpha$ is uncorrelated with its growth or consumption on resource $\beta$.

#### B.2 Model assumptions

Throughout the text, we make the following assumptions about the model that we collectively report below:

- The species and resource pool are large but finite ($1 \ll M \ll \infty$). (In statistical physics jargon, we do not take the thermodynamic limit $M \to \infty$.)
- All species and resources are statistically identical. As a consequence, consumption coefficients $c_{i\alpha}$ and yield conversion factors $y_{i\alpha}$ are each drawn independent of the species identity $i$ or resource identity $\alpha$.
- Resources are substitutable (i.e., they are the same type of resource, e.g., they are all carbon sources). This is why a species' total growth rate involves the sum of its growth rates over all individual resources, i.e., $\sum_{\alpha=1}^{M} g_{i\alpha} R_\alpha$.
- A species' combined resource consumption rate is largely independent of the resource pool size, $M$. Thus as a species consumes a greater number of resources $M$, its per-resource consumption coefficient $c_{i\alpha}$ decreases proportionately with $M$, i.e., $c_{i\alpha} \propto M$. Biologically this is reasonable since a cell or organism's total resource uptake rate cannot increase arbitrarily with the number of resources it consumes. The total consumption rate of a bacterial cell, for instance, depends on the coverage of transporters on its surface. It is thus reasonable to expect this quantity to saturate at a large number of resources $M \gg 1$.
- We will focus on community steady states which are feasible (contain species and resource abundances which are non-negative) and uninvadable (no extinct species should be able to invade the steady-state community, i.e., extinct species should have negative invasion growth rates).

### C Using the cavity method to determine community diversity, structure, and stability

In this study, we investigate how properties of the resource pool and species pool (such as the resource pool size $M$, variance in consumption coefficients $\sigma_c^2$ and yield conversion factors $\sigma_y^2$) affect community stability. To address this question, we first need to determine how the species and resource pool determine the properties of assembled communities, such as the species and resource survival probability ($\phi_N$ and $\phi_R$), average abundances ($\langle N \rangle$ and $\langle R \rangle$), and fluctuations in abundances ($\langle N^2 \rangle$ and $\langle R^2 \rangle$). We can then analyse how sensitive the steady states of these communities are to small perturbations, which allows us to derive the model's stability condition.

To do this, we use the cavity method, a statistical physics approach that describes the distribution of species and resource abundances in terms of the statistical properties of their underlying parameters. It allows us to determine the statistical properties of typical steady-state communities after community assembly, as well as assess their stability. This is in contrast with classical random matrix approaches, such as those of May [1, 2], which assess the stability of a prescribed steady-state community. These approaches typically neglect computing the properties of typical assembled communities, and instead prescribe them *ad hoc*, e.g., by assuming that all species coexist at a given set of abundances. Thus, the explicit advantage of the cavity method is that the stability condition we derive using it is much more generic and expected to apply more broadly to randomly assembled communities.

#### C.1 Steps of the cavity method

Below, we outline the steps of the cavity calculation. These represent the standard procedure, with each step detailed in the subsections that follow.

1. We begin with a steady-state community containing $S$ species and $M$ resources. We will assume that both $S$ and $M$ are large, i.e., $S \gg 1$ and $M \gg 1$.
2. To this community, we introduce a "cavity" species with index 0 and a "cavity" resource with index 0 into the community. Both the cavity species and resource are statistically identical to other species and resources in the community, respectively. This is ensured by assuming that parameters determining their interactions with other species and resources are drawn from the same distributions as for other species and resources. This results in a new community with $S + 1$ species and $M + 1$ resources. Our goal is to compute the steady-state abundances of species and resources in this larger community.
3. We calculate how the introduction of the cavity species and resource affect the resident community members. Since both $S$ and $M$ are large, adding only one cavity species and resource acts as a small perturbation to the dynamics of the original community residents. We use this to approximately compute the steady-state abundances of the resident species and resources after the introduction of these cavity variables.
4. The changes to the steady-state abundances of the residents will in turn affect the dynamics of the cavity species and resource. This allows us to derive self-consistency equations for the steady-state abundances of the cavity species and resource respectively. We use this to derive an ansatz for the distribution of the abundance of the cavity species $N_0$ and resource $R_0$.
5. Since the cavity species and resource are statistically identical to the resident species and resources respectively, we invoke self-averaging. Thus the distribution of all species abundances will follow that of the cavity species, and of all resource abundances will follow that of the cavity resource. In other words, the abundance of a cavity species across different community realisation is equivalent to the abundance distribution of all resident species in a single community.
6. To assess community stability, we introduce small perturbations to the abundances of all surviving species and resources. We then compute their sensitivity to these perturbations. When the sensitivities to perturbations are no longer finite or well-defined, the cavity solution is no longer stable and community dynamics do not reach steady-state.

#### C.2 Step 1: Steady-state community before introducing the cavity species and resource

We start with a steady-state community with $S$ species before the cavity variables 0 are introduced. The steady-state abundances of resident species $i \setminus 0$ (read as "$i$ without 0") are described by the following equations:

$$\frac{\mathrm{d}N_{i\setminus 0}}{\mathrm{d}t} = 0 = N_{i\setminus 0}\left(\sum_{\alpha=1}^{M} y_{i\alpha}c_{i\alpha}R_{\alpha\setminus 0} - d_i\right), \tag{12}$$

By substituting the the statistics of our parameters (Eqs. (2)–(6)), we can write the per capita growth rate in terms of an average and its fluctuations, as:

$$0 = N_{i\setminus 0}\Bigg(\underbrace{\frac{\mu_y\mu_c}{M}\sum_{\alpha=1}^{M} R_{\alpha\setminus 0}}_{\substack{\text{average total growth rate}\\ \text{from consuming resources}}} + \underbrace{\sum_{\alpha=1}^{M}\left(\frac{\mu_y\sigma_c}{\sqrt{M}}z_{c,i\alpha} + \frac{\mu_c\sigma_y}{M}z_{y,i\alpha} + \frac{\sigma_y\sigma_c}{\sqrt{M}}z_{y,i\alpha}z_{c,i\alpha}\right)R_{\alpha\setminus 0}}_{\text{fluctuations}} - d_i\Bigg). \tag{13}$$

Similarly, for resident resources $\alpha \setminus 0$ , we have:

$$\frac{\mathrm{d}R_{\alpha\setminus 0}}{\mathrm{d}t} = 0 = R_{\alpha\setminus 0}\left(b_\alpha - R_{\alpha\setminus 0} - \sum_{i=1}^{S} c_{i\alpha}N_{i\setminus 0}\right) = R_{\alpha\setminus 0}\Bigg(b_\alpha - \underbrace{\frac{\mu_c}{M}\sum_{i=1}^{S} N_{i\setminus 0}}_{\substack{\text{average total}\\ \text{consumption rate}}} - \underbrace{\sum_{i=1}^{S}\frac{\sigma_c}{\sqrt{M}}z_{y,i\alpha}N_{i\setminus 0}}_{\text{fluctuations}} - R_{\alpha\setminus 0}\Bigg). \tag{14}$$

#### C.3 Steps 2 and 3: Introducing the cavity species and resource, and their effect on community residents

Next, we invade the existing community with the cavity species and resource, and allow the system to reach a new steady state. The introduction of the cavity species and resource result in changes in the steady-state abundances of resident community species from $N_{i\setminus 0} \to N_i$ and resources from $R_{\alpha\setminus 0} \to R_\alpha$. The new steady-state abundances for residents after invasion by the cavity variables thus obey the following equations:

$$\frac{\mathrm{d}N_i}{\mathrm{d}t} = 0 = \overset{\substack{\text{perturbed}\\ \text{abundance}\\ \text{of species } i\\ \downarrow}}{N_i}\Bigg(\sum_{\alpha=1}^{M} y_{i\alpha}c_{i\alpha}\overset{\substack{\text{perturbed}\\ \text{abundance}\\ \text{of resource } \alpha\\ \downarrow}}{R_\alpha} - d_i + y_{i0}c_{i0}\overset{\substack{\text{abundance of cavity}\\ \text{resource 0}\\ \downarrow}}{R_0}\Bigg) \tag{15}$$

Notice that the last term in this equation contains the effect of the cavity resource on the dynamics of the abundance of species $i$. This effect can be interpreted as a small perturbation to the death rate of species $i$:

$$d_i \xrightarrow{\text{perturb}} d_i - \underbrace{y_{i0}c_{i0}R_0}_{\text{perturbation } \Delta d_i} . \tag{16}$$

When the number of resources is large $M \gg 1$, this perturbation is small compared to the unperturbed death rate $d_i$ since $c_{i0} \sim \frac{1}{M}$ (see Appendix B.3).

Similarly, the new steady-state abundances of resident resources after invasion by the cavity variables obey the following equations:

$$\frac{\mathrm{d}R_\alpha}{\mathrm{d}t} = 0 = R_\alpha \left( b_\alpha - c_{0\alpha} \overset{\substack{\text{abundance of cavity}\\ \text{species 0}\\ \downarrow}}{N_0} - R_\alpha - \sum_{i=1}^{S} c_{i\alpha} N_i \right) \tag{17}$$

Using similar logic as for species, the second term in this equation contains the effect of the cavity species on the dynamics of the abundance of resource $\alpha$. This effect can be interpreted as a small perturbation to the supply rate of resource $\alpha$:

$$b_\alpha \xrightarrow{\text{perturb}} b_\alpha - \underbrace{c_{0\alpha} N_0}_{\text{perturbation } \Delta b_\alpha} . \tag{18}$$

This perturbation is also small compared to the unperturbed supply rate $b_\alpha$ when $M \gg 1$ since $c_{0\alpha} \sim \frac{1}{M}$. Thus, we observe that both resident species and resource dynamics experience a small perturbation to their death and supply rates respectively. This allows us to relate the post-perturbation abundance of species $i$, $N_i$ with its pre-perturbation abundance $N_{i\backslash 0}$ using a linear response approximation, given by:

$$\begin{aligned} N_i &\approx N_{i\backslash 0} + \sum_{j=1}^{S} \underbrace{\frac{\partial N_i}{\partial d_j}}_{v_{ij}^{(N)}} \Delta d_j + \sum_{\beta=1}^{M} \underbrace{\frac{\partial N_i}{\partial b_\beta}}_{\chi_{i\beta}^{(N)}} \Delta b_\beta . \\ &\approx N_{i\backslash 0} - \sum_{j=1}^{S} v_{ij}^{(N)} (y_{j0} c_{j0} R_0) - \sum_{\beta=1}^{M} \chi_{i\beta}^{(N)} (c_{0\beta} N_0). \end{aligned} \tag{19}$$

Here, we have introduced susceptibility matrices $v_{ij}^{(N)}$ and $\chi_{i\beta}^{(N)}$ characterising the response of the abundance of species $i$ to small perturbations in the death rate of species $j$ and supply rate of resource $\beta$, respectively. These matrices are in general large and unknown, as they depend on the properties of the dynamics. Nevertheless, we introduce them here for convenience, with the hope of computing the required unknowns later in a self-consistent manner.

Similarly, using a linear response approximation we can also relate the post-perturbation steady-state abundance of resource $\alpha$, $R_\alpha$ with its post-perturbation steady-state abundance $R_{\alpha\backslash 0}$, as:

$$\begin{aligned} R_\alpha &\approx R_{\alpha\backslash 0} + \sum_{j=1}^{S} \underbrace{\frac{\partial R_\alpha}{\partial d_j}}_{v_{\alpha j}^{(R)}} \Delta d_j + \sum_{\beta=1}^{M} \underbrace{\frac{\partial R_\alpha}{\partial b_\beta}}_{\chi_{\alpha\beta}^{(R)}} \Delta b_\beta \\ &\approx R_{\alpha\backslash 0} - \sum_{j=1}^{S} v_{\alpha j}^{(R)} (y_{j0} c_{j0} R_0) - \sum_{\beta=1}^{M} \chi_{\alpha\beta}^{(R)} (c_{0\beta} N_0). \end{aligned} \tag{20}$$

Here we introduced the susceptibility matrices $v_{\alpha j}^{(R)}$ and $\chi_{\alpha\beta}^{(R)}$ characterising the response of the abundance of resource $\alpha$ to small perturbations in the death rate of species $j$ and supply rate of resource $\beta$, respectively. These matrix elements are also unknowns, and the relevant quantities will need to be determined self-consistently. Below, we collect all the four susceptibility matrices defined for the model for reference later, as

$$v_{ij}^{(N)} = \frac{\partial N_i}{\partial d_j}, \quad v_{\alpha j}^{(R)} = \frac{\partial R_i}{\partial d_j}, \quad \chi_{i\beta}^{(N)} = \frac{\partial N_i}{\partial b_\beta}, \quad \chi_{\alpha\beta}^{(N)} = \frac{\partial R_\alpha}{\partial b_\beta}, \tag{21}$$

Note that $v_{ij}^{(N)}$ and $\chi_{\alpha\beta}^{(N)}$ are known as the “diagonal-block" susceptibilities and $v_{\alpha j}^{(R)}$ and $\chi_{i\beta}^{(N)}$ are the “off-block" susceptibilities.

#### C.4 Step 4: Self-consistent dynamics of the cavity species and resource

Once the cavity species and resource invade the community and reach steady state, they will have abundances $N_0$ and $R_0$. These abundances are given by solutions to the steady-state condition for $N_0$ and $R_0$, given as:

$$\frac{\mathrm{d}N_0}{\mathrm{d}t} = 0 = N_0 \left( \sum_{\alpha=1}^{M} y_{0\alpha} c_{0\alpha} R_\alpha + y_{00} c_{00} R_0 - d_0 \right) \tag{22}$$

and:

$$\frac{\mathrm{d}R_0}{\mathrm{d}t} = 0 = R_0 \left( b_0 - R_0 - \sum_{i=1}^{S} c_{i0} N_i - c_{00} N_0 \right). \tag{23}$$

Note that these equations naturally depend on the steady-state abundances of all community residents $N_i$ and $R_\alpha$, which we have already computed in Eqs. (19) and (20).

##### C.4.1 Cavity species equation

We will first solve for the steady-state abundance of the cavity species $N_0$. Firstly, we will substitute in our linear approximation for $R_\alpha$ from eq. (19) to obtain

$$0 = N_0 \left( \sum_{\alpha=1}^{M} y_{0\alpha} c_{0\alpha} \left( R_{\alpha\setminus 0} - \sum_{j=1}^{S} v_{\alpha j}^{(R)} (y_{j0} c_{j0} R_0) - \sum_{\beta=1}^{M} \chi_{\alpha\beta}^{(R)} (c_{0\beta} N_0) \right) + y_{00} c_{00} R_0 - d_0 \right). \tag{24}$$

Next, we substitute model parameters with the distributions they are sampled from (Eqs. (2)–(6)) i.e., the ensemble, and obtain:

$$\begin{aligned} 0 = N_0 \Bigg( \sum_{\alpha=1}^{M} \underbrace{\left( \mu_y + \sigma_y z_{y,0\alpha} \right)}_{y_{0\alpha}} \underbrace{\left( \frac{\mu_c}{M} + \frac{\sigma_c}{\sqrt{M}} z_{c,0\alpha} \right)}_{c_{0\alpha}} \Bigg( R_{\alpha\setminus 0} - \sum_{j=1}^{S} v_{\alpha j}^{(R)} \underbrace{\left( \mu_y + \sigma_y z_{y,j0} \right)}_{y_{j0}} \underbrace{\left( \frac{\mu_c}{M} + \frac{\sigma_c}{\sqrt{M}} z_{c,j0} \right)}_{c_{j0}} R_0 \\ - \sum_{\beta=1}^{M} \chi_{\alpha\beta}^{(R)} \underbrace{\left( \frac{\mu_c}{M} + \frac{\sigma_c}{\sqrt{M}} z_{c,0\beta} \right)}_{c_{0\beta}} N_0 \Bigg) \\ + \underbrace{\left( \mu_y + \sigma_y z_{y,00} \right)}_{y_{00}} \underbrace{\left( \frac{\mu_c}{M} + \frac{\sigma_c}{\sqrt{M}} z_{c,00} \right)}_{c_{00}} R_0 - \underbrace{\left( \mu_d + \sigma_d z_{d,0} \right)}_{d_0} \Bigg). \end{aligned} \tag{25}$$

We then expand out these brackets and split the terms up in terms of averages and fluctuations (terms containing standard normal variables).

$$
\begin{aligned}
0 = N_0 \Bigg( & \overbrace{\frac{\mu_y\mu_c}{M}\sum_{\alpha=1}^{M} R_{\alpha\backslash 0} + \frac{\mu_y\mu_c}{M}R_0 - \mu_d - \left(\frac{\mu_y\mu_c}{M}\right)^2 R_0 \sum_{\alpha=1}^{M}\sum_{j=1}^{S} v_{\alpha j}^{(R)} - \frac{\mu_y\mu_c^2}{M^2}N_0\sum_{\alpha=1}^{M}\sum_{\beta=1}^{M}\chi_{\alpha\beta}^{(R)}}^{\textbf{Means}} \\
& \overbrace{+ \frac{\mu_y\sigma_c}{\sqrt{M}}\sum_{\alpha=1}^{M} z_{c,0\alpha}R_{\alpha\backslash 0} + \frac{\mu_y\sigma_c}{\sqrt{M}} z_{c,00}R_0 + \frac{\mu_c\sigma_y}{M}\sum_{\alpha=1}^{M} z_{y,0\alpha}R_{\alpha\backslash 0} + \frac{\mu_c\sigma_y}{M}z_{y,00}R_0}^{\textbf{Fluctuations}} \\
& + \frac{\sigma_y\sigma_c}{\sqrt{M}}\sum_{\alpha=1}^{M} z_{y,0\alpha}z_{c,0\alpha}R_{\alpha\backslash 0} + \frac{\sigma_y\sigma_c}{\sqrt{M}} z_{y,00}z_{c,00}R_0 \\
& - \frac{\mu_y^2\mu_c\sigma_c}{M^{3/2}}R_0\sum_{\alpha=1}^{M}\sum_{j=1}^{S} z_{c,0\alpha}v_{\alpha j}^{(R)} - \frac{\mu_y\mu_c^2\sigma_y}{M^2}R_0\sum_{\alpha=1}^{M}\sum_{j=1}^{S} z_{y,0\alpha}v_{\alpha j}^{(R)} - \frac{\mu_y\mu_c\sigma_y\sigma_c}{M^{3/2}}R_0\sum_{\alpha=1}^{M}\sum_{j=1}^{S} z_{y,0\alpha}z_{c,0\alpha}v_{\alpha j}^{(R)} \\
& - \frac{\mu_y^2\mu_c\sigma_c}{M^{3/2}}R_0\sum_{\alpha=1}^{M}\sum_{j=1}^{S} z_{c,j0}v_{\alpha j}^{(R)} - \frac{\mu_y^2\sigma_c^2}{M}R_0\sum_{\alpha=1}^{M}\sum_{j=1}^{S} z_{c,0\alpha}z_{c,j0}v_{\alpha j}^{(R)} - \frac{\mu_y\mu_c\sigma_y\sigma_c}{M^{3/2}}R_0\sum_{\alpha=1}^{M}\sum_{j=1}^{S} z_{y,0\alpha}z_{c,j0}v_{\alpha j}^{(R)} \\
& - \frac{\mu_y\sigma_y\sigma_c^2}{M}R_0\sum_{\alpha=1}^{M}\sum_{j=1}^{S} z_{y,0\alpha}z_{c,0\alpha}z_{c,j0}v_{\alpha j}^{(R)} \\
& - \frac{\mu_y\mu_c^2\sigma_y}{M^2}R_0\sum_{\alpha=1}^{M}\sum_{j=1}^{S} z_{y,j0}v_{\alpha j}^{(R)} - \frac{\mu_y\mu_c\sigma_y\sigma_c}{M^{3/2}}R_0\sum_{\alpha=1}^{M}\sum_{j=1}^{S} z_{c,0\alpha}z_{y,j0}v_{\alpha j}^{(R)} - \frac{\mu_c^2\sigma_y^2}{M^2}R_0\sum_{\alpha=1}^{M}\sum_{j=1}^{S} z_{y,0\alpha}z_{y,j0}v_{\alpha j}^{(R)} \\
& - \frac{\mu_c\sigma_y^2\sigma_c}{M^{3/2}}R_0\sum_{\alpha=1}^{M}\sum_{j=1}^{S} z_{y,0\alpha}z_{c,0\alpha}z_{y,j0}v_{\alpha j}^{(R)} \\
& - \frac{\mu_y\mu_c\sigma_y\sigma_c}{M^{3/2}}R_0\sum_{\alpha=1}^{M}\sum_{j=1}^{S} z_{y,j0}z_{c,j0}v_{\alpha j}^{(R)} - \frac{\mu_y\sigma_y\sigma_c^2}{M}R_0\sum_{\alpha=1}^{M}\sum_{j=1}^{S} z_{c,0\alpha}z_{y,j0}z_{c,j0}v_{\alpha j}^{(R)} \\
& - \frac{\mu_c\sigma_y^2\sigma_c}{M^{3/2}}R_0\sum_{\alpha=1}^{M}\sum_{j=1}^{S} z_{y,0\alpha}z_{y,j0}z_{c,j0}v_{\alpha j}^{(R)} - \frac{\sigma_y^2\sigma_c^2}{M}R_0\sum_{\alpha=1}^{M}\sum_{j=1}^{S} z_{y,0\alpha}z_{c,0\alpha}z_{y,j0}z_{c,j0}v_{\alpha j}^{(R)} \\
& - \frac{\mu_y\mu_c\sigma_c}{M^{3/2}}N_0\sum_{\alpha=1}^{M}\sum_{\beta=1}^{M} z_{c,0\alpha}\chi_{\alpha\beta}^{(R)} - \frac{\mu_c^2\sigma_y}{M^2}N_0\sum_{\alpha=1}^{M}\sum_{\beta=1}^{M} z_{y,0\alpha}\chi_{\alpha\beta}^{(R)} - \frac{\mu_c\sigma_y\sigma_c}{M^{3/2}}N_0\sum_{\alpha=1}^{M}\sum_{\beta=1}^{M} z_{y,0\alpha}z_{c,0\alpha}\chi_{\alpha\beta}^{(R)} \\
& - \frac{\mu_y\mu_c\sigma_c}{M^{3/2}}N_0\sum_{\alpha=1}^{M}\sum_{\beta=1}^{M} z_{c,0\beta}\chi_{\alpha\beta}^{(R)} - \frac{\mu_y\sigma_c^2}{M}N_0\sum_{\alpha=1}^{M}\sum_{\beta=1}^{M} z_{c,0\alpha}z_{c,0\beta}\chi_{\alpha\beta}^{(R)} - \frac{\mu_c\sigma_y\sigma_c}{M^{3/2}}N_0\sum_{\alpha=1}^{M}\sum_{\beta=1}^{M} z_{y,0\alpha}z_{c,0\beta}\chi_{\alpha\beta}^{(R)} \\
& - \frac{\sigma_y\sigma_c^2}{M}N_0\sum_{\alpha=1}^{M}\sum_{\beta=1}^{M} z_{y,0\alpha}z_{c,0\alpha}z_{c,0\beta}\chi_{\alpha\beta}^{(R)} - \sigma_d z_{d,0}\Bigg). \qquad (26)
\end{aligned}
$$

To solve this equation and obtain the steady-state abundance $N_0$, we first note that many of the terms in Eq. (26) involve sums over many random variables (since the sums go over $S$ or $M$, both of which we assume are large). This means that we can assume that these sums follow a Gaussian distribution, according to the Central Limit Theorem. Subsequently, we can reduce the term in the bracket on the r.h.s of eq. (26) to a single Gaussian variable with a given mean ($\mu$) and standard deviation ($\sigma$):

$$0 = N_0\left(\mu + \sigma Z_N\right).$$

To determine the mean and variance of this Gaussian variable, we must systematically computing the first ($\langle X\rangle$) and second ($\langle X^2\rangle$) moments of each term because $\mu = \left(\sum\langle \text{terms inside bracket}\rangle\right)$ and $\sigma = \sqrt{\sum\langle(\text{terms inside bracket})^2\rangle - \mu^2}$.

We will find that when $M \gg 1$, some of these terms become sub-leading compared to the leading order term in $M$. We will neglect sub-leading terms in our final solution, which greatly simplifies the expression without worsening its agreement with simulations.

#### Rules of thumb for calculating statistical moments in the cavity calculation

Before we calculate the first and second order moments, we will first outline some assumptions and mathematical rules we use to calculate them.

**When to retain leading-order terms and discard sub-leading terms**

As we calculate the first and second moments of the sums in eq. (26), we will find that terms are of different orders of magnitude in $M$: $O(1)$, $O(1/\sqrt{M})$, $O(1/M)$ etc. Cavity calculations in thermodynamic limit ($M \to \infty$) typically discard sub-leading $M$-dependent terms and only retain the leading order $O(1)$ terms. Nevertheless, these expressions accurately capture simulations performed on ecosystems with finitely-many species and resources, suggesting that the discarded terms remain negligible even outside the thermodynamic limit. Since our calculation is performed in the limit where $M$ large but finite ($1 \ll M \ll \infty$), like those simulations, we assume that we retain terms that tend to be leading order in the thermodynamic-limit discard terms that are sub-leading. Although this approach is naive, we believe it is justified given the strong agreement between our analytical calculations and simulations.

In our final solution, we find that the leading order terms are of order $\geq O(1/\sqrt{M})$ while sub-leading terms are usually smaller than $O(1/\sqrt{M})$. However, we claim some $O(1/\sqrt{M})$ are still sub-leading based on the arguments made earlier, and subsequently discard them. These terms will be discussed later in the calculation.

**Keeping or discarding terms containing susceptibilities**

Similarly, we will keep only leading-order terms containing susceptibilities. The magnitude and order of these terms is either assumed self-consistently, or have been demonstrated in previous work [3–5, 8]. These assumptions are self-consistently justified, as evidenced by the agreement between our simulations and calculations. They follow below.

Firstly, we assume that the first and the second moments of off-block susceptibilities ($v^{(R)}_{\alpha j}$ and $\chi^{(N)}_{i\beta}$) are always sub-leading or approximately zero, and thus can be ignored.

The on-block susceptibilities ($v^{N}_{ij}$ and $\chi^{(R)}_{\alpha\beta}$) are diagonally dominant. We break them down into their average non-diagonal and diagonal entries. We assume that the first moment of the diagonal entries (related to the trace of the susceptibilities) are

$$\langle v^{(N)}_{ii} \rangle = \frac{1}{S}\sum_{i}^{S} v^{(N)}_{ii} = O(1); \quad \langle \chi^{(R)}_{\alpha\alpha} \rangle = \frac{1}{M}\sum_{\alpha}^{M} \chi^{(R)}_{\alpha\alpha} = O(1), \tag{27}$$

while the first moment of the non-diagonal entries are 0:

$$\langle v^{(N)}_{i,j\neq i} \rangle \approx 0; \quad \langle \chi^{(R)}_{\alpha,\alpha\neq\beta} \rangle \approx 0. \tag{28}$$

The second moment of all entries in the diagonal-block matrices are

$$\langle (v^{(N)}_{ij})^2 \rangle = \frac{1}{S^2}\sum_{ij}^{S} (v^{(N)}_{ij})^2 = O\left(\frac{1}{S}\right); \quad \langle (\chi^{(R)}_{\alpha\beta})^2 \rangle = \frac{1}{M^2}\sum_{\alpha\beta}^{M} (\chi^{(R)}_{\alpha\beta})^2 = O\left(\frac{1}{M}\right). \tag{29}$$

**We assume most variables are weakly correlated, so we treat them as independent variables.**

This allows us to write $\langle X^p Y^q \rangle = \langle X^p \rangle \langle Y^q \rangle$. For example,

$$\left\langle \left( \frac{\mu_y \sigma_c}{\sqrt{M}} \sum_{\alpha=1}^{M} z_{c,0\alpha} R_{\alpha\backslash 0} \right)^2 \right\rangle = \frac{(\mu_y\sigma_c)^2}{M} \sum_{\alpha,\beta}^{M} \left\langle z_{c,0\alpha} z_{c,0\beta} R_{\alpha\backslash 0} R_{\beta\backslash 0} \right\rangle$$

$$= \frac{(\mu_y\sigma_c)^2}{M} \left( \sum_{\alpha}^{M} \underbrace{\left\langle z_{c,0\alpha}^2 \right\rangle}_{1} \underbrace{R_{\alpha\backslash 0}^2}_{\langle R^2 \rangle} + \sum_{\alpha,\beta\neq\alpha}^{M} \underbrace{\left\langle z_{c,0\alpha} z_{c,0\beta} \right\rangle}_{0} \underbrace{\left\langle R_{\alpha\backslash 0} R_{\beta\backslash 0} \right\rangle}_{\approx 0} \right) \tag{30}$$

$$= \frac{(\mu_y\sigma_c)^2}{M} M \langle R^2 \rangle = (\mu_y\sigma_c)^2 \langle R^2 \rangle.$$

Since $z$ and $R$ are uncorrelated and since $\langle z_{c,0\alpha} \rangle = 0$. Similarly, 0 is assigned to terms that contain, for example, $\sum_{\alpha=1}^{M} z_{y,0\alpha} z_{c,0\alpha}$, $\sum_{\alpha=1}^{M} \sum_{j=1}^{S} z_{y,j0} v_{\alpha j}^{(R)}$, $\sum_{\alpha=1}^{M} \sum_{j=1}^{S} z_{y,j0} z_{c,j0} v_{\alpha j}^{(R)}$, and $\sum_{\alpha=1}^{M} \sum_{\beta=1}^{M} \chi_{\alpha\beta}^{(R)} z_{c,0\beta}$.

As an example for terms that we retain, consider

$$\left\langle -\frac{\mu_y \sigma_c^2}{M} N_0 \sum_{\alpha=1}^{M} \sum_{\beta=1}^{M} \chi_{\alpha\beta}^{(R)} z_{c,0\alpha} z_{c,0\beta} \right\rangle = -\frac{\mu_y \sigma_c^2 N_0}{M} \sum_{\alpha,\beta}^{M} \langle \chi_{\alpha\beta}^{(R)} \rangle \langle z_{c,0\alpha} z_{c,0\beta} \rangle$$

$$= -\frac{\mu_y \sigma_c^2 N_0}{M} \left( \sum_{\alpha=1}^{M} \langle \chi_{\alpha\alpha}^{(R)} \rangle \langle z_{c,0\alpha}^2 \rangle + \sum_{\alpha,\beta\neq\alpha}^{M} \langle \chi_{\alpha\beta}^{(R)} \rangle \langle z_{c,0\alpha} z_{c,0\beta} \rangle \right). \tag{31}$$

When $\alpha = \beta$, which occurs $M$ times, these entries have a non-zero value since $\langle z_{c,0\alpha}^2 \rangle = 1$, and all other entries where $\alpha \neq \beta$ are 0:

$$-\frac{\mu_y \sigma_c^2 N_0}{M} \left( \sum_{\alpha=1}^{M} \langle \chi_{\alpha\alpha}^{(R)} \rangle \langle z_{c,0\alpha}^2 \rangle + \sum_{\alpha,\beta\neq\alpha}^{M} \langle \chi_{\alpha\beta}^{(R)} \rangle \langle z_{c,0\alpha} z_{c,0\beta} \rangle \right)$$

$$= -\frac{\mu_y \sigma_c^2 N_0}{M} M \left( \cancelto{\chi^{(R)}}{\langle \chi_{\alpha\alpha}^{(R)} \rangle} \cancelto{1}{\langle z_{c,0\alpha}^2 \rangle} - (M^2 - M) \langle \chi_{\alpha\beta\neq\alpha}^{(R)} \rangle \cancelto{0}{\langle z_{c,0\alpha} z_{c,0\beta\neq\alpha} \rangle} \right) \tag{32}$$

$$= -\mu_y \sigma_c^2 \chi^{(R)} N_0.$$

where $\chi^{(R)}$ is the average diagonal entry of the susceptibility matrix $\chi_{\alpha\beta}^{(R)}$, and quantifies the average change in the abundance of resource $\alpha$ upon a small perturbation to its intrinsic supply rate. Therefore, even though the susceptibility matrix $\chi_{\alpha\beta}^{(R)}$ is large and unknown, the only quantity that matters to compute steady-state properties is its average diagonal entry. This quantity is still unknown, and we will later use its definition to arrive at a self-consistency equation to solve for it.

Now we have outlined out rules, we can now compute the first and second moments of the terms in eq. (26).

#### First moments

##### Means

The first leading order term in the means is

$$\left\langle \frac{\mu_y \mu_c}{M} \sum_{\alpha=1}^{M} R_{\alpha\setminus 0} + \frac{\mu_y \mu_c}{M} R_0 \right\rangle = \frac{\mu_y \mu_c}{M} \left( \sum_{\alpha=1}^{M} \cancelto{\langle R\rangle}{\langle R_{\alpha\setminus 0}\rangle} + \cancelto{\langle R\rangle}{\langle R_0\rangle} \right) = \frac{\mu_y \mu_c}{M} (M+1) \langle R \rangle \approx \mu_y \mu_c \langle R \rangle. \tag{33}$$

Note that here we have introduced the mean steady-state species abundance $\langle R\rangle = \frac{1}{M}\sum_{\alpha=1}^{M} R_{\alpha\setminus 0}$ or $\frac{1}{M}\sum_{\alpha=1}^{M} R_{\alpha}$, an unknown which we must later determine self-consistently. Because we assume $M \gg 1$, we assume adding one more resource (or species) i.e., the cavity resource does not significantly change the statistical properties of the existing community. In other words, a community with $S$ species and $M$ resources has a similar average and fluctuations in net growth rate compared to community with $S+1$ species and $M+1$ resources. Therefore, the average net growth rate of the original community members does not significantly change upon being invaded by the cavity species and resource.

The other leading-order term is

$$\langle -\mu_d \rangle = -\mu_d. \tag{34}$$

The other sums contributing the means are sub-leading or negligible when $M$ is large, so are discarded:

$$\left\langle -\left(\frac{\mu_y \mu_c}{M}\right)^2 R_0 \sum_{\alpha=1}^{M} \sum_{j=1}^{S} v_{\alpha j}^{(R)} \right\rangle = -\left(\frac{\mu_y \mu_c}{M}\right)^2 R_0 \left\langle \sum_{\alpha=1}^{M} \sum_{j=1}^{S} \underbrace{v_{\alpha j}^{(R)}}_{O(1/\sqrt{M})} \right\rangle \approx 0. \tag{35}$$

$$\left\langle -\frac{\mu_y \mu_c^2}{M^2} N_0 \sum_{\alpha=1}^{M} \sum_{\beta=1}^{M} \chi_{\alpha\beta}^{(R)} \right\rangle = -\frac{\mu_y \mu_c^2}{M^2} N_0 \left\langle \sum_{\alpha}^{M} \underbrace{\chi_{\alpha\alpha}^{(R)}}_{O(1)} + \sum_{\alpha,\beta\neq\alpha}^{M} \underbrace{\chi_{\alpha\beta\neq\alpha}^{(R)}}_{\approx 0} \right\rangle \approx -\frac{\mu_y \mu_c^2}{M^2} N_0 \, O(M) \approx 0. \tag{36}$$

The term in the angle brackets is approximately $O(M)$ because we sum over $M$ terms of $O(1)$. However, the resulting term when multiplied by $1/M^2$ is $O(1/M)$ and hence sub-leading.

**Fluctuations**

Most fluctuations have a first moment of 0. For example,

$$\left\langle \frac{\mu_y \sigma_c}{\sqrt{M}} \sum_{\alpha=1}^{M} z_{c,0\alpha} R_{\alpha\setminus 0} + \frac{\mu_y \sigma_c}{\sqrt{M}} z_{c,00} R_0 \right\rangle = \frac{\mu_y \sigma_c}{\sqrt{M}} \left( \sum_{\alpha=1}^{M} \cancelto{0}{\langle z_{c,0\alpha}\rangle} \cancelto{\langle R\rangle}{\langle R_{\alpha\setminus 0}\rangle} + \cancelto{0}{\langle z_{c,00}\rangle} \cancelto{\langle R\rangle}{\langle R_0\rangle} \right) = 0. \tag{37}$$

We will not calculate the other zero-value moments here, and leave it as an exercise to the reader.

The fluctuation that has a non-zero leading-order first moment is, as discussed in eq. (32),

$$\left\langle -\frac{\mu_y \sigma_c^2}{M} N_0 \sum_{\alpha=1}^{M} \sum_{\beta=1}^{M} \chi_{\alpha\beta}^{(R)} z_{c,0\alpha} z_{c,0\beta} \right\rangle = -\mu_y \sigma_c^2 \chi^{(R)} N_0. \tag{38}$$

**Final first moment**

Combining all terms, the final first moment is

$$\mu_y \mu_c \langle R \rangle - \mu_d - \mu_y \sigma_c^2 \chi^{(R)} N_0. \tag{39}$$

#### Second moments

Now, we will repeat the exercise above, but to calculate the expected values of the squares of each of term in eq.(26).

**Means**

The second moment of the first mean is

$$
\begin{aligned}
\left\langle \left( \frac{\mu_y \mu_c}{M} \sum_{\alpha=1}^{M} R_{\alpha\backslash 0} + \frac{\mu_y \mu_c}{M} R_0 \right)^2 \right\rangle &= \left(\frac{\mu_y \mu_c}{M}\right)^2 \left( \sum_{\alpha,\beta}^{M} \langle R_{\alpha\backslash 0} R_{\beta\backslash 0} \rangle + 2 \sum_{\alpha=1}^{M} \langle R_{\alpha\backslash 0} R_0 \rangle + \langle R_0^2 \rangle \right) \\
&= \left(\frac{\mu_y \mu_c}{M}\right)^2 \left( \sum_{\alpha=1}^{M} \langle R^2_{\alpha\backslash 0} \rangle + \sum_{\alpha,\beta\neq\alpha}^{M} \langle R_{\alpha\backslash 0} R_{\beta\neq\alpha\backslash 0} \rangle + 2 \sum_{\alpha=1}^{M} \langle R_{\alpha\backslash 0} R_0 \rangle + \langle R_0^2 \rangle \right).
\end{aligned}
\tag{40}
$$

Since $R_{\alpha\backslash 0}$, $R_{\beta\neq\alpha\backslash 0}$ and $R_0$ are weakly correlated, this expression becomes

$$
\begin{aligned}
&\left(\frac{\mu_y \mu_c}{M}\right)^2 \left( \sum_{\alpha=1}^{M} \langle R^2_{\alpha\backslash 0} \rangle + \sum_{\alpha,\beta\neq\alpha}^{M} \langle R_{\alpha\backslash 0} R_{\beta\neq\alpha\backslash 0} \rangle + 2 \sum_{\alpha=1}^{M} \langle R_{\alpha\backslash 0} R_0 \rangle + \langle R_0^2 \rangle \right). \\
&= \left(\frac{\mu_y \mu_c}{M}\right)^2 \left( M \cancelto{\langle R^2\rangle}{\langle R^2_{\alpha\backslash 0} \rangle} + (M^2 - M) \cancelto{\langle R\rangle}{\langle R_{\alpha\backslash 0}\rangle} \cancelto{\langle R\rangle}{\langle R_{\beta\backslash 0}\rangle} + 2M \cancelto{\langle R\rangle}{\langle R_{\alpha\backslash 0}\rangle} \cancelto{\langle R\rangle}{\langle R_0\rangle} + \cancelto{\langle R^2\rangle}{\langle R_0^2 \rangle} \right) \\
&= \left(\frac{\mu_y \mu_c}{M}\right)^2 \left( (M+1)\langle R^2 \rangle + (M^2 + M) \langle R \rangle^2 \right) \\
&= (\mu_y \mu_c)^2 \left( \frac{M+1}{M^2} \langle R^2 \rangle + \left(1 + \frac{1}{M}\right) \langle R \rangle^2 \right).
\end{aligned}
\tag{41}
$$

where $\langle R^2 \rangle = \frac{1}{M} \sum_{\alpha=1}^{M} R_\alpha^2$ is the mean of the square, or the second moment, of the steady-state resource abundance distribution. This is another unknown which we must later determine self-consistently.

Because $M$ is large, we assume the terms of order $\leq O(1/M)$ are negligible compared to the leading order term in eq. (41) Therefore, we assume the expression reduces to

$$
\approx (\mu_y \mu_c)^2 \langle R \rangle^2. \tag{42}
$$

The second moment of the second mean term is

$$
\langle \mu_d^2 \rangle = \mu_d^2. \tag{43}
$$

The third mean term is

$$
\left\langle \left( - \left(\frac{\mu_y \mu_c}{M}\right)^2 R_0 \sum_{\alpha=1}^{M} \sum_{j=1}^{S} v^{(R)}_{\alpha j} \right)^2 \right\rangle = \left(\frac{\mu_y \mu_c}{M}\right)^4 R_0^2 \left( \sum_{\alpha}^{M} \sum_{j}^{S} \langle (v^{(R)}_{\alpha j})^2 \rangle + \sum_{\alpha,\beta\neq\alpha}^{M} \sum_{j,k\neq j}^{S} \langle v^{(R)}_{\alpha j} \rangle \langle v^{(R)}_{\beta k} \rangle \right). \tag{44}
$$

The first sum contains $\langle (v_{\alpha j}^{(R)})^2 \rangle$, which is not a well-studied quantity. Therefore it is unclear whether it is negligible, leading order or sub-leading. We will assume it is very small or sub-leading (which seems reasonable since $\langle v_{\alpha j}^{(R)} \rangle \approx 0$, and thus discard it. We know that $\langle v_{\alpha j}^{(R)} \rangle$ is sub-leading, so we discard the second sum as well, and thus the whole term.

The last term mean term is sub-leading, and is thus discarded:

$$\left\langle \left( -\frac{\mu_y \mu_c^2}{M^2} N_0 \sum_{\alpha=1}^{M} \sum_{\beta=1}^{M} \chi_{\alpha\beta}^{(R)} \right)^2 \right\rangle = \left( \frac{\mu_y \mu_c^2}{M^2} \right)^2 N_0^2 \left( M^2 \underbrace{\langle (\chi_{\alpha\beta}^{(R)})^2 \rangle}_{O(1/M)} + (M^4 - M^2) \cancelto{\approx 0}{\langle \chi_{\alpha\beta}^{(R)} \rangle \langle \chi_{cd}^{(R)} \rangle} \right)^{\approx 0}$$
$$= O\left( \frac{1}{M^3} \right) \approx 0 \tag{45}$$

when $M$ is large.

**Fluctuations**

As with the means, one can show that the second moments of many terms are negligible or sub-leading. We leave those terms as an exercise to the reader. After removing the negligible terms, we are are only left with the following:

$$\left\langle \left( \frac{\mu_y \sigma_c}{\sqrt{M}} \sum_{\alpha=1}^{M} z_{c,0\alpha} R_{\alpha\backslash 0} + \frac{\mu_y \sigma_c}{\sqrt{M}} z_{c,00} R_0 \right)^2 \right\rangle$$
$$= \frac{(\mu_y \sigma_c)^2}{M} \left\langle \left( \sum_{\alpha=1}^{M} z_{c,0\alpha} R_{\alpha\backslash 0} \right) \left( \sum_{\beta=1}^{M} z_{c,0\beta} R_{\beta\backslash 0} \right) + 2 \sum_{\alpha=1}^{M} z_{c,0\alpha} z_{c,00} R_{\alpha\backslash 0} R_0 + z_{c,00}^2 R_0^2 \right\rangle$$
$$= \frac{(\mu_y \sigma_c)^2}{M} \left( \sum_{\alpha=1}^{M} \langle z_{c,0\alpha}^2 \rangle \langle R_{\alpha\backslash 0}^2 \rangle + \sum_{\alpha,\beta\neq\alpha}^{M} \langle z_{c,0\alpha} z_{c,0\beta\neq\alpha} \rangle \langle R_{\alpha\backslash 0} R_{\beta\neq\backslash 0} \rangle + 2 \sum_{\alpha=1}^{M} \langle z_{c,0\alpha} z_{c,00} \rangle \langle R_{\alpha\backslash 0} R_0 \rangle \right.$$
$$\left. + \langle z_{c,00}^2 \rangle \langle R_0^2 \rangle \right). \tag{46}$$

The second and third terms in eq. (46) equal 0. The first and last terms have non-zero values. Thus,

$$\frac{(\mu_y \sigma_c)^2}{M} \left( M \cancelto{1}{\langle z_{c,0\alpha}^2 \rangle} \cancelto{\langle R^2 \rangle}{\langle R_{\alpha\backslash 0}^2 \rangle} + (M^2 - M) \cancelto{0}{\langle z_{c,0\alpha} z_{c,0\beta\neq\alpha} \rangle} \langle R_{\alpha\backslash 0} R_{\beta\backslash 0} \rangle + 2M \cancelto{0}{\langle z_{c,0\alpha} z_{c,00} \rangle} \langle R_{\alpha\backslash 0} R_0 \rangle + \cancelto{1}{\langle z_{c,00}^2 \rangle} \langle R_0^2 \rangle \right)^{\langle R^2 \rangle}$$
$$= \frac{(\mu_y \sigma_c)^2}{M} (M+1) \langle R^2 \rangle \approx (\mu_y \sigma_c)^2 \langle R^2 \rangle \tag{47}$$

when $M$ is large.

Continuing with the rest of non-negligible fluctuation terms, we are left with:

$$\left\langle \left( \frac{\mu_c \sigma_y}{M} \sum_{\alpha=1}^{M} z_{y,0\alpha} R_{\alpha\backslash 0} + \frac{\mu_c \sigma_y}{M} z_{y,00} R_0 \right)^2 \right\rangle \approx \frac{\mu_c^2 \sigma_y^2}{M} \langle R^2 \rangle \,. \tag{48}$$

$$\left\langle\left(\frac{\sigma_y\sigma_c}{\sqrt{M}}\sum_{\alpha=1}^{M} z_{y,0\alpha}z_{c,0\alpha}R_{\alpha\setminus 0}+\frac{\sigma_y\sigma_c}{\sqrt{M}}z_{y,00}z_{c,00}R_0\right)^2\right\rangle \approx \sigma_c^2\sigma_y^2\left\langle R^2\right\rangle. \tag{49}$$

$$\left\langle\left(-\sigma_d z_{d,0}\right)^2\right\rangle = \sigma_d^2. \tag{50}$$

$$\begin{aligned}
&\left\langle\left(\frac{\mu_y\sigma_c^2}{M}N_0\sum_{\alpha=1}^{M}\sum_{\beta=1}^{M} z_{c,0\alpha}z_{c,0\beta}\chi_{\alpha\beta}^{(R)}\right)^2\right\rangle \\
&= \left(\frac{\mu_y\sigma_c^2}{M}\right)^2 N_0^2\Bigg(\sum_{\substack{\alpha,c=\alpha,\\ \beta,d=\beta}}^{M}\cancelto{1}{\langle z_{c,0\alpha}^2\rangle}\cancelto{1}{\langle z_{c,0\beta}^2\rangle}\cancelto{\chi^{(R)}}{\langle\chi_{\alpha\alpha}^{(R)}\rangle}\cancelto{\chi^{(R)}}{\langle\chi_{\beta\beta}^{(R)}\rangle} + \sum_{\substack{\alpha,\beta\\ c=\alpha,d=\beta}}^{M}\langle z_{c,0\alpha}^2 z_{c,0\beta}^2\rangle\underbrace{\langle(\chi_{\alpha\beta}^{(R)})^2\rangle}_{O(1/M)} \\
&\qquad + \sum_{\alpha,\beta,c,d}^{M}\cancelto{0}{\langle z_{c,0\alpha}z_{c,0\beta}z_{c,0c}z_{c,0}\rangle}\langle\chi_{\alpha\beta}^{(R)}\rangle\langle\chi_{cd}^{(R)}\rangle\Bigg) \\
&= \left(\frac{\mu_y\sigma_c^2}{M}\right)^2 N_0^2\left(\sum_{\alpha\beta}^{M}(\chi^{(R)})^2+O(M)\right) \\
&\approx \left(\mu_y\sigma_c^2\right)^2 N_0^2(\chi^{(R)})^2
\end{aligned} \tag{51}$$

when $M$ is large. We assume we can neglect some of the terms in the above expression, even though they are of order $1/M$, such as the term containing $\langle(\chi_{\alpha\beta}^{(R)})^2\rangle$. However, these turn out to be much smaller than the dominant contribution we keep (as evidenced by the agreement between our calculations and simulations). Therefore, we similarly neglect other fluctuations containing $\chi_{\alpha\beta}^{(R)}$ of $O(1/M)$.

**Final second moment**

The final second moment is

$$\begin{aligned}
&(\mu_y\mu_c)^2\langle R\rangle^2+\mu_d^2+(\mu_y\sigma_c)^2\langle R^2\rangle+\frac{(\mu_c\sigma_y)^2}{M}\langle R^2\rangle+(\sigma_y\sigma_c)^2\langle R^2\rangle+\sigma_d^2+\left(\mu_y\sigma_c^2\right)^2N_0^2(\chi^{(R)})^2 \\
&= (\mu_y\mu_c)^2\langle R\rangle^2+\mu_d^2+\langle R^2\rangle\left(\sigma_c^2\left(\mu_y^2+\sigma_y^2\right)+\frac{(\mu_c\sigma_y)^2}{M}\right)+\sigma_d^2+\left(\mu_y\sigma_c^2\right)^2N_0^2(\chi^{(R)})^2.
\end{aligned} \tag{52}$$

#### Final cavity species dynamics

Now we will combine all the first and second moments computed above to write a simplified expression for eq. (26) as a Gaussian variable:

$$0 = N_0\left(\mu+\sigma Z_N\right).$$

The mean is the sum of the leading-order first moments:

$$\mu = \mu_y\mu_c\langle R\rangle-\mu_d-\mu_y\sigma_c^2\chi^{(R)}N_0$$

The variance is the sum of the leading-order second moments minus the square of the first moments:

$$
\begin{aligned}
\sigma^2 =&(\mu_y\mu_c)^2\langle R\rangle^2 + \mu_d^2 + \langle R^2\rangle\left(\sigma_c^2\left(\mu_y^2+\sigma_y^2\right)+\frac{(\mu_c\sigma_y)^2}{M}\right) + \sigma_d^2 + \left(\mu_y\sigma_c^2\right)^2 N_0^2(\chi^{(R)})^2 \\
&-\left((\mu_y\mu_c\langle R\rangle)^2 + \mu_d^2 + \left(\mu_y\sigma_c^2 N_0\chi^{(R)}\right)^2\right) \\
=&\langle R^2\rangle\left(\sigma_c^2\left(\mu_y^2+\sigma_y^2\right)+\frac{(\mu_c\sigma_y)^2}{M}\right)+\sigma_d^2,
\end{aligned}
$$

so the standard deviation is

$$
\sigma = \sqrt{\langle R^2\rangle\left(\sigma_c^2\left(\mu_y^2+\sigma_y^2\right)+\frac{(\mu_c\sigma_y)^2}{M}\right)+\sigma_d^2}.
$$

Therefore, we can write a simplified version of eq. (26) as

$$
\begin{aligned}
0 =&N_0\left(\underbrace{\mu_y\mu_c\langle R\rangle - \mu_d}_{\mu_{g_N}} - \underbrace{\mu_y\sigma_c^2\chi^{(R)}N_0}_{\text{self-feedback}} + \underbrace{\sqrt{\sigma_d^2+\langle R^2\rangle\left(\sigma_c^2(\mu_y^2+\sigma_y^2)+\frac{(\mu_c\sigma_y)^2}{M}\right)}Z_N}_{\sigma_{g_N}}\right) \\
=&N_0\left(\mu_{g_N}+\sigma_{g_N}Z_N-\mu_y\sigma_c^2\chi^{(R)}N_0\right).
\end{aligned} \tag{53}
$$

Here, we defined new variables to encapsulate the means and fluctuations in the growth rate of the cavity species into individual terms, namely $\mu_{g_N}$ and $\sigma_{g_N}$, respectively. $Z_N$ is a standard normal (Gaussian) variable $Z_N \sim \mathcal{N}(0,1)$ which arises from the sum of the non-zero leading-order fluctuations.

We can finally solve this equation to compute the cavity species' abundance ($N_0$) at steady state. There are two solutions that satisfy eq. ((53)): When $N_0 = 0$ (extinction), or $N_0 = (\mu_{g_N} + \sigma_g^N Z_N)/\mu_y\sigma_c^2\chi^{(R)}$ (survival, found from rearranging the bracket in eq. (53)). The latter solution represents a normal distribution rather than a fixed value, since $Z_N$ is a standard normal variable. This means the cavity species abundance follows a truncated normal distribution i.e., it takes values from the normal distribution when $N_0$ is positive and equals zero otherwise. Therefore, we can write the truncated normal distribution describing $N_0$ as

$$
N_0 = \max\left\{0, \frac{\mu_{g_N}+\sigma_{g_N}Z_N}{\mu_y\sigma_c^2\chi^{(R)}}\right\}. \tag{54}
$$

This equation suggests that the cavity species will either go extinct (have zero abundance) or be present at steady-state. Whether the species goes extinct or survives depends probabilistically on the specific community realisation. We assign the probability of the survival of the cavity species as $\phi_N$, which is yet another unknown whose value must be determined consistently. Note that $\phi_N = M^*/M$, where $S^*$ is the number of surviving species and $S$ is the number of available species in the pool.

##### C.4.2 Cavity resource equation

We can now repeat similar steps to solve Eq. (23) compute the steady-state abundance of resource 0. Reminding ourselves that the expression for the cavity resource dynamics is

$$0 = R_0 \left( b_0 - R_0 - \sum_{i=1}^{S} c_{i0} N_i - c_{00} N_0 \right),$$

we can substitute in our linear response approximation for $N_i$ from Eq. (19). From this we obtain

$$0 = R_0 \left( b_0 - R_0 - \sum_{i=1}^{S} c_{i0} \left( N_{i\setminus 0} - \sum_{j=1}^{S} v_{ij}^{(N)} (y_{j0} c_{j0} R_0) - \sum_{\beta=1}^{M} \chi_{i\beta}^{(N)} (c_{0\beta} N_0) \right) - c_{00} N_0 \right). \tag{55}$$

Next, we substitute the model parameters with the distributions they are sampled from (eqs. (2)–(6)) i.e., the ensemble. From this we obtain:

$$\begin{aligned} 0 = R_0 \Bigg( & \underbrace{\mu_b + \sigma_b z_{b,0}}_{b_0} - R_0 - \sum_{i=1}^{S} \underbrace{\left( \frac{\mu_c}{M} + \frac{\sigma_c}{\sqrt{M}} z_{c,i0} \right)}_{c_{i0}} \Bigg( N_{i\setminus 0} - \sum_{j=1}^{S} v_{ij}^{(N)} \underbrace{(\mu_y + \sigma_y z_{y,j0})}_{y_{j0}} \underbrace{\left( \frac{\mu_c}{M} + \frac{\sigma_c}{\sqrt{M}} z_{c,j0} \right)}_{c_{j0}} R_0 \\ & - \sum_{\beta=1}^{S} \chi_{i\beta}^{(N)} \underbrace{\left( \frac{\mu_c}{M} + \frac{\sigma_c}{\sqrt{M}} z_{c,0\beta} \right)}_{c_{0\beta}} N_0 \Bigg) \\ & - \underbrace{\left( \frac{\mu_c}{M} + \frac{\sigma_c}{\sqrt{M}} z_{c,00} \right)}_{c_{00}} N_0 \Bigg). \end{aligned} \tag{56}$$

Next, we expand out the terms in the net resource growth rate (the term inside the bracket) and partition them into means (terms that do not contain standard normal variables) and fluctuations (terms that do):

$$\begin{aligned} 0 = N_0 \Bigg( & \overbrace{\mu_b - \frac{\mu_c}{M} \sum_{i=1}^{S} N_{i\setminus 0} - \frac{\mu_c}{M} N_0 + \frac{\mu_y \mu_c^2}{M^2} R_0 \sum_{i=1}^{S} \sum_{j=1}^{S} v_{ij}^{(N)} + \frac{\mu_c^2}{M^2} N_0 \sum_{i=1}^{S} \sum_{\beta=1}^{M} \chi_{i\beta}^{(N)}}^{\textbf{Means}} - R_0 \\ & \overbrace{\begin{aligned} & + \sigma_b z_{b,0} \\ & - \frac{\sigma_c}{\sqrt{M}} \sum_{i=1}^{S} z_{c,i0} N_{i\setminus 0} - \frac{\sigma_c}{\sqrt{M}} z_{c,00} N_0 \\ & + \frac{\mu_y \mu_c \sigma_c}{M^{3/2}} R_0 \sum_{i=1}^{S} \sum_{j=1}^{S} z_{c,j0} v_{ij}^{(N)} + \frac{\mu_c^2 \sigma_y}{M^2} R_0 \sum_{i=1}^{S} \sum_{j=1}^{S} z_{y,j0} v_{ij}^{(N)} + \frac{\mu_c \sigma_y \sigma_c}{M^{3/2}} R_0 \sum_{i=1}^{S} \sum_{j=1}^{S} z_{y,j0} z_{c,j0} v_{ij}^{(N)} \\ & + \frac{\mu_y \mu_c \sigma_c}{M^{3/2}} R_0 \sum_{i=1}^{S} \sum_{j=1}^{S} z_{c,i0} v_{ij}^{(N)} + \frac{\mu_y \sigma_c^2}{M} R_0 \sum_{i=1}^{S} \sum_{j=1}^{S} z_{c,i0} z_{c,j0} v_{ij}^{(N)} + \frac{\mu_c \sigma_y \sigma_c}{M^{3/2}} R_0 \sum_{i=1}^{S} \sum_{j=1}^{S} z_{y,j0} z_{c,i0} v_{ij}^{(N)} \\ & + \frac{\sigma_y \sigma_c^2}{M} R_0 \sum_{i=1}^{S} \sum_{j=1}^{S} z_{c,i0} z_{y,j0} z_{c,j0} v_{ij}^{(N)} \\ & + \frac{\mu_c \sigma_c}{M^{3/2}} N_0 \sum_{i=1}^{S} \sum_{\beta=1}^{M} z_{c,0\beta} \chi_{i\beta}^{(N)} \\ & + \frac{\mu_c \sigma_c}{M^{3/2}} N_0 \sum_{i=1}^{S} \sum_{\beta=1}^{M} z_{c,i0} \chi_{i\beta}^{(N)} + \frac{\sigma_c^2}{M} N_0 \sum_{i=1}^{S} \sum_{\beta=1}^{M} z_{c,i0} z_{c,0\beta} \chi_{i\beta}^{(N)} \end{aligned}}^{\textbf{Fluctuations}} \Bigg). \end{aligned} \tag{57}$$

To solve this equation and obtain the steady-state abundance $R_0$, we will again invoke the central limit theorem and assume the sums follow Gaussian distributions. Subsequently, as with the cavity species, we can reduce the term in the bracket on the r.h.s of eq. (57) to a single Gaussian variable with a given mean ($\mu$) and standard deviation ($\sigma$).

$$0 = R_0\left(\mu - R_0 + \sigma Z_R\right).$$

To determine the mean and variance of this Gaussian variable, we must also systematically compute the first and second moments of each term. Once again, we will find that when $M \gg 1$, some of these terms become sub-leading compared to the leading order term in $M$. We will neglect sub-leading. To obtain the leading-order terms, we will use the same "rules of thumb" as the cavity species.

#### First moments

##### Means

$$\langle \mu_b \rangle = \mu_b. \tag{58}$$

$$\left\langle -\frac{\mu_c}{M}\sum_{i=1}^{S} N_{i\backslash 0} - \frac{\mu_c}{M}N_0 \right\rangle = -\frac{\mu_c}{M}\left(\sum_{i=1}^{S}\cancelto{\langle N\rangle}{\langle N_{i\backslash 0}\rangle} + \cancelto{\langle N\rangle}{\langle N_0\rangle}\right) = -\frac{\mu_c(S+1)}{M}\langle N\rangle \approx -\frac{\mu_c S}{M}\langle N\rangle = -\mu_c\gamma^{-1}\langle N\rangle \tag{59}$$

when $M \gg 1$.

Note that here we have introduced the mean steady-state species abundance $\langle N\rangle = \frac{1}{S}\sum_{i=1}^{S} N_{i\backslash 0}$ or $\frac{1}{S}\sum_{i=1}^{S} N_i$, an unknown which we must later determine self-consistently. Because we assume $M \gg 1$, we assume adding one more species (or resource) i.e., the cavity species, does not significantly change the statistical properties of the existing community. In other words, a community with $S$ species and $M$ resources has a similar average and fluctuations in net growth rate compared to community with $S+1$ species and $M+1$ resources. Therefore, the average net growth rate of the original community members does not change upon being invaded by the cavity species and resource.

We can discard the first moment of the next term because it is approximately 0:

$$\left\langle \frac{\mu_y\mu_c^2}{M^2}R_0\sum_{i=1}^{S}\sum_{j=1}^{S} v_{ij}^{(N)} \right\rangle = \frac{\mu_y\mu_c^2}{M^2}R_0\left(\sum_{i}\langle v_{ii}^{(N)}\rangle + \sum_{i,j\neq i}\underbrace{\langle v_{ij\neq i}^{(N)}\rangle}_{\approx 0}\right) \approx 0. \tag{60}$$

The term in the angle brackets is approximately 0 because there are far more $v_{ij\neq i}^{(N)}$ terms — $S^2 - S$ terms — than $v_{ii}^{(N)}$ terms, of which there are $S$. Therefore, the negligible terms dominate, so the whole term is approximately 0.

We can discard the next mean term because it is sub-leading in terms of $M$:

$$\left\langle \frac{\mu_c^2}{M^2}N_0\sum_{i=1}^{S}\sum_{\beta=1}^{M}\chi_{i\beta}^{(N)} \right\rangle = \frac{\mu_c^2}{M^2}N_0\underbrace{\left\langle \sum_{i=1}^{S}\sum_{\beta=1}^{M}\chi_{i\beta}^{(N)} \right\rangle}_{O(1/S)} = O\left(\frac{1}{M^2\gamma^{-1}}\right). \tag{61}$$

##### Fluctuations

The first moments of most fluctuations are 0 or a sub-leading, and thus are left as an exercise to the reader. The only fluctuation term that has a non-zero first moments is

$$\left\langle \frac{\mu_y\sigma_c^2}{M}R_0\sum_{i=1}^{S}\sum_{j=1}^{S} z_{c,i0}z_{c,j0}v_{ij}^{(N)}\right\rangle = \frac{\mu_y\sigma_c^2}{M}R_0\left(\sum_{i=1}^{S}\cancelto{1}{\langle z_{c,i0}^2\rangle}\cancelto{v^{(N)}}{\langle v_{ii}^{(N)}\rangle} + \sum_{i,j\neq i}^{S}\cancelto{0}{\langle z_{c,i0}z_{c,j0}\rangle}\cancelto{\approx 0}{\langle v_{ij\neq i}^{(N)}\rangle}\right) \tag{62}$$

$$= \frac{\mu_y\sigma_c^2}{M}R_0(Sv^{(N)}) = \mu_y\sigma_c^2\gamma^{-1}v^{(N)}R_0.$$

We remind the readers that $v^{(N)} = 1/M\sum_{i,j}^{S} v_{ij}^{(N)} = 1/M\ \mathrm{Trace}(v_{ij}^{(N)})$ is the average diagonal entry of the susceptibility matrix $v_{ij}^{(N)}$, and quantifies the average change in the abundance of species $i$ upon a small perturbation to its death rate. This quantity is still unknown, and we will later use its definition to arrive at a self-consistency equation to solve for it.

**Final first moment**

Therefore, the final first moment contributing to the per capita growth rate of the cavity resource is

$$\mu_b - \mu_c\gamma^{-1}\langle N\rangle + \mu_y\sigma_c^2\gamma^{-1}v^{(N)}R_0. \tag{63}$$

#### Second moments

Now, we will repeat the exercise above, but to calculate the expected values of the squares of each of term in eq.(57).

**Means**

$$\langle\mu_b^2\rangle = \mu_b^2. \tag{64}$$

$$\left\langle\left(-\frac{\mu_c}{M}\sum_{i=1}^{S}N_{i\setminus 0} - \frac{\mu_c}{M}N_0\right)^2\right\rangle = \left(\frac{\mu_c}{M}\right)^2\left(\sum_{i,j}^{S}\langle N_{i\setminus 0}N_{j\setminus 0}\rangle + 2\sum_{i}^{S}\langle N_{i\setminus 0}N_0\rangle + \langle N_0^2\rangle\right)$$

$$= \left(\frac{\mu_c}{M}\right)^2\left(\sum_{i}^{S}\cancelto{\langle N^2\rangle}{\langle N_{i\setminus 0}^2\rangle} + \sum_{i,j\neq i}^{S}\cancelto{\langle N\rangle}{\langle N_{i\setminus 0}\rangle}\cancelto{\langle N\rangle}{\langle N_{j\neq i\setminus 0}\rangle} + 2\sum_{i}^{S}\cancelto{\langle N\rangle}{\langle N_{i\setminus 0}\rangle}\cancelto{\langle N\rangle}{\langle N_0\rangle} + \cancelto{\langle N^2\rangle}{\langle N_0^2\rangle}\right) \tag{65}$$

$$= \left(\frac{\mu_c}{M}\right)^2\left((S+1)\langle N^2\rangle + (S^2+S)\langle N\rangle^2\right)$$

$$= \mu_c^2\left(\frac{\langle N^2\rangle}{M} + \gamma^{-1}\left(\gamma^{-1}+\frac{1}{M}\right)\langle N\rangle^2\right).$$

When $M \gg 1$, we assume the $O(1/M)$ terms are small relative to the leading order term, so we discard them. Therefore, the term reduces to

$$\approx \mu_c^2\gamma^{-2}\langle N\rangle^2. \tag{66}$$

The third mean term is is sub-leading, and is thus discarded:

$$
\begin{aligned}
\left\langle \left( \frac{\mu_y \mu_c^2}{M^2} R_0 \sum_{i=1}^{S} \sum_{j=1}^{S} v_{ij}^{(N)} \right)^2 \right\rangle &= \left( \frac{\mu_y \mu_c^2}{M^2} \right)^2 R_0^2 \left( \sum_{ij}^{S} \underbrace{\langle (v_{ij}^{(N)})^2 \rangle}_{O(1/S)} \sum_{ijk\neq il\neq j} \underbrace{\langle v_{ij}^{(N)} \rangle}_{\approx 0} \underbrace{\langle v_{kl}^{(N)} \rangle}_{\approx 0} \right) \\
&= \left( \frac{\mu_y \mu_c^2}{M^2} \right)^2 R_0^2 \left( O(S) \right) = O\left( \frac{1}{M^3} \right) \\
&\approx 0
\end{aligned}
\tag{67}
$$

when $M \gg 1$.

The last term mean term is

$$
\left\langle \left( \frac{\mu_c^2}{M^2} N_0 \sum_{i=1}^{S} \sum_{\beta=1}^{M} \chi_{i\beta}^{(N)} \right)^2 \right\rangle = \left( \frac{\mu_c^2}{M^2} \right)^2 N_0^2 \left( \sum_{i}^{S} \sum_{\beta}^{M} \langle (\chi_{i\beta}^{(N)})^2 \rangle + \sum_{i,j\neq i}^{S} \sum_{\beta,c\neq b}^{M} \langle \chi_{i\beta}^{(N)} \rangle \langle \chi_{jc}^{(N)} \rangle \right). \tag{68}
$$

The first sum contains $\langle (\chi_{i\beta}^{(N)})^2 \rangle$, which is not a well-studied quantity. Therefore, it is unclear whether it is negligible, leading order or sub-leading. We will assume it is very small or sub-leading (which seems reasonable since $\chi_{i\beta}^{(N)})^2 \rangle \approx 0$, and thus discard it. The second sum contains $\langle (\chi_{i\beta}^{(N)}) \rangle$, which we know is approximately 0, and thus we will discard it. Therefore, the whole term is disregard.

**Fluctuations**

As with the means, one can show that the second moments of many terms are negligible or sub-leading. We leave those terms as an exercise to the reader. After removing the negligible terms, we are are only left with the following:

$$
\left\langle (\sigma_b z_{b,0})^2 \right\rangle = \sigma_b^2 \cancelto{1}{\langle z_{b,0}^2 \rangle} = \sigma_b. \tag{69}
$$

$$
\begin{aligned}
&\left\langle \left( -\frac{\sigma_c}{\sqrt{M}} \sum_{i=1}^{S} z_{c,i0} N_{i\backslash 0} - \frac{\sigma_c}{\sqrt{M}} z_{c,00} N_0 \right)^2 \right\rangle \\
&= \frac{\sigma_c^2}{M} \left( \sum_{i=1}^{S} \langle z_{c,i0}^2 \rangle \langle N_{i\backslash 0}^2 \rangle + \sum_{i,j\neq i}^{S} \langle z_{c,i0} z_{c,j0} \rangle \langle N_{i\backslash 0} N_{j\backslash 0} \rangle + \langle z_{c,00}^2 \rangle \langle N_0^2 \rangle + 2 \sum_{j=1}^{S} \langle z_{c,00} z_{c,j0} \rangle \langle N_0 N_{j\backslash 0} \rangle \right) \\
&= \frac{\sigma_c^2}{M} \left( \sum_{i=1}^{S} \cancelto{1}{\langle z_{c,i0}^2 \rangle} \cancelto{\langle N^2 \rangle}{\langle N_{i\backslash 0}^2 \rangle} + \sum_{i,j\neq i}^{S} \cancelto{0}{\langle z_{c,i0} \rangle \langle z_{c,j0} \rangle} \cancelto{\approx 0}{\langle N_{i\backslash 0} N_{j\backslash 0} \rangle} + \cancelto{1}{\langle z_{c,00}^2 \rangle} \cancelto{\langle N^2 \rangle}{\langle N_0^2 \rangle} + 2 \sum_{j=1}^{S} \cancelto{0}{\langle z_{c,00} \rangle \langle z_{c,j0} \rangle} \cancelto{0}{\langle N_0 N_{j\backslash 0} \rangle} \right) \\
&= \frac{\sigma_c^2}{M} (S+1) \langle N^2 \rangle \approx \sigma_c^2 \gamma^{-1} \langle N^2 \rangle.
\end{aligned}
\tag{70}
$$

when $S \gg 1$.

Note that here we have introduced the mean of the square of the steady-state species abundance, or equivalently the second moment of the species abundance distribution, $\langle N^2 \rangle = \frac{1}{S} \sum_{i=1}^{S} N_{i\backslash 0}^2$ or $\frac{1}{S} \sum_{i=1}^{S} N_i^2$, another unknown which we must later determine self-consistently. Because we assume $M \gg 1$, we assume adding one more species (or resource) i.e., the cavity species, does not significantly change the statistical properties of the existing community. In other words, a community with $S$ species and $M$ resources has a similar average and fluctuations in net growth rate compared to community with $S+1$ species and $M+1$ resources. Therefore, the variance in the net growth rate of the original community does not change upon being invaded by the cavity species and resource.

$$
\begin{aligned}
\left\langle \left( \frac{\mu_y\sigma_c^2}{M} R_0 \sum_{i=1}^{S} \sum_{j=1}^{S} z_{c,i0} z_{c,j0} v_{ij}^{(N)} \right)^2 \right\rangle = \left(\frac{\mu_y\sigma_c^2}{M}\right)^2 R_0^2 \Bigg( & \sum_{\substack{i,j=i\\k,l=k}}^{S} \cancelto{1}{\langle z_{c,i0}^2\rangle} \cancelto{1}{\langle z_{c,k0}^2\rangle} \cancelto{v^{(N)}}{\langle v_{ii}^{(N)}\rangle} \cancelto{v^{(N)}}{\langle v_{kk}^{(N)}\rangle} \\
& + \sum_{\substack{i,j\\k=i,l=k}}^{S} \cancelto{1}{\langle z_{c,i0}^2\rangle} \cancelto{1}{\langle z_{c,j0}^2\rangle} \underbrace{\langle (v_{ij}^{(N)})^2\rangle}_{O(1/S)} \\
& + \sum_{ijkl}^{S} \cancelto{0}{\langle z_{c,i0} z_{c,j0} z_{c,k0} z_{c,l0}\rangle} \underbrace{\langle v_{ij}^{(N)}\rangle}_{\approx 0} \underbrace{\langle v_{kl}^{(N)}\rangle}_{\approx 0} \Bigg) \\
= \left(\frac{\mu_y\sigma_c^2}{M}\right)^2 R_0^2 \Bigg( & \sum_{i,j}^{S} (v^{(N)})^2 + O(S) \Bigg).
\end{aligned} \tag{71}
$$

When $M$ is large, eq. (71) reduces to

$$
\approx \left(\frac{\mu_y\sigma_c^2}{M}\right)^2 R_0^2 \sum_{i,j}^{S} (v^{(N)})^2 = \frac{(\mu_y\sigma_c^2)^2}{M^2} R_0^2 M(M-1)(v^{(N)})^2 \approx (\mu_y\sigma_c^2)^2 R_0^2 (v^{(N)})^2. \tag{72}
$$

**Final second moment**

Therefore, the final leading order second moments are

$$
\mu_b^2 + \mu_c^2\gamma^{-2}\langle N\rangle^2 + \sigma_b^2 + \sigma_c^2\gamma^{-1}\langle N^2\rangle + (\mu_y\sigma_c^2)^2 R_0^2 (v^{(N)})^2. \tag{73}
$$

#### Final cavity resource dynamics

Now we will combine all the first and second moments computed above to write a simplified expression for eq. (57) as a Gaussian variable:

$$
0 = R_0\left(\mu - R_0 + \sigma Z_R\right).
$$

The mean is

$$
\mu = \mu_b - \mu_c\gamma^{-1}\langle N\rangle + \mu_y\sigma_c^2\gamma^{-1}v^{(N)}R_0.
$$

The variance is

$$
\begin{aligned}
\sigma^2 =& \mu_b^2 + \mu_c^2\gamma^{-2}\langle N\rangle^2 + \sigma_b^2 + \sigma_c^2\gamma^{-1}\langle N^2\rangle + (\mu_y\sigma_c^2)^2 R_0^2 (v^{(N)})^2 \\
& - \left( \mu_b^2 + \left(\mu_c\gamma^{-1}\langle N\rangle\right)^2 + \left(\mu_y\sigma_c^2\gamma^{-1}v^{(N)}R_0\right)^2 \right) \\
=& \sigma_b^2 + \sigma_c^2\gamma^{-1}\langle N^2\rangle,
\end{aligned}
$$

so the standard deviation is

$$\sigma = \sqrt{\sigma_b^2 + \sigma_c^2 \gamma^{-1} \langle N^2 \rangle}.$$

Therefore, we can write a simplified version of eq. (57) as

$$\begin{aligned} 0 =& R_0 \left( \underbrace{\mu_b - \mu_c \gamma^{-1} \langle N \rangle}_{\mu_{g_R}} + \underbrace{\mu_y \sigma_c^2 \gamma^{-1} v^{(N)} R_0 - R_0}_{\text{self-feedback}} + \underbrace{\sqrt{\sigma_b^2 + \sigma_c^2 \gamma^{-1} \langle N^2 \rangle}}_{\sigma_{g_R}} Z_R \right) \\ =& R_0 \left( \mu_{g_R} + \sigma_{g_R} Z_R - (1 - \mu_y \sigma_c^2 \gamma^{-1} v^{(N)}) R_0 \right). \end{aligned} \tag{74}$$

Here, we defined new variables to encapsulate the means and fluctuations in the net growth rate of the cavity resource into individual terms, namely $\mu_{g_R}$ and $\sigma_{g_R}$, respectively. $Z_R$ is a standard normal (Gaussian) variable $Z_R \sim \mathcal{N}(0, 1)$ which arises from the sum of the non-zero leading-order fluctuations.

We can finally solve this equation to compute the cavity resource abundance $R_0$ at steady state, similar to the case of the cavity species. In doing so, we get the following solution:

$$R_0 = \max \left\{ 0, \frac{\mu_{g_R} + \sigma_{g_R} Z_R}{1 - \mu_y \sigma_c^2 \gamma^{-1} v^{(N)}} \right\}. \tag{75}$$

This equation suggests that the cavity resource will either go extinct (have zero abundance) or be present at steady-state. Whether the resource goes extinct or survives depends probabilistically on the specific community realisation. We assign the probability of the survival of the cavity resource as $\phi_R$, which is yet another unknown whose value must be determined consistently. Note that $\phi_R = M^*/M$, where $M^*$ is the number of surviving resources and $M$ is the number of available resources in the pool.

#### C.5 Step 5: Deriving the self-consistency equations and community-level properties

We have now obtained expressions for the steady-state abundances for the cavity species and resource, namely Eqs. (54) and (75). These equations suggest that the cavity species and resource abundances follow truncated normal distributions. Now, we invoke self-averaging [3, 5]: since the cavity species and resource are statistically similar to the resident species and resources in the community, respectively, we expect the steady-state abundance distributions of the cavity species and resource to be identical to those of the resident species and resources, respectively. However, note that these equations still contain unknowns that we must solve for, namely:

- $\phi_N$ and $\phi_R$, the survival probability of species and resources, respectively
- $\langle N \rangle$ and $\langle R \rangle$, the average species and resource abundances, respectively
- $\langle N^2 \rangle$ and $\langle R^2 \rangle$, the second moments the in species and resource abundances (which represent their fluctuations)
- $v^{(N)}$ and $\chi^{(R)}$, which describe the average trace of the susceptibilities: the mean change in a species's and resource's steady-state abundance upon a small perturbation in their own death and supply rate, respectively.

We thus have 8 unknowns, and must obtain 8 equations to solve for them in a self-consistent manner. To obtain these 8 equations, note that several of these unknowns are the moments of the species and resource abundance distributions truncated at 0 abundance (which represents extinct species and resources): the zeroth moments represent the survival probability of species ($\phi_N$) and resources ($\phi_R$), the first moments represent the average abundances of surviving species $\langle N\rangle$ and resources $\langle R\rangle$, and similarly for the second moments $\langle N^2\rangle$ and $\langle R^2\rangle$. To solve for them, we To solve for them under the ansatz that the abundances follow truncated normal distributions, we can exploit mathematical identities for these moments. Specifically, the $j^{\text{th}}$ moment of a Gaussian distribution with mean $\mu$ and variance $\sigma^2$ truncated at 0, denoted as $w_j(\mu,\sigma)$, is

$$w_j(\mu,\sigma) = \left\langle x^j \Big( \underbrace{\max\{0, \mu + \sigma Z_x\}}_{\substack{\text{(Note this expression has}\\ \text{the same form as Eqs. 54 and 75)}}} \Big) \right\rangle = \int_{0^+}^{\infty} x^j \frac{1}{\sqrt{2\pi\sigma^2}} \exp\left(-\frac{(x-\mu)^2}{2\sigma^2}\right) dx \tag{76}$$

$$= \sigma^j \int_{-\frac{\mu}{\sigma}}^{\infty} \frac{1}{2\pi} \exp\left(-\frac{z^2}{2}\right) \left(z + \frac{\mu}{\sigma}\right)^j dz. \tag{77}$$

This function is related to the error function and is therefore numerically fast to compute, which will eventually help us solve these equations. Following this, the $j^{th}$ moments of the species and resource abundance distributions are, respectively,

$$\begin{aligned} w_j(\mu_{g_N}, \sigma_{g_N}) &= \left(\frac{\sigma_{g_N}}{\mu_y \sigma_c^2 \chi^{(R)}}\right)^j \int_{-\triangle g_N}^{\infty} \frac{1}{\sqrt{2\pi}} \exp\left(-\frac{z^2}{2}\right) (z + \triangle\, g_N)^j \, dz, \\ w_j(\mu_{g_R}, \sigma_{g_R}) &= \left(\frac{\sigma_{g_R}}{1 - \mu_y \sigma_c^2 \gamma^{-1} v^{(N)}}\right)^j \int_{-\triangle g_R}^{\infty} \frac{1}{\sqrt{2\pi}} \exp\left(-\frac{z^2}{2}\right) (z + \triangle\, g_R)^j \, dz, \\ &\text{where } \triangle\, g_N = \frac{\mu_{g_N}}{\sigma_{g_N}} \text{ and } \triangle\, g_R = \frac{\mu_{g_R}}{\sigma_{g_R}}. \end{aligned} \tag{78}$$

#### C.5.1 The statistical moments of the species abundance distribution

The 0th moment of the species abundance distribution post truncation at 0, i.e., the species survival probability ($\phi_N = S^*/S$), is

$$w_0(\mu_{g_N}, \sigma_{g_N}) = \phi_N \text{ or } \frac{S^*}{S} = \int_{-\triangle g_N}^{\infty} \frac{1}{\sqrt{2\pi}} \exp\left(-\frac{z^2}{2}\right) dz = \Phi(\triangle\, g_N), \tag{79}$$

where

$$\Phi(\triangle\, g_N) = \frac{1}{2}\left(1 + \underset{\substack{\uparrow\\ \text{Gaussian}\\ \text{error}\\ \text{function}}}{\text{erf}} \left(\frac{\triangle\, g_N}{\sqrt{2}}\right)\right). \tag{80}$$

The 1st moment, or average species abundance ($\langle N\rangle$), is

$$\begin{aligned} w_1(\mu_{g_N}, \sigma_{g_N}) = \langle N\rangle &= \left(\frac{\sigma_{g_N}}{\mu_y \sigma_c^2 \chi^{(R)}}\right) \int_{-\triangle g_N}^{\infty} \frac{1}{\sqrt{2\pi}} \exp\left(-\frac{z^2}{2}\right) (z + \triangle\, g_N) \, dz \\ &= \left(\frac{\sigma_{g_N}}{\mu_y \sigma_c^2 \chi^{(R)}}\right) \left(\frac{1}{\sqrt{2\pi}} \exp\left(-\frac{\triangle\, {g_N}^2}{2}\right) + \triangle\, g_N\, \Phi(\triangle\, g_N)\right). \end{aligned} \tag{81}$$

The 2nd moment, which contributes to the variance in species abundances ($\langle N^2 \rangle$), is

$$\begin{aligned} w_2(\mu_{g_N}, \sigma_{g_N}) = \langle N^2 \rangle &= \left( \frac{\sigma_{g_N}}{\mu_y \sigma_c^2 \chi^{(R)}} \right)^2 \int_{-\triangle g_N}^{\infty} \frac{1}{\sqrt{2\pi}} \exp\left( -\frac{z^2}{2} \right) (z + \triangle\, g_N)^2 \, dz \\ &= \left( \frac{\sigma_{g_N}}{\mu_y \sigma_c^2 \chi^{(R)}} \right)^2 \left( \frac{\triangle\, g_N}{\sqrt{2\pi}} \exp\left( -\frac{\triangle\, {g_N}^2}{2} \right) + \left( 1 + \triangle\, {g_N}^2 \right) \Phi\left( \triangle\, g_N \right) \right). \end{aligned} \tag{82}$$

##### C.5.2 The statistical moments of the resource abundance distribution

The resource survival probability $\phi_R = M^*/M$, is

$$w_0(\mu_{g_R}, \sigma_{g_R}) = \phi_R \text{ or } M^*/M = \Phi(\triangle\, g_R). \tag{83}$$

The average resource abundance $\langle R \rangle$, is

$$w_1(\mu_{g_R}, \sigma_{g_R}) = \langle R \rangle = \left( \frac{\sigma_{g_R}}{1 - \mu_y \sigma_c^2 \gamma^{-1} v^{(N)}} \right) \left( \frac{1}{\sqrt{2\pi}} \exp\left( -\frac{\triangle\, {g_R}^2}{2} \right) + \triangle\, g_R\, \Phi\left( \triangle\, g_R \right) \right). \tag{84}$$

The 2nd moment, which helps contributes to the variance in resource abundances $\langle R^2 \rangle$ is

$$w_2(\mu_{g_R}, \sigma_{g_R}) = \langle R^2 \rangle = \left( \frac{\sigma_{g_R}}{1 - \mu_y \sigma_c^2 \gamma^{-1} v^{(N)}} \right)^2 \left( \frac{\triangle\, g_R}{\sqrt{2\pi}} \exp\left( -\frac{\triangle\, {g_R}^2}{2} \right) + \left( 1 + \triangle\, {g_R}^2 \right) \Phi\left( \triangle\, g_R \right) \right). \tag{85}$$

##### C.5.3 The susceptibilities

We can use the above self-consistency equations to solve for the trace of the susceptibilities, $v^{(N)}$ and $\chi^{(R)}$.

Starting with $v^{(N)}$, we remind ourselves that $v^{(N)} = \langle v_{ii}^{(N)} \rangle = \langle \partial N_i / m_i \rangle$. Since by self-averaging, the distribution of $N_0$ is identical to the distribution of $N_i$, we can solve for $\langle \partial N_0 / m_0 \rangle$ to solve for $\langle \partial N_i / m_i \rangle$.

$$v^{(N)} = \left\langle \frac{\partial N_i}{\partial d_i} \right\rangle = \left\langle \frac{\partial N_0}{\partial d_0} \right\rangle. \tag{86}$$

We substitute $N_0$ in Eq. (86) with the distribution of the cavity species abundance derived in eq. (54). Recall that the the probability that $N_0 > 0$ is $\phi_N$ — the species survival probability — and the probability it is extinct is $1 - \phi_N$. This gives us the following expression for $v^{(N)}$:

$$\begin{aligned} v^{(N)} &= \frac{\partial}{\partial d_0} \left\langle \max\left\{ 0, \frac{\mu_{g_N} + \sigma_{g_N} Z_N}{\mu_y \sigma_c^2 \chi^{(R)}} \right\} \right\rangle = \frac{\partial}{\partial d_0} \left\langle (1 - \phi_N)(0) + \phi_N \left( \frac{\mu_{g_N} + \sigma_{g_N} Z_N}{\mu_y \sigma_c^2 \chi^{(R)}} \right) \right\rangle \\ &= \frac{\partial}{\partial d_0} \left\langle \phi_N \left( \frac{\mu_{g_N} + \sigma_{g_N} Z_N}{\mu_y \sigma_c^2 \chi^{(R)}} \right) \right\rangle. \end{aligned} \tag{87}$$

We then invoke the chain rule to transform $\partial / \partial d_0$ in terms of $\mu_d$ — $\partial / \partial d_0 = (\partial / \partial \mu_d) \times (\partial \mu_d / \partial d_0)$.

$$
\begin{aligned}
\frac{\partial}{\partial d_0}\left\langle \phi_N\left(\frac{\mu_{g_N}+\sigma_{g_N}Z_N}{\mu_y\sigma_c^2\chi^{(R)}}\right)\right\rangle &= \frac{\partial}{\partial \mu_d}\cdot\cancelto{1}{\frac{\partial\mu_d}{\partial d_0}}\left\langle \phi_N\left(\frac{\mu_{g_N}+\sigma_{g_N}Z_N}{\mu_y\sigma_c^2\chi^{(R)}}\right)\right\rangle = \frac{\partial}{\partial \mu_d}\left\langle \phi_N\frac{\overbrace{\cancelto{0}{\mu_y\mu_c\langle R\rangle}-\cancelto{1}{\mu_d}}^{\mu_{g_N}}+\cancelto{0}{\sigma_{g_N}Z_N}}{\mu_y\sigma_c^2\chi^{(R)}}\right\rangle \\
&= -\frac{\phi_N}{\mu_y\sigma_c^2\chi^{(R)}}.
\end{aligned}
\tag{88}
$$

Therefore, the expression for $v^{(N)}$ is

$$
v^{(N)} = -\frac{\phi_N}{\mu_y\sigma_c^2\chi^{(R)}}. \tag{89}
$$

Repeating the process with $\chi^{(R)}$ gives us

$$
\chi^{(R)} = \left\langle\frac{\partial R_\alpha}{\partial b_\alpha}\right\rangle = \left\langle\frac{\partial R_0}{\partial b_0}\right\rangle = \frac{\partial}{\partial b_0}\left\langle \max\left\{0, \frac{\mu_{g_R}+\sigma_{g_R}Z_R}{1-\mu_y\sigma_c^2\gamma^{-1}v^{(N)}}\right\}\right\rangle. \tag{90}
$$

Recalling that $\phi_R$ is the resource survival probability,

$$
\begin{aligned}
\frac{\partial}{\partial b_0}\left\langle \max\left\{0, \frac{\mu_{g_R}+\sigma_{g_R}Z_R}{1-\mu_y\sigma_c^2\gamma^{-1}v^{(N)}}\right\}\right\rangle &= \frac{\partial}{\partial b_0}\left\langle (1-\phi_R)(0)+\phi_R\left(\frac{\mu_{g_R}+\sigma_{g_R}Z_R}{1-\mu_y\sigma_c^2\gamma^{-1}v^{(N)}}\right)\right\rangle \\
&= \frac{\partial}{\partial b_0}\left\langle \phi_R\left(\frac{\mu_{g_R}+\sigma_{g_R}Z_R}{1-\mu_y\sigma_c^2\gamma^{-1}v^{(N)}}\right)\right\rangle.
\end{aligned}
\tag{91}
$$

$$
\begin{aligned}
\frac{\partial}{\partial b_0}\left\langle \phi_R\left(\frac{\mu_{g_R}+\sigma_{g_R}Z_R}{1-\mu_y\sigma_c^2\gamma^{-1}v^{(N)}}\right)\right\rangle &= \frac{\partial}{\partial \mu_b}\cancelto{1}{\frac{\partial\mu_b}{\partial b_0}}\left\langle \phi_R\left(\frac{\mu_{g_R}+\sigma_{g_R}Z_R}{1-\mu_y\sigma_c^2\gamma^{-1}v^{(N)}}\right)\right\rangle \\
&= \frac{\partial}{\partial \mu_b}\left\langle \phi_R\left(\frac{\overbrace{\cancelto{1}{\mu_b}-\cancelto{0}{\mu_e\gamma^{-1}\langle N\rangle}}^{\mu_{g_R}}+\cancelto{0}{\sigma_{g_R}Z_R}}{1-\mu_y\sigma_c^2\gamma^{-1}v^{(N)}}\right)\right\rangle \\
&= \frac{\phi_R}{1-\mu_y\sigma_c^2\gamma^{-1}v^{(N)}}.
\end{aligned}
\tag{92}
$$

The expressions for $v^{(N)}$ in eq. (89) and $\chi^{(R)}$ in eq. (92) implicitly involve each other. Therefore, we can solve these equations simultaneously to obtain expressions for $v^{(N)}$ and $\chi^{(R)}$.

$$
v^{(N)} = -\frac{\phi_N}{\mu_y\sigma_c^2(\phi_R-\phi_N\gamma^{-1})}, \qquad \chi^{(R)} = \phi_R-\phi_N\gamma^{-1}. \tag{93}
$$

We have now obtained all 8 equations required to solve the 8 unknowns. Below, we collect all these equations together.

### Final set of self-consistency equations

**The species abundance distribution**

$$N_0 = \max\left\{0, \frac{\mu_{g_N} + \sigma_{g_N} Z_N}{\mu_y \sigma_c^2 \chi^{(R)}}\right\}, \tag{94}$$

$$\phi_N \text{ i.e., } \frac{S^*}{S} = \Phi(\triangle\, g_N), \tag{95}$$

$$\langle N\rangle = \left(\frac{\sigma_{g_N}}{\mu_y \sigma_c^2 \chi^{(R)}}\right)\left(\frac{1}{\sqrt{2\pi}}\exp\left(-\frac{\triangle\, {g_N}^2}{2}\right) + \triangle\, g_N\ \Phi\left(\triangle\, g_N\right)\right), \tag{96}$$

$$\langle N^2\rangle = \left(\frac{\sigma_{g_N}}{\mu_y \sigma_c^2 \chi^{(R)}}\right)^2\left(\frac{\triangle\, g_N}{\sqrt{2\pi}}\exp\left(-\frac{\triangle\, {g_N}^2}{2}\right) + \left(1 + \triangle\, {g_N}^2\right)\Phi\left(\triangle\, g_N\right)\right), \tag{97}$$

$$\text{where } \mu_{g_N} = \mu_y \mu_c \langle R\rangle - \mu_d, \quad \sigma_{g_N} = \sqrt{\left(\sigma_c^2(\mu_y^2 + \sigma_y^2) + \frac{(\mu_c \sigma_y)^2}{M}\right)\langle R^2\rangle + \sigma_d^2} \tag{98}$$

$$\text{and } \triangle\, g_N = \frac{\mu_{g_N}}{\sigma_{g_N}}. \tag{99}$$

**The resource abundance distribution**

$$R_0 = \max\left\{0, \frac{\mu_{g_R} + \sigma_{g_R} Z_R}{1 - \mu_y \sigma_c^2 \gamma^{-1} v^{(N)}}\right\}, \tag{100}$$

$$\phi_R \text{ i.e., } \frac{M^*}{M} = \Phi(\triangle\, g_R), \tag{101}$$

$$\langle R\rangle = \left(\frac{\sigma_{g_R}}{1 - \mu_y \sigma_c^2 \gamma^{-1} v^{(N)}}\right)\left(\frac{1}{\sqrt{2\pi}}\exp\left(-\frac{\triangle\, {g_R}^2}{2}\right) + \triangle\, g_R\ \Phi\left(\triangle\, g_R\right)\right), \tag{102}$$

$$\langle R^2\rangle = \left(\frac{\sigma_{g_R}}{1 - \mu_y \sigma_c^2 \gamma^{-1} v^{(N)}}\right)^2\left(\frac{\triangle\, g_R}{\sqrt{2\pi}}\exp\left(-\frac{\triangle\, {g_R}^2}{2}\right) + \left(1 + \triangle\, {g_R}^2\right)\Phi\left(\triangle\, g_R\right)\right), \tag{103}$$

$$\text{where } \mu_{g_R} = \mu_b - \mu_c \gamma^{-1}\langle N\rangle, \quad \sigma_{g_R} = \sqrt{\sigma_b^2 + \sigma_c^2 \gamma^{-1}\langle N^2\rangle} \quad \text{and } \triangle\, g_R = \frac{\mu_{g_R}}{\sigma_{g_R}}. \tag{104}$$

**The susceptibilities**

$$v^{(N)} = -\frac{\phi_N}{\mu_y \sigma_c^2(\phi_R - \phi_N \gamma^{-1})}, \tag{105}$$

$$\chi^{(R)} = \phi_R - \phi_N \gamma^{-1}. \tag{106}$$

These equations can be solved numerically across given a set of ensemble parameters from (Eqs. (2)–(6)). See Appendix D.3 for details of our numerical solution procedure.

#### C.6 Step 6: Deriving the stability condition

Having derived properties of typical steady-state communities assembled from a pool of $S$ species and $M$ resources, we now proceed to derive when we expect these steady-states to be dynamically stable. For this, we will assess the sensitivity of the community steady-state to small perturbations in the abundances of all surviving species and resources at steady-state. Since the communities are assembled using randomly drawn parameters, the sensitivities of the abundances of different species and resources will also be random variables. Thus, to characterise these sensitivities, we will compute their first two moments: the mean (first moment) and fluctuations (second moment). When either of these moments diverges the sensitivities cease to be well-defined and the community becomes unstable. We will find that it is the second moment that is informative of stability, not the first. The stability boundary thus corresponds to the condition for this second moment to diverge.

We start with eq. (53) describing cavity species dynamics, written explicitly to include fluctuations in growth rate (but not the average growth rate, as we assume the perturbation will not change the statistical properties of species and resource abundances). This is given by

$$0 = N_0\left(\mu_{g_N} + -\sigma_d z_{d,0} + \left(\frac{\mu_y\sigma_c}{\sqrt{M}}\sum_{\alpha=1}^{M} z_{c,0\alpha}R_{\alpha\backslash 0} + \frac{\mu_c\sigma_y}{M}\sum_{\alpha=1}^{M} z_{y,0\alpha}R_{\alpha\backslash 0} + \frac{\sigma_y\sigma_c}{\sqrt{M}}\sum_{\alpha=1}^{M} z_{y,0\alpha}z_{c,0\alpha}R_{\alpha\backslash 0}\right) - \mu_y\sigma_c^2\chi^{(R)}N_0\right). \tag{107}$$

Since this steady state is uninvadable, the community should will be stable to invasion by extinct species and resources. The only source of instability could be when we perturb the abundances of the surviving species and resources. We will thus do so for all surviving species $N^+_{i\backslash 0}$ and resources $R^+_{\alpha\backslash 0}$; here the + indicates survivors. Thus, we may write the abundance of a surviving cavity species $N_0^+$ as:

$$N_0^+ = \frac{1}{\mu_y\sigma_c^2\chi^{(R)}}\left(\mu_{g_N} - \sigma_d z_{d,0} + \frac{\mu_y\sigma_c}{\sqrt{M}}\sum_{\substack{\alpha=1,\\ R^+_{\alpha\backslash 0}>0}}^{M} z_{c,0\alpha}R^+_{\alpha\backslash 0} + \frac{\mu_c\sigma_y}{M}\sum_{\substack{\alpha=1,\\ R^+_{\alpha\backslash 0}>0}}^{M} z_{y,0\alpha}R^+_{\alpha\backslash 0} + \frac{\sigma_y\sigma_c}{\sqrt{M}}\sum_{\substack{\alpha=1,\\ R^+_{\alpha\backslash 0}>0}}^{M} z_{y,0\alpha}z_{c,0\alpha}R^+_{\alpha\backslash 0}\right). \tag{108}$$

We now perturb the abundances of all surviving resources $R^+_{\alpha\backslash 0}$ by a random small amount $\varepsilon\eta_\alpha$ to the surviving resources, where $\varepsilon \ll 1$ sets the scale of the perturbation and keeps it small, and $\eta_\alpha \sim \mathcal{N}(0,1)$ is a standard normal variable indicating that the perturbations are symmetric around 0 and Gaussian-distributed. Thus, the surviving resource abundances become $R^+_{\alpha\backslash 0} \to R^+_{\alpha\backslash 0} + \varepsilon\eta_\alpha$. After the perturbation, the steady-state condition for the abundance of the surviving cavity species is given by

$$\begin{aligned} N_0^+ = \frac{1}{\mu_y\sigma_c^2\chi^{(R)}}\Bigg(&\mu_{g_N} - \sigma_d z_{d,0} + \frac{\mu_y\sigma_c}{\sqrt{M}}\sum_{\substack{\alpha=1,\\ R^+_{\alpha\backslash 0}>0}}^{M} z_{c,0\alpha}\left(R^+_{\alpha\backslash 0} + \varepsilon\eta_\alpha\right) + \frac{\mu_c\sigma_y}{M}\sum_{\substack{\alpha=1,\\ R^+_{\alpha\backslash 0}>0}}^{M} z_{y,0\alpha}\left(R^+_{\alpha\backslash 0} + \varepsilon\eta_\alpha\right) \\ &+ \frac{\sigma_y\sigma_c}{\sqrt{M}}\sum_{\substack{\alpha=1,\\ R^+_{\alpha\backslash 0}>0}}^{M} z_{y,0\alpha}z_{c,0\alpha}\left(R^+_{\alpha\backslash 0} + \varepsilon\eta_\alpha\right)\Bigg). \end{aligned} \tag{109}$$

Next, we compute the sensitivity of this species $\frac{dN_0^+}{d\varepsilon}$ by taking its derivative with respect to the perturbation $\varepsilon$, given by:

$$
\frac{dN_0^+}{d\varepsilon} = \frac{1}{\mu_y \sigma_c^2 \chi^{(R)}} \left( \frac{\mu_y \sigma_c}{\sqrt{M}} \sum_{\substack{\alpha=1,\\ R^+_{\alpha\backslash 0}>0}}^{M} z_{c,0\alpha} \left( \frac{\mathrm{d}R^+_{\alpha\backslash 0}}{\mathrm{d}\varepsilon} + \eta_\alpha \right) + \frac{\mu_c \sigma_y}{M} \sum_{\substack{\alpha=1,\\ R^+_{\alpha\backslash 0}>0}}^{M} z_{y,0\alpha} \left( \frac{\mathrm{d}R^+_{\alpha\backslash 0}}{\mathrm{d}\varepsilon} + \eta_\alpha \right) \right.
$$
$$
\left. + \frac{\sigma_y \sigma_c}{\sqrt{M}} \sum_{\substack{\alpha=1,\\ R^+_{\alpha\backslash 0}>0}}^{M} z_{y,0\alpha} z_{c,0\alpha} \left( \frac{\mathrm{d}R^+_{\alpha\backslash 0}}{\mathrm{d}\varepsilon} + \eta_\alpha \right) \right). \tag{110}
$$

To determine stability, we now compute its first and second moments. The first moment $\langle dN_0^+/d\varepsilon \rangle$, is always finite and 0. To see this, note that:

$$
\left\langle \frac{dN_0^+}{d\varepsilon} \right\rangle = \frac{1}{\mu_y \sigma_c^2 \chi^{(R)}} \left( \frac{\mu_y \sigma_c}{\sqrt{M}} \sum_{\substack{\alpha=1,\\ R^+_{\alpha\backslash 0}>0}}^{M} \cancelto{0}{\langle z_{c,0\alpha} \rangle} \left( \left\langle \frac{\mathrm{d}R^+_{\alpha\backslash 0}}{\mathrm{d}\varepsilon} \right\rangle + \cancelto{0}{\langle \eta_\alpha \rangle} \right) + \frac{\mu_c \sigma_y}{M} \sum_{\substack{\alpha=1,\\ R^+_{\alpha\backslash 0}>0}}^{M} \cancelto{0}{\langle z_{y,0\alpha} \rangle} \left( \left\langle \frac{\mathrm{d}R^+_{\alpha\backslash 0}}{\mathrm{d}\varepsilon} \right\rangle + \cancelto{0}{\langle \eta_\alpha \rangle} \right) \right.
$$
$$
\left. + \frac{\sigma_y \sigma_c}{\sqrt{M}} \sum_{\substack{\alpha=1,\\ R^+_{\alpha\backslash 0}>0}}^{M} \cancelto{0}{\langle z_{y,0\alpha} z_{c,0\alpha} \rangle} \left( \left\langle \frac{\mathrm{d}R^+_{\alpha\backslash 0}}{\mathrm{d}\varepsilon} \right\rangle + \cancelto{0}{\langle \eta_\alpha \rangle} \right) \right)
$$
$$
= 0.
$$

This is expected due to the symmetric nature of the perturbations. We now compute the second moment $\langle (dN_0^+/d\varepsilon)^2 \rangle$, given by:

$$
\left\langle \left( \frac{\mathrm{d}N_0^+}{\mathrm{d}\varepsilon} \right)^2 \right\rangle = \left( \frac{1}{\mu_y \sigma_c^2 \chi^{(R)}} \right)^2 \left\langle \frac{(\mu_y \sigma_c)^2}{M} \left( \sum_{\substack{\alpha=1,\\ R^+_{\alpha\backslash 0}>0}}^{M} z_{c,0\alpha} \left( \frac{\mathrm{d}R^+_{\alpha\backslash 0}}{\mathrm{d}\varepsilon} + \eta_\alpha \right) \right)^2 + \frac{(\mu_c \sigma_y)^2}{M^2} \left( \sum_{\substack{\alpha=1,\\ R^+_{\alpha\backslash 0}>0}}^{M} z_{y,0\alpha} \left( \frac{\mathrm{d}R^+_{\alpha\backslash 0}}{\mathrm{d}\varepsilon} + \eta_\alpha \right) \right)^2 \right.
$$
$$
+ \frac{(\sigma_y \sigma_c)^2}{M} \left( \sum_{\substack{\alpha=1,\\ R^+_{\alpha\backslash 0}>0}}^{M} z_{y,0\alpha} z_{c,0\alpha} \left( \frac{\mathrm{d}R^+_{\alpha\backslash 0}}{\mathrm{d}\varepsilon} + \eta_\alpha \right) \right)^2
$$
$$
+ \frac{2\mu_c \mu_y \sigma_y \sigma_c}{M^{3/2}} \left( \sum_{\substack{\alpha=1,\\ R^+_{\alpha\backslash 0}>0}}^{M} z_{c,0\alpha} \left( \frac{\mathrm{d}R^+_{\alpha\backslash 0}}{\mathrm{d}\varepsilon} + \eta_\alpha \right) \right) \left( \sum_{\substack{\alpha=1,\\ R^+_{\alpha\backslash 0}>0}}^{M} z_{y,0\alpha} \left( \frac{\mathrm{d}R^+_{\alpha\backslash 0}}{\mathrm{d}\varepsilon} + \eta_\alpha \right) \right)
$$
$$
+ \frac{2\mu_y \sigma_y \sigma_c^2}{M} \left( \sum_{\substack{\alpha=1,\\ R^+_{\alpha\backslash 0}>0}}^{M} z_{c,0\alpha} \left( \frac{\mathrm{d}R^+_{\alpha\backslash 0}}{\mathrm{d}\varepsilon} + \eta_\alpha \right) \right) \left( \sum_{\substack{\alpha=1,\\ R^+_{\alpha\backslash 0}>0}}^{M} z_{c,0\alpha}^2 z_{y,0\alpha} \left( \frac{\mathrm{d}R^+_{\alpha\backslash 0}}{\mathrm{d}\varepsilon} + \eta_\alpha \right) \right)
$$
$$
\left. + \frac{2\mu_c \sigma_y \sigma_c^2}{M^{3/2}} \left( \sum_{\substack{\alpha=1,\\ R^+_{\alpha\backslash 0}>0}}^{M} z_{y,0\alpha} \left( \frac{\mathrm{d}R^+_{\alpha\backslash 0}}{\mathrm{d}\varepsilon} + \eta_\alpha \right) \right) \left( \sum_{\substack{\alpha=1,\\ R^+_{\alpha\backslash 0}>0}}^{M} z_{y,0\alpha} z_{c,0\alpha}^2 \left( \frac{\mathrm{d}R^+_{\alpha\backslash 0}}{\mathrm{d}\varepsilon} + \eta_\alpha \right) \right) \right\rangle. \tag{111}
$$

This expression reduces to

$$
\left\langle \left( \frac{\mathrm{d}N_0^+}{\mathrm{d}\varepsilon} \right)^2 \right\rangle = \frac{\phi_R \left( \sigma_c^2 (\mu_y^2 + \sigma_y^2) + (\mu_c \sigma_y)^2/M \right)}{\left( \mu_y \sigma_c^2 \chi^{(R)} \right)^2} \left( \left\langle \left( \frac{\mathrm{d}R^+_{\alpha\backslash 0}}{\mathrm{d}\varepsilon} \right)^2 \right\rangle + 1 \right). \tag{112}
$$

Since the cavity resource is statistically identical to other communities members, we can replace $\mathrm{d}R^+_{\alpha\setminus 0}/\mathrm{d}\varepsilon$ with $\mathrm{d}R^+_0/\mathrm{d}\varepsilon$. Therefore, the sensitivity of the cavity species to perturbation is given by

$$\left\langle\left(\frac{\mathrm{d}N^+_0}{\mathrm{d}\varepsilon}\right)^2\right\rangle = \frac{\phi_R\left(\sigma_c^2(\mu_y^2+\sigma_y^2)+(\mu_c\sigma_y)^2/M\right)}{\left(\mu_y\sigma_c^2\chi^{(R)}\right)^2}\left(\left\langle\left(\frac{\mathrm{d}R^+_0}{\mathrm{d}\varepsilon}\right)^2\right\rangle+1\right). \tag{113}$$

Repeating this process with the cavity resource, we get:

$$R^+_0 = \frac{1}{1-\mu_y\sigma_c^2\gamma^{-1}v^{(N)}}\left(\mu_{g_R}+\sigma_b z_{b,0}+\frac{\sigma_c}{\sqrt{M}}\sum_{i=1}^{S} z_{c,i0}N^+_{i\setminus 0}\right). \tag{114}$$

$$\frac{\mathrm{d}R^+_0}{\mathrm{d}\varepsilon} = \frac{1}{1-\mu_g\sigma_c^2\gamma^{-1}v^{(N)}}\left(\frac{\sigma_c}{\sqrt{M}}\sum_{i=1}^{S} z_{c,i0}\left(\frac{\mathrm{d}N^+_{i\setminus 0}}{\mathrm{d}\varepsilon}+\eta_i\right)\right). \tag{115}$$

$$\begin{aligned}\left\langle\left(\frac{\mathrm{d}R^+_0}{\mathrm{d}\varepsilon}\right)^2\right\rangle &= \frac{\sigma_c^2}{M(1-\mu_y\sigma_c^2\gamma^{-1}v^{(N)})^2}\left\langle\left(\sum_{i=1}^{S} z_{c,i0}\left(\frac{\mathrm{d}N^+_{i\setminus 0}}{\mathrm{d}\varepsilon}+\eta_i\right)\right)^2\right\rangle\\ &= \frac{\sigma_c^2\gamma^{-1}\phi_N}{(1-\mu_y\sigma_c^2\gamma^{-1}v^{(N)})^2}\left(\left\langle\left(\frac{\mathrm{d}N^+_{i\setminus 0}}{\mathrm{d}\varepsilon}\right)^2\right\rangle+1\right),\end{aligned} \tag{116}$$

or equivalently, by invoking the statistical identity of the cavity species and other residents, we get:

$$\left\langle\left(\frac{\mathrm{d}R^+_0}{\mathrm{d}\varepsilon}\right)^2\right\rangle = \frac{\sigma_c^2\gamma^{-1}\phi_N}{(1-\mu_y\sigma_c^2\gamma^{-1}v^{(N)})^2}\left(\left\langle\left(\frac{\mathrm{d}N^+_0}{\mathrm{d}\varepsilon}\right)^2\right\rangle+1\right). \tag{117}$$

Solving for $\left\langle\left(dN^+_0/d\varepsilon\right)^2\right\rangle$ and $\left\langle\left(dR^+_0/d\varepsilon\right)^2\right\rangle$ gives us the following expressions

$$\begin{aligned}&\left\langle\left(\frac{\mathrm{d}N^+_0}{\mathrm{d}\varepsilon}\right)^2\right\rangle = \frac{B+1}{AB-1}, \qquad \left\langle\left(\frac{\mathrm{d}R^+_0}{\mathrm{d}\varepsilon}\right)^2\right\rangle = \frac{A+1}{AB-1},\\ &\text{where } \frac{1}{A} = \frac{\phi_R\left(\sigma_c^2(\mu_y^2+\sigma_y^2)+(\mu_c\sigma_y)^2/M\right)}{\left(\mu_y\sigma_c^2\chi^{(R)}\right)^2} \text{ and } \frac{1}{B} = \frac{\sigma_c^2\gamma^{-1}\phi_N}{(1-\mu_y\sigma_c^2\gamma^{-1}v^{(N)})^2}.\end{aligned} \tag{118}$$

Communities undergo a transition from stability to instability when these second moments of the sensitivities diverge. This occurs when $AB-1=0$:

$$\frac{\left(\mu_y\sigma_c^2\chi^{(R)}\right)^2\left(1-\mu_y\sigma_c^2\gamma^{-1}v^{(N)}\right)^2}{\phi_R\left(\sigma_c^2(\mu_y^2+\sigma_y^2)+(\mu_c\sigma_y)^2/M\right)\left(\sigma_c^2\gamma^{-1}\phi_N\right)}-1=0. \tag{119}$$

Plugging in the expressions for $v^{(N)}$ and $\chi^{(R)}$ from eq. (93) and rearranging this equation simplifies it to

$$\overbrace{\frac{\phi_N \gamma^{-1}}{\phi_R}}^{\text{species packing ratio}} \times \overbrace{\frac{\sigma_c^2(\mu_y^2+\sigma_y^2)+(\mu_c\sigma_y)^2/M}{(\mu_y\sigma_c)^2}}^{1/(\text{interaction reciprocity})^2} -1 = 0. \tag{120}$$

We can see that the stability condition depends on two key properties. First, the species packing ratio, which is the ratio of the number of surviving species to surviving resources ($S^*/M^*$). Here, it is expressed as $\phi_N\gamma^{-1}/\phi_R$, where $\phi_N$ and $\phi_R$ are the species and resource survival probabilities, and $\gamma^{-1} = S/M$ is the ratio of the species and resource pool size. The second term $(\sigma_c^2(\mu_y^2+\sigma_y^2)+(\mu_c\sigma_y)^2/M)/(\mu_y\sigma_c)^2$ is the reciprocal of (interaction reciprocity)$^2$ or the correlation between growth ($g_{i\alpha}$) and consumption ($c_{i\alpha}$) coefficients — see Eq. 11.

Therefore, communities remain stable when

$$\underset{\text{interaction reciprocity}}{\rho_{c,g}} \quad > \quad \underset{\sqrt{\text{species packing ratio}}}{\sqrt{\frac{S^*}{M^*}}}, \quad \text{where } \rho_{c,g} = \sqrt{\frac{(\mu_y\sigma_c)^2}{\sigma_c^2(\mu_y^2+\sigma_y^2)+(\mu_c\sigma_y)^2/M}} = \frac{1}{\sqrt{1+\frac{\sigma_y^2}{\mu_y^2}\left(1+\frac{\mu_c^2}{M\sigma_c^2}\right)}} \tag{121}$$

$$\text{and } \frac{S^*}{M^*} = \frac{\phi_N\gamma^{-1}}{\phi_R} = \frac{\Phi(\triangle\, g_N)}{\Phi(\triangle\, g_R)}.$$

Therefore, communities transition from stability to instability when the $\sqrt{\text{packing ratio}}$ exceeds interaction reciprocity. Our simulations show that when communities cross the stability boundary, their dynamics transition from reaching a unique stable state to persistent fluctuations consistent with chaos (i.e., with a positive maximal Lyapunov exponent; see Appendix D).

#### Stability equations

$$\left\langle \left(\frac{\mathrm{d}N_0^+}{\mathrm{d}\varepsilon}\right)^2 \right\rangle = \frac{\phi_R\left(\sigma_c^2(\mu_y^2+\sigma_y^2)+(\mu_c\sigma_y)^2/M\right)}{(\mu_y\sigma_c^2\chi^{(R)})^2}\left(\left\langle \left(\frac{\mathrm{d}R_0^+}{\mathrm{d}\varepsilon}\right)^2 \right\rangle + 1\right). \tag{122}$$

$$\left\langle \left(\frac{\mathrm{d}R_0^+}{\mathrm{d}\varepsilon}\right)^2 \right\rangle = \frac{\sigma_c^2\phi_N\gamma^{-1}}{\left(1-\mu_y\sigma_c^2\gamma^{-1}v^{(N)}\right)^2}\left(\left\langle \left(\frac{dN_0}{d\varepsilon}\right)^2 \right\rangle + 1\right). \tag{123}$$

$$\textbf{Stability condition: } \rho_{c,g} > \sqrt{\frac{\phi_N\gamma^{-1}}{\phi_R}}. \tag{124}$$

### D The effective Lotka-Volterra model (derived from the consumer-resource model)

A natural question is whether resource dynamics actually drive the stability transitions we observe in consumer-resource models (CRM), or whether they can be explained by resource-mediated competition alone. If the latter is true, a resource-implicit effective Lotka-Volterra model (eLV), which is obtained through eliminating resource dynamics in the CRM, should capture the same transitions.

In eLV models, communities are (often) stable when the strength of self-inhibition ($A_{ii}$). exceeds regulation from inter-species interactions ($A_{ij}$). Therefore, we can determine whether a community is likely to be stable by comparing the statistical properties of inter-species interactions vs self-inhibition. In this section, we derive the eLV from our consumer-resource model and characterize how these statistical properties emerge from the underlying properties of the species and resource pool. We then show multiple examples of where the eLV fails to capture the stabilising effects of resource pool size in the CRM (e.g., over more parameter values, including information on whether resources are extinct or not), supporting our claims in the main text.

#### D.1 The Consumer-Resource model reduces to the generalised Lotka-Volterra model under fast resource dynamics

When resource dynamics are much faster than consumer dynamics, we can argue that resource dynamics are at pseudo-steady state at time $t$. Under this scenario, the resource dynamics in eq. (1) become

$$\frac{dR_\alpha}{dt} = 0 = R_\alpha^{(ss)}\left(b_\alpha - R_\alpha^{(ss)} - \sum_{i=1}^{S} c_{i\alpha} N_i\right), \tag{125}$$

meaning we can write down the expression for resource abundances as

$$R_\alpha^{(ss)} = \max\left\{0,\ b_\alpha - \sum_{i=1}^{S} c_{i\alpha} N_i\right\}. \tag{126}$$

Eq. (126) tells us that at pseudo-steady state, resource abundances depend only on consumer abundances. Substituting the expression for $R_\alpha$ from eq. (126) into the consumer dynamics from eq. (1) gives us

$$\frac{dN_i}{dt} = N_i\left(\sum_{\alpha=1}^{M} y_{i\alpha} c_{i\alpha} R_\alpha^{(ss)} - d_i\right) = N_i\left(\sum_{\alpha=1}^{M} y_{i\alpha} c_{i\alpha}\left(b_\alpha - \sum_{i=1}^{S} c_{i\alpha} N_i\right) \underbrace{\Theta(R_\alpha^{(ss)})}_{\substack{=0 \text{ when } R_\alpha^{(ss)}=0,\\ =1 \text{ otherwise.}}} - d_i\right), \tag{127}$$

where $\Theta(R_\alpha^{(ss)})$ is the Heaviside function and indicates when a resource is present or extinct — it equals 0 when $R_\alpha^{(ss)} = 0$ and 1 otherwise (i.e., when the resource has survived). Therefore, consumer dynamics no longer explicitly depend on resource abundances, only whether they are present or absent. Instead, consumer dynamics only depend explicitly on consumer abundances, so are described by an effective Lotka-Volterra model. This can be seen from rearranging eq. (127):

$$
\begin{aligned}
\frac{\mathrm{d}N_i}{\mathrm{d}t} =& N_i\left(\sum_{\alpha=1}^{M} y_{i\alpha}c_{i\alpha}b_\alpha\Theta(R_\alpha^{(ss)}) - \sum_{\alpha=1}^{M} y_{i\alpha}c_{i\alpha}\sum_{j=1}^{S} c_{j\alpha}N_j\Theta(R_\alpha^{(ss)}) - d_i\right) \\
=& N_i\left(\underbrace{\sum_{\alpha=1}^{M} y_{i\alpha}c_{i\alpha}b_\alpha\Theta(R_\alpha^{(ss)}) - d_i}_{r_i} - \sum_{j=1}^{S}\underbrace{\sum_{\alpha=1}^{M} y_{i\alpha}c_{i\alpha}c_{j\alpha}\Theta(R_\alpha^{(ss)})}_{A_{ij}}N_j\right) \\
=& N_i\left(r_i - \sum_{j=1}^{S} A_{ij}N_j\right),
\end{aligned}
\tag{128}
$$

where $r_i$ is species $i$'s intrinsic growth coefficient, $A_{ij}$ is the competition coefficient, or the of species $j$ on $i$. Finally, we will decompose the species competition term $\sum_{j=1}^{S} A_{ij}N_j$ in self-inhibition and inter-consumer interactions, so the form of the eLV matches the gLV. This gives us the final version of the eLV:

$$
\begin{aligned}
\frac{\mathrm{d}N_i}{\mathrm{d}t} =& N_i\left(r_i - A_{ii}N_i - \sum_{j\neq i}^{S} A_{ij}N_j\right), \\
& \text{where } r_i = \sum_{\alpha=1}^{M} y_{i\alpha}c_{i\alpha}b_\alpha\Theta(R_\alpha^{(ss)}) - d_i, \\
& A_{ii} = \sum_{\alpha=1}^{M} y_{i\alpha}c_{i\alpha}^2\Theta(R_\alpha^{(ss)}), \\
& A_{ij} = \sum_{\alpha=1}^{M} y_{i\alpha}c_{i\alpha}c_{j\alpha}\Theta(R_\alpha^{(ss)}).
\end{aligned}
\tag{129}
$$

Species' growth and competition coefficients depend on their growth and consumption of shared resources. For example, the competition coefficient of species $j$ on $i$ ($A_{ij}$) is sum over all resources of the growth coefficient of species $j$ on resource $\alpha$ $i(y_{i\alpha}c_{i\alpha})$ multiplied by the consumption by $i$ of the same resource.

#### D.2 The statistical properties of inter-consumer interactions and self-inhibition in the eLV

Self-inhibition ($A_{ii}$) and inter-species competition coefficients ($A_{ij}$) consist of sums of weakly-correlated random variables. Therefore, we can describe $A_{ii}$ and $A_{ij}$ each as a Gaussian variable, according to the central limit theorem. To construct these variables, we calculated their means, variances and correlations (detailed in the Extended Information), which are given below:

##### D.2.1 Self-inhibition

Self-inhibition ($A_{ii}$) can be described as the Gaussian variable

$$
A_{ii} = \mu_{A_{ii}} + \sigma_{A_{ii}}z_{D,i},
$$

where

$$
\begin{aligned}
\mu_{A_{ii}} &= \phi_R \left( \frac{\mu_y \mu_c^2}{M} + \mu_y \sigma_c^2 \right), \\
\sigma_{A_{ii}}^2 &= \frac{(\mu_y \sigma_c^2)^2 \phi_R}{M} \left( 3 + \frac{4\mu_c^2}{\sigma_c^2 M} + \frac{\sigma_y^2}{\mu_y^2} \left( 3 + \frac{6\mu_c^2}{\sigma_c^2 M} + \left( \frac{\mu_c^2}{\sigma_c^2 M} \right)^2 \right) \right),
\end{aligned}
$$

and $z_{D,i}$ is a standard normal variable of form

$$
\begin{aligned}
\langle z_{D,i} \rangle &= 0, \\
\langle z_{D,i}^2 \rangle &= 1.
\end{aligned} \tag{130}
$$

##### D.2.2 Inter-species competition coefficient

The inter-species competition coefficient ($A_{ij}$) can be described as the Gaussian variable

$$
A_{ij} = \mu_{A_{ij}} + \sigma_{A_{ij}} z_{A,ij},
$$

where

$$
\begin{aligned}
\mu_{A_{ij}} &= \frac{\mu_y \mu_c^2}{M} \phi_R, \\
\sigma_{A_{ij}}^2 &= \frac{(\mu_y \sigma_c^2)^2 \phi_R}{M} \left( 1 + \frac{2\mu_c^2}{\sigma_c^2 M} + \frac{\sigma_y^2}{\mu_y^2} \left( 1 + \frac{\mu_c^2}{\sigma_c^2 M} \right)^2 \right),
\end{aligned}
$$

and $z_{A,ij}$ is a standard normal variable of form

$$
\begin{aligned}
\langle z_{A,ij} \rangle &= 0, \\
\langle z_{A,ij}^2 \rangle &= 1, \\
\langle z_{A,ij} z_{A,ji} \rangle = \rho_A^{(D)} &= \frac{\left( 1 + \frac{2\mu_c^2}{\sigma_c^2 M} \right)}{1 + \frac{2\mu_c^2}{\sigma_c^2 M} + \frac{\sigma_y^2}{\mu_y^2} \left( 1 + \frac{\mu_c^2}{\sigma_c^2 M} \right)^2}, \\
\langle z_{A,ij} z_{A,ik} \rangle = \rho_A^{(R)} &= \frac{\frac{\mu_c^2}{\sigma_c^2 M} \left( 1 + \frac{\sigma_y^2}{\mu_y^2} \left( 1 + \frac{\mu_c^2}{\sigma_c^2 M} \right) \right)}{1 + \frac{2\mu_c^2}{\sigma_c^2 M} + \frac{\sigma_y^2}{\mu_y^2} \left( 1 + \frac{\mu_c^2}{\sigma_c^2 M} \right)^2}.
\end{aligned} \tag{131}
$$

$\rho_A^{(D)}$ describes the cross-diagonal correlation of inter-species competition coefficients and $\rho_A^{(R)}$ describes the row correlation. Note that $A_{ij}$ also has column correlations $\langle z_{A,ji} z_{A,ki} \rangle$ and transpose row-column correlations $z_{A,ij} z_{A,ki}$, but these appear to be small when $M \gg 1$ and thus are discarded. (See the Extended Information for their calculation.)

### D.3 How properties of the species and resource pool affect inter-species competition and self-inhibition

In this section, we examine how changing the resource pool size ($M$) and sources of interaction heterogeneity ($\sigma_c$ and $\sigma_y$) influence the mean and variance of inter-species competition and self-inhibition. These statistical properties control stability in the generalised Lotka-Volterra model [1–3, 10, 11], so understanding how they may help readers intuit stability transitions in the eLV, and subsequently why they differ to the CRM.

#### D.3.1 Resource pool size

Firstly, we will examine how the resource pool size $M$ affects self-inhibition and inter-species competition. Equations (130) and (131) indicate that increasing $M$ affects all statistical properties of self-inhibition and inter-species competition.

**D.3.1.1 Mean self-inhibition and inter-species competition** Increasing $M$ decreases both the mean self-inhibition ($\mu_{A_{ii}}$) and inter-species competition ($\mu_{A_{ij}}$) at identical rates (Fig. S1 A). This occurs because both means share the same $M$-dependent component — $\mu_y \mu_c^2/M$. However, $\mu_{A_{ii}}$ and $\mu_{A_{ij}}$ also depend indirectly on $M$ through the resource survival fraction ($\phi_R$), which increases with $M$ (Fig. S9). When we account for this effect, the increasing $phi_R$ marginally dampens the direct effect of $M$ on $\mu_{A_{ii}}$ and $\mu_{A_{ij}}$, but the qualitative relationship remains unchanged — increasing $M$ decreases $\mu_{A_{ii}}$ and $\mu_{A_{ij}}$ (Fig. S1 B). As $M \to \infty$, the mean self-inhibition approaches $\mu_y \sigma_c^2$ while inter-species competition approaches 0, but this occurs at values of $M$ beyond those explored in our study. Therefore, within the limit of large but finite $M$, increasing the resource pool size decreases self-inhibition and inter-species competition proportionally.

Some readers may be interested in how the mean total competition a consumer experiences from all other community members experienced is affected by the resource pool size. The mean total competition experienced by a consumer is

$$\mu_{A_{ij}} S^* = \frac{\mu_y \mu_c^2}{M} \phi_R \times \phi_N S = \left(\mu_y \mu_c^2\right) \phi_N \phi_R \gamma^{-1}. \tag{132}$$

Increasing the resource pool size $M$ has no direct effect on th mean total competition (Fig. S1 A), showing that consumers are not weakening competition by, for example, specialising on different resources. In fact, when we account for the indirect effects of $M$ on $\phi_N$ and $\phi_R$, increasing $M$ actually increases the mean total competition experienced by a consumer (Fig. S1 B).

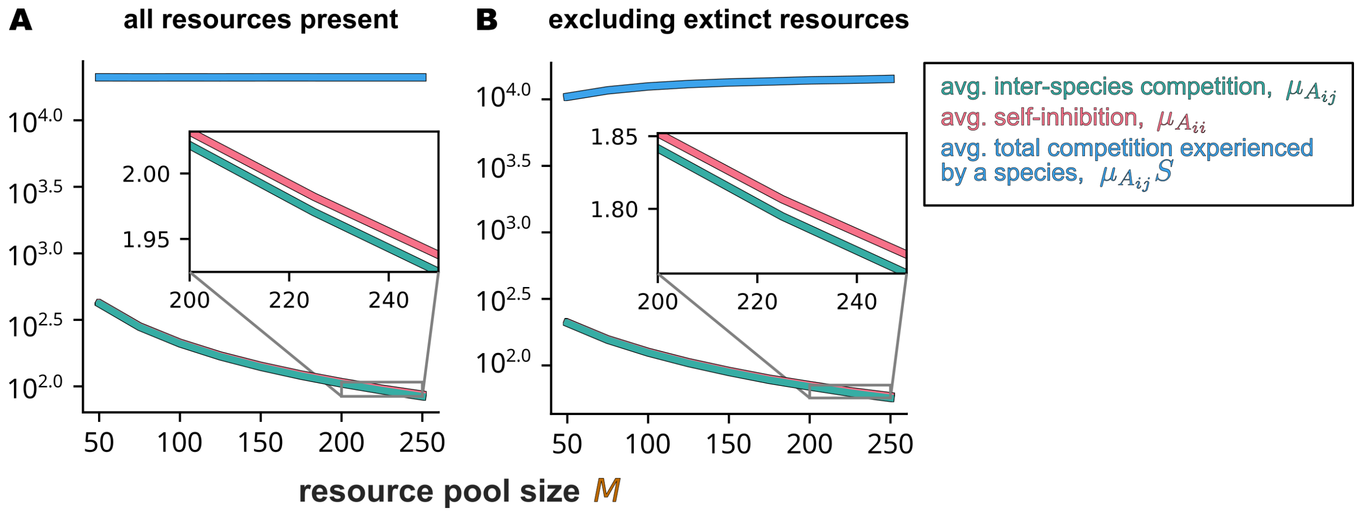

Figure S1: The effect of resource pool size on the average inter-species interaction strength and self-inhibition in the eLV **A** The effect of resource pool size when no resource dynamics are modelled i.e., we assume the resource survival fraction $\phi_R = 1$. All interaction statistics are computed from the interaction matrices of the eLVs, which were constructed using the consumer-resource models in the main text. $\mu_c = 145$, other parameters are default. **B** The effect of resource pool size when resource dynamics i.e., $\phi_R$ and $\phi_N$, are considered.

**D.3.1.2 Standard deviation in self-inhibition and inter-species competition** Moving onto the standard deviations in self-inhibition ($\sigma_{A_{ii}}$) and the inter-species competition coefficient ($\sigma_{A_ij}$), increasing the resource pool size decreases both quantities (Fig. S2 A). Again, the indirect positive effect of $M$ on $\phi_R$ dampens, but does not qualitatively change, the effect of $M$ on these deviations (Fig. S2 B).

The standard deviation in total inter-species competition coefficient experienced by a consumer is

$$\begin{aligned}\sigma_{A_{ij}}\sqrt{S} &= \sqrt{\frac{(\mu_y\sigma_c^2)^2\phi_R}{M}\left(1+\frac{2\mu_c^2}{\sigma_c^2 M}+\frac{\sigma_y^2}{\mu_y^2}\left(1+\frac{\mu_c^2}{\sigma_c^2 M}\right)^2\right)} \times \sqrt{\phi_N S} \\ &= \mu_y\sigma_c^2\sqrt{\phi_N\phi_R\gamma^{-1}\left(1+\frac{2\mu_c^2}{\sigma_c^2 M}+\frac{\sigma_y^2}{\mu_y^2}\left(1+\frac{\mu_c^2}{\sigma_c^2 M}\right)^2\right)}.\end{aligned} \tag{133}$$

This expression shows that increasing $M$ decreases the standard deviation in the inter-species competition (Fig. S2 B).

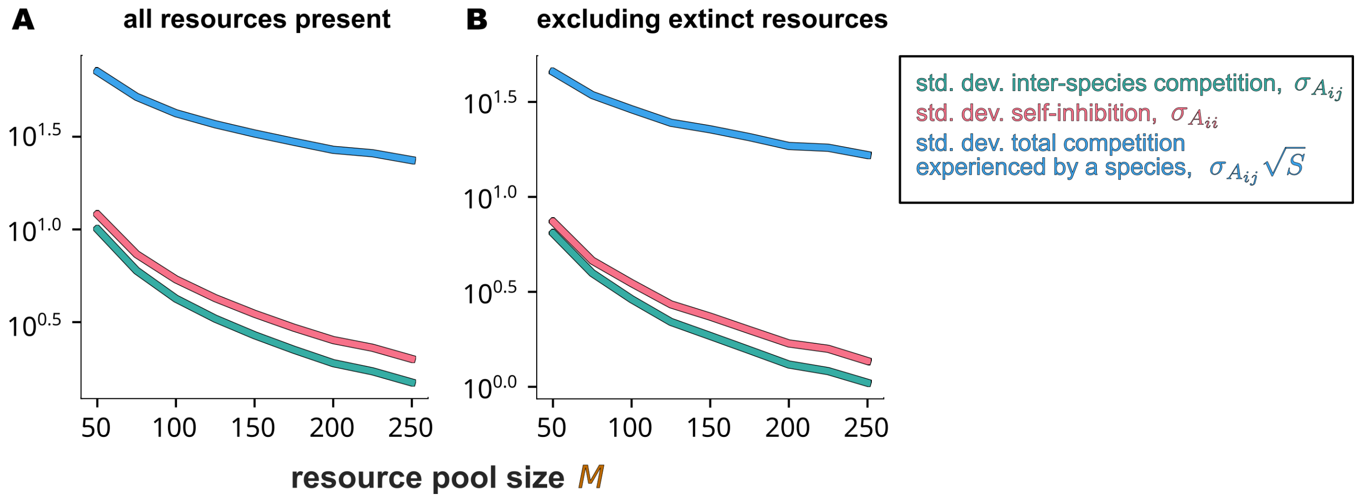

Figure S2: The effect of resource pool size on the standard deviation in inter-species interaction strength and self-inhibition in the eLV **A** The effect of resource pool size when no resource dynamics are modelled i.e., we assume the resource survival fraction $\phi_R = 1$. All interaction statistics are calculated numerically from the interaction matrices of the eLVs, which were constructed using the consumer-resource models in the main text. $\mu_c = 145$, other parameters are default. **B** The effect of resource pool size when resource dynamics i.e., $\phi_R$ and $\phi_N$ are considered.

##### D.3.2 Different sources of interaction heterogeneity

Now, we will look at how different sources of interaction heterogeneity — the standard deviation in the consumption coefficient $\sigma_c$ and the yield conversion $\sigma_y$ — affect self-inhibition and inter-species competition.

**D.3.2.1 Mean self-inhibition and inter-species competition** Increasing the standard deviation in consumption ($\sigma_c$) has no effect on the mean inter-species competition coefficient but increases the mean self-inhibition (Fig. S3 A). In contrast, increasing the standard deviation in yield conversion $\sigma_y$ will not affect either means (Fig. S3 B). (We assume incorporating $\phi_R$ and $\phi_N$ does not change the effect of $\sigma_c$ and $\sigma_y$ on $\mu_{A_{ii}}$ and $\mu_{A_{ij}}$ as these terms have many direct dependencies on the latter quantities.)

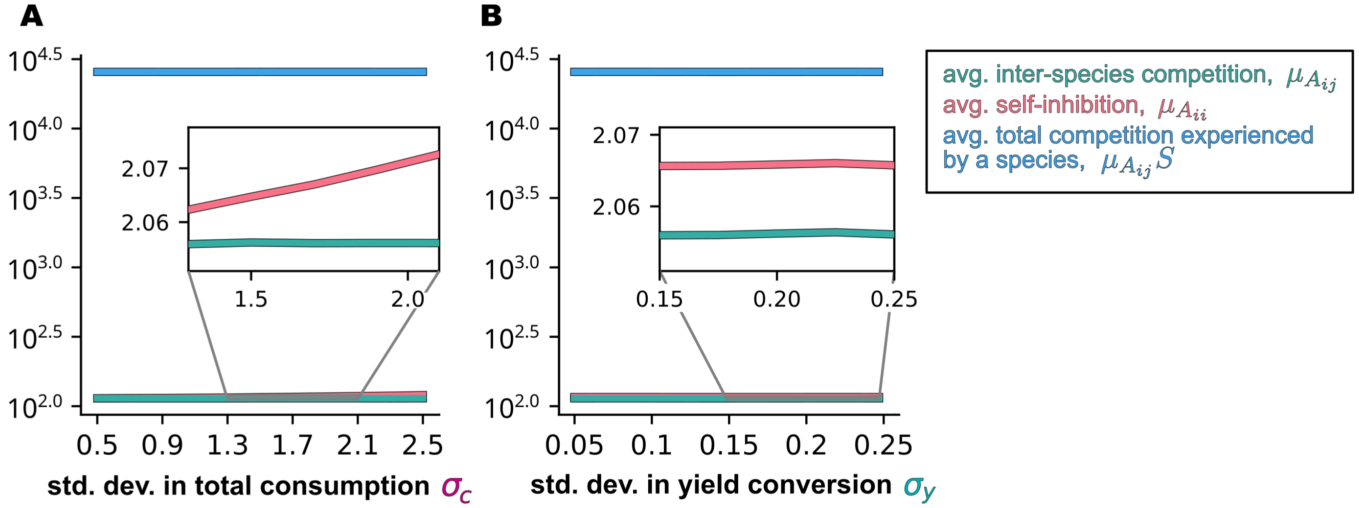

Figure S3: The effect of the standard deviation in the consumption coefficient and yield conversion on the average inter-species interaction strength and self-inhibition in the eLV. **A** The effect of the standard deviation in consumption ($\sigma_c$) when no resource dynamics are modelled i.e., we do not consider which resources survive or are extinct. The effect of variance in the yield conversion factor ($\sigma_y$). The y-axis scale is log base 10. All interaction statistics are calculated numerically from the interaction matrices of the eLVs, which were constructed using the consumer-resource models in the main text. $\mu_c = 160$, other parameters are default.

**D.3.2.2 Standard deviation in self-inhibition and inter-species competition** Moving onto the standard deviation, increasing the standard deviation in the consumption coefficient $\sigma_c$ should increase both the variance in inter-species interactions and self-inhibition (Fig. S4 A). Increasing the standard deviation in yield conversion $\sigma_y$ will likely have a similar effect, as is shown (Fig. S4 B).

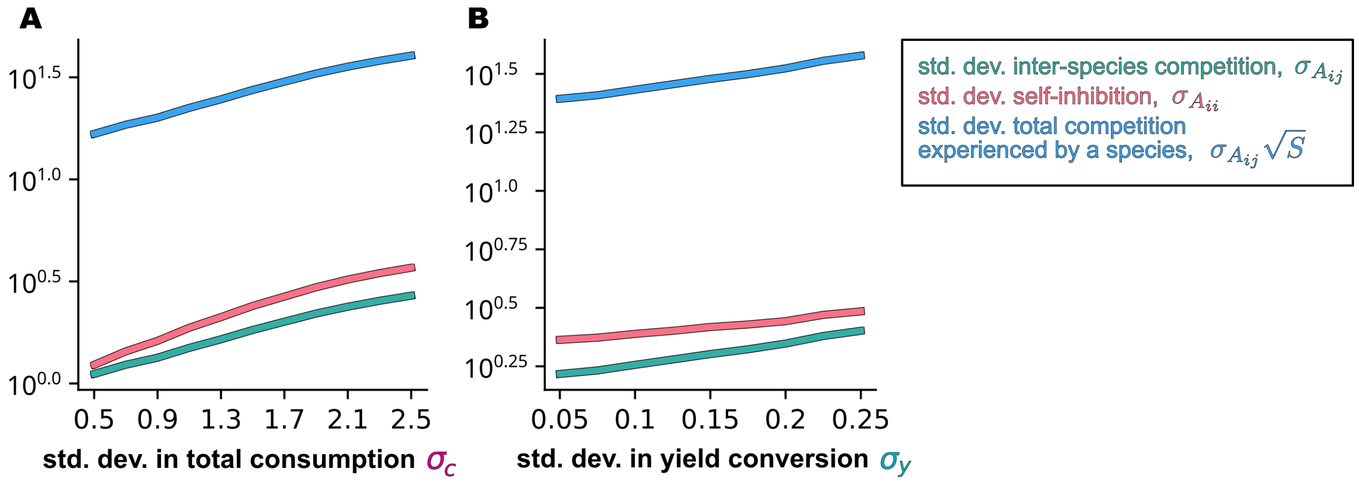

Figure S4: The effect of variance in the consumption coefficient and yield conversion on the variance in inter-species interaction strength and self-inhibition in the eLV. **A** The effect of the standard deviation in consumption ($\sigma_c$) when no resource dynamics are modelled i.e., we assume $\phi_R = 1$. All interaction statistics are calculated numerically from the interaction matrices of the eLVs, which were constructed using the consumer-resource models in the main text. $\mu_c = 160$, other parameters are default. **B** The effect of variance in the yield conversion factor ($\sigma_y$).

#### D.4 The stability condition for the effective Lotka-Volterra model is not equivalent to the consumer-resource model

To determine why the stabilising effect of resource diversity in the CRM is not captured by the eLV, we derived the eLV's stability condition. We did this using the same approach as the CRM. We first determined the statistical properties of typical steady-state communities after community assembly using the cavity method. Because we were not sure that whether the typical steady-state communities in the CRM would be the same as the eLV, we did not include any emergent properties of the CRM, such as the resource survival fraction, in our eLV calculation; **i.e., we assume $\phi_R = 1$.** This should not matter if explicit resource dynamics do not drive the stability transitions in the CRM, as properties of the resource abundance distribution should not meaningfully affect the stability condition.

After deriving the species abundance distribution at steady state, we derived the stability condition by determining when the sensitivity of species abundances to perturbations would diverge to infinity. We found that in the eLV model, communities were stable when

$$\sigma^2_{A_{ij}} S\phi_N < (\mu_{A_{ii}} - \mu_{A_{ij}} - \sigma^2_{A_{ij}} Sv\rho_A^{(D)})^2 + (\sigma_{A_{ii}} - \sigma_{A_{ij}})^2, \tag{134}$$

where $v$ is the average susceptibility of species $i$ to perturbations by the cavity species in the eLV, NOT the CRM. This stability threshold is effectively identical to the threshold for generalised Lotka-Volterra model, as derived by Bunin [3] and Barbier and Arnoldi [6]:

$$\sigma^2_{A_{ij}} \phi_N < (1 - \sigma^2_{A_{ij}} v\rho_A^{(D)})^2. \tag{135}$$

Differences only emerge between stability conditions because Bunin [3] and Barbier and Arnoldi [6] made the following assumptions:

1. $\sigma^2_{A_{ij}} \propto 1/S$, so $\sigma^2_{A_{ij}} S = O(1)$ rather than $O(S)$. *Once we substitute in our value for $\sigma_{A_{ij}}$ derived from the CRM, $\sigma^2_{A_{ij}} S = O(1)$ in our stability condition.*
2. Self-inhibition $A_{ii}$ is fixed to 1 and $A_{ij} \propto 1/S$. Therefore,
   (a) $\mu_{A_{ii}} - \mu_{A_{ij}}$ becomes $1 - \mu_{A_{ij}}/S \approx 1$ when $M$ is very large.
   (b) $(\sigma_{A_{ii}} - \sigma_{A_{ij}})^2$ becomes $(0 - \sigma_{A_{ij}}/\sqrt{S})^2 \approx 0$ when $M$ is very large

Substituting $v$ (derived in the Extended information) and rearranging eq. (134) gives us stability condition in terms of the statistical properties of species interactions and the species survival fraction in the eLV ($\phi_N$):

$$\sigma^2_{A_{ij}} S\phi_N(1+\rho_A^{(D)}) < \frac{(\mu_{A_{ii}} - \mu_{A_{ij}})^2}{2} - \frac{1}{2}(\mu_{A_{ii}} - \mu_{A_{ij}})\sqrt{(\mu_{A_{ii}} - \mu_{A_{ij}})^2 - 4\sigma^2_{A_{ij}} S\rho_A^{(D)}\phi_N} + (\sigma_{A_{ii}} - \sigma_{A_{ij}})^2. \tag{136}$$

Finally, we plug in the expressions for the statistical properties of species interactions and self-inhibition derived in eq.s (131) and (130). This gives us a stability condition for the eLV in terms of the properties of species and resource pool:

$$\begin{aligned}
&\phi_N\gamma^{-1}\left(2 + \frac{4\mu_c^2}{\sigma_c^2 M} + \frac{\sigma_y^2}{\mu_y^2}\left(1 + \frac{\mu_c^2}{\sigma_c^2 M}\right)^2\right) \\
&< \frac{1}{2} - \frac{1}{2}\sqrt{1 - 4\phi_N\gamma^{-1}\left(1 + \frac{2\mu_c^2}{\sigma_c^2 M}\right)} \\
&+ \frac{1}{M}\left(\sqrt{2\left(1 + \frac{2\mu_c^2}{\sigma_c^2 M}\right) + \frac{\sigma_y^2}{\mu_y^2}\left(3 + \frac{6\mu_c^2}{\sigma_c^2 M} + \left(\frac{\mu_c^2}{\sigma_c^2 M}\right)^2\right)} - \sqrt{1 + \frac{2\mu_c^2}{\sigma_c^2 M} + \frac{\sigma_y^2}{\mu_y^2}\left(1 + \frac{\mu_c^2}{\sigma_c^2 M}\right)^2}\right)^2.
\end{aligned} \tag{137}$$

If we only consider the same leading-order $M$ terms as the CRM i.e., terms of order $\geq O(1/M)$, the stability condition becomes

$$\phi_N\gamma^{-1}\left(2 + \frac{4\mu_c^2}{\sigma_c^2 M} + \frac{\sigma_y^2}{\mu_y^2}\left(1 + \frac{2\mu_c^2}{\sigma_c^2 M}\right)\right) < \frac{1}{2} - \frac{1}{2}\sqrt{1 - 4\phi_N\gamma^{-1}\left(1 + \frac{2\mu_c^2}{\sigma_c^2 M}\right)} + \frac{1}{M}\left(\sqrt{2 + \frac{3\sigma_y^2}{\mu_y^2}} - \sqrt{1 + \frac{\sigma_y^2}{\mu_y^2}}\right)^2. \tag{138}$$

This stability condition is clearly not equivalent to the condition derived from consumer-resource model. Therefore, increasing the resource pool size will likely have a different impact on stability in the eLV to the CRM (although pool size could stabilise the eLV through alternative mechanisms).

Even in the thermodynamic limit where $M \to \infty$, the stability condition of the eLV is still fundamentally different to the CRM:

$$\phi_N\gamma^{-1}\left(2 + \frac{\sigma_y^2}{\mu_y^2}\right) < \frac{1}{2} - \frac{1}{2}\sqrt{1 - 4\phi_N\gamma^{-1}}. \tag{139}$$

where the CRM's stability condition in the thermodynamic limit is

$$\frac{\phi_N\gamma^{-1}}{\phi_R}\left(1 + \frac{\sigma_y^2}{\mu_y^2}\right) < 1. \tag{140}$$

Therefore, it is unclear whether the eLV is guaranteed to be stable when the CRM is in the thermodynamic limit. This insight, alongside evidence in from finite-sized systems, suggests that explicit resource dynamics are critical drivers of stability in the consumer-resource model.

### D.5 More simulated stability transitions in the effective Lotka-Volterra model

In this section, we give more examples of when the eLV fails to predict stability transitions in the CRM.

#### D.5.1 The eLV does not capture the unstable region of the CRM

In the main text, we showed an example of how the eLV cannot predict the stabilising effect of resource diversity for $\mu_c = 145$. To show this example captures a general phenomenon that the eLV cannot capture the CRM, we simulated and estimated the stability of eLV communities over different values of total resource consumption coefficients ($\mu_c$). For example, communities with high $\mu_c$ in consumer-resource models are completely unstable, but the eLV failed to capture this, or the stabilising effect of resource pool size. Instead, many eLV communities remained stable when $\mu_c$ was high and across a range of resource pool sizes (Fig. S5 A). This provides further evidence that resource dynamics are critical drivers of stability transitions in the consumer-resource model.

#### D.5.2 Resource pool size vs stability when the resource survival fraction is considered

In the main text when investigating stability relationships in the eLV, we did not consider how any resource dynamics shape species interactions. However, our derivation of the eLV from the CRM showed that species interactions and self-inhibition depend on emergent properties of resource dynamics, namely the resource survival fraction ($\phi_R$). We checked whether incorporating $\phi_R$ i.e., which resources survived or went extinct into the eLV's parameters restored the stabilising effect of resource diversity. However, we found the eLV still unable to predict the stability transitions observed in the CRM S5 B). This suggests that the change in exact resource abundances over time is the critical driver of stability transitions in the consumer-resource model, rather than being unable to measure which resources survive or go extinct without resource dynamics.

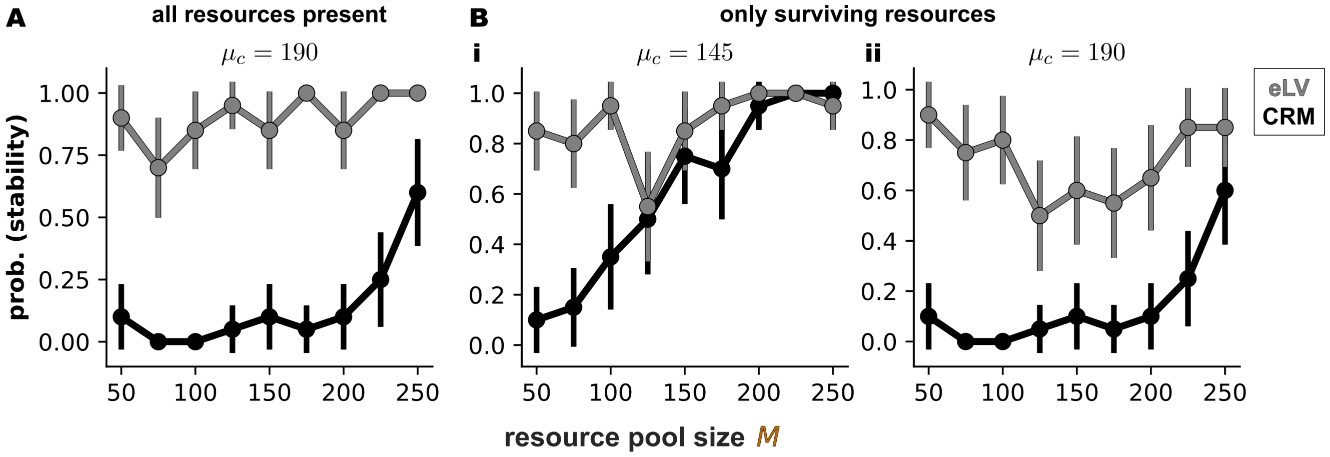

Figure S5: More examples of how the effective Lotka-Volterra model cannot predict the stabilising effect of stability. **A** The eLV cannot predict the emergence of persistent chaos or instability in the consumer-resource model. Shown are simulations for $\mu_c = 190$, assuming all resources survive. **B** Including surviving and extinct resources in the eLV does not rescue the stabilising effect of resource diversity. Shown are simulations for **i** $\mu_c = 145$ and **ii** $\mu_c = 190$. Species growth and competition coefficients are only parametrised using surviving resources (as species cannot grown or on interact through extinct resources).

##### D.5.3 The effects of different sources of interaction heterogeneity on stability

We also tested whether the eLV can capture how different sources of interaction heterogeneity induce opposing stability transitions in the CRM. In the CRM, increasing the variance in the consumption coefficient ($\sigma_c^2$) stabilised communities, whereas variance in yield conversion ($\sigma_y^2$) destabilised communities. Once again, the eLV could not quantitatively capture the stability transitions in the CRM (Fig. S6 A i and B i). The eLV did not undergo a stability transitions at the same critical values of $\sigma_c$ and $\sigma_y$ as the CRM, instead reaminig stable over a greater range of $\sigma_c$ and $\sigma_y$. In addition, the eLV was only weakly destabilised (the probability communities were stable $\neq 0$) at low values of $\sigma_c$ and high values of $\sigma_y$, respectively. Incorporating the resource survival fraction did not significantly improve eLV predictions — although the eLV appeared to better predict the effect of $\sigma_c$ on stability, it was worse at predicting the effect of $\sigma_y$ (so any apparent improvements were probably due to chance; Fig. S6 A ii and B ii). This provides further evidence that the resource dynamics themselves are critical drivers of stability relationships in the CRM.

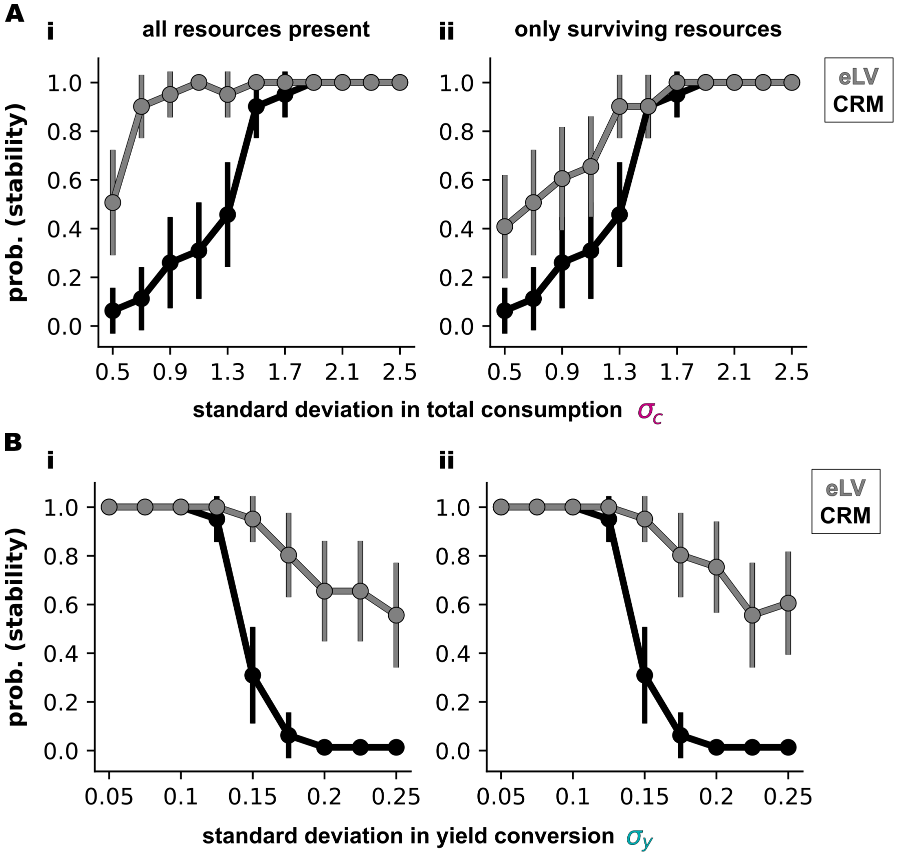

Figure S6: The eLV does not match the effect of different sources of interaction heterogeneity on stability in the CRM. **A** The effect of increasing the standard deviation in consumption $\sigma_c$ in the eLV and the consumer-resource model. **i** All resources are present **ii** only including surviving resources, obtained through running CRM simulations. **B** The effect of increasing the standard deviation in yield conversion $\sigma_y$ in the eLV and the consumer-resource model. **i** All resources are present **ii** only including surviving resources, obtained through running CRM simulations.

### E Numerical details

All codes for running simulations and solving self-consistency equations can be found in this repository: `https://github.com/JamilaRowlandChandler/CRM-Resource-diversity-vs-Stability`. This repository also contains notebooks with examples for running simulations and solving self-consistency equations.

#### E.1 Simulations

##### E.1.1 Consumer-Resource model

The model used in simulations is

$$\frac{\mathrm{d}N_i}{\mathrm{d}t} = N_i\left(\sum_{\alpha=1}^{M} y_{i\alpha}c_{i\alpha}R_\alpha - d_i\right) + \zeta, \quad \frac{\mathrm{d}R_\alpha}{\mathrm{d}t} = R_\alpha\left(b_\alpha - \frac{R_\alpha}{K_\alpha} - \sum_{i=1}^{S} c_{i\alpha}N_i\right) + \zeta, \tag{141}$$

where

$$y_{i\alpha} = \mu_y + \sigma_y z_{y,i\alpha}, \quad c_{i\alpha} = \frac{\mu_c}{M} + \frac{\sigma_c}{\sqrt{M}} z_{y,i\alpha}, \quad b_\alpha = 1, \quad d_i = 1, \quad K_\alpha = 1, \quad \zeta = 10^{-8}, \tag{142}$$

where $\zeta$ is the "migration rate". This term is added to reduce numerical instability in our simulations, which occurs when some species and resource abundances are very small but not extinct. However, adding this term does not change the type of dynamics observed in our system (e.g., fluctuations persist with and without migration). We neglect it in our analytical calculations because it is very small. (Alternatively, one could add an extinction threshold within the ODE solver, but the solve is often poor and does not agree with analytical calculations.)

We generated communities and simulated their dynamics in Python, mainly using *numpy* and scipy's ODE solver solve_ivp. For each set of parameter distributions (corresponding to each cell of the stability diagrams in Fig. 3A i and Fig. 4A), 20 communities were sampled. The parameters of each community were sampled from their respective normal distributions using *numpy.random.randn*. Community dynamics were then simulated from 2 sets of initial abundances, sampled from Uniform$(\text{min.} = \zeta, \text{max.} = 2/M)$ as in [10]. Dynamics were simulated for $t = 7000$ using the *LSODA* routine from scipy.integrate.solve_ivp, which is built for efficiently dealing with stiff (and non-stiff) ODE problems. An *unbounded growth* condition was also included to terminate the simulation when abundances grew unbounded. Hyper-parameters: the solver's relative tolerance was set to $10^{-7}$, absolute tolerates to $10^{-9}$. Dynamics were saved at $dt = 35$.

**Figure 3A i** – $M$ varied between 50 and 250 in increments of 25, $\mu_c$ varied between 100 and 250 in increments of 15. All other parameters were fixed to $\sigma_c = 1.6$, $\mu_y = 1$, $\sigma_y = 0.13069$, $b_\alpha = 1$, $d_i = 1$.

**Figure 5A** – $M$ varied between 50 and 250 in increments of 25. In **i**, $\sigma_c$ varied between 0.5 and 2.5 in increments of 0.2, and in **ii** $\sigma_y$ varied between 0.05 and 0.25 in increments of 0.075. Unless specified, other parameters were fixed to $\mu_c = 160$ $\sigma_c = 1.6$, $\mu_y = 1$, $\sigma_y = 0.13$ $(3.s.f)$, $b_\alpha = 1$, $d_i = 1$.

##### E.1.2 Effective Lotka-Volterra model

The protocol for simulating the eLV is very similar to the consumer-resource model.

$$\frac{dN_i}{dt} = N_i\left(r_i + \sum_{j=1}^{S} A_{ij}N_j\right) + \zeta. \tag{143}$$

**For simulations,**

$$r_i = \sum_{\alpha=1}^{M} y_{i\alpha} c_{i\alpha} \Theta(R_\alpha^{(ss)}) - \underbrace{1}_{d_i}, \quad A_{ij} = \sum_{\alpha=1}^{M} y_{i\alpha} c_{i\alpha} c_{j\alpha} \Theta(R_\alpha^{(ss)}), \quad \zeta = 10^{-8}, \tag{144}$$

To make our eLV and CRM simulations the most comparable, we obtained the effective Lotka-Volterra model for each CRM community. To generate parameters for each eLV, we extracted the consumption coefficients ($c_{i\alpha}$) and yield conversion factors ($y_{i\alpha}$) for each CRM community, and used these them to generate the species growth coefficients ($r_i$) and competition coefficients ($A_{ij}$). When we did not use any information about resource dynamics (like in the main text Fig. 4) so did not know which resources survive or go extinct, we used the coefficients from all resources to parametrise the eLV (i.e., $\Theta(R_\alpha^{(ss)}) = 1$ for all resources). When we used information on which resources survived and went extinct (like in Fig. S5 B), we only used the coefficients of surviving resources to parametrise the eLV. To determine which resources survived, we set an ad-hoc extinction threshold for each community — the threshold was set so that the proportion of resources exceeding it equalled the resource survival fraction from the cavity calculation ($\phi_R$). This choice was made to maximize the chance for the eLV to match stability properties with the CRM.

The simulation protocol was identical to Consumer-Resource models.

**Figure 4C** – eLVs were generated from Consumer-Resource models in Fig. 3A. For these CRM communities, $M$ varied between 50 and 250 in increments of 25. All other parameters were fixed to $\mu_c = 145$, $\sigma_c = 1.6$, $\mu_y = 1$, $\sigma_y = 0.13069$, $b_\alpha = 1$, $d_i = 1$.

#### E.2 Estimating species diversity

In main text fig. 3A, we defined the number of coexisting species as the number of species with abundances greater than extinction threshold $e$ at the end of simulations. We arbitrarily set $e$ to $10^{-4}$.

#### E.3 Estimating community stability

We determined the stability of each community by numerically estimating its maximum Lyapunov exponent. The maximum Lyapunov exponent is the largest eigenvalue of the community's Jacobian matrix. Therefore, it determines how sensitive each species and resource is to changes in abundances, i.e., whether community composition is stable in the face of perturbations.

In a continuous-time system, we can define the maximum Lyapunov exponent as follows [12]. Let's say we have two nearby trajectories of species and resource abundances $\mathbf{x}(t)$ in phase space. The first trajectory is denoted as $\mathbf{x}(t)$ and the nearby trajectory is denoted as $\mathbf{x}(t) + \boldsymbol{\delta}(t)$, where $\boldsymbol{\delta}(0)$ is the initial distance between the trajectories. In the limit of the initial distance $\boldsymbol{\delta}(0)$ going to zero and the time for which the two trajectories evolve going to $\infty$, we can define the maximum Lyapunov exponent $\lambda_{\max}$ in terms of the long-term distance $\boldsymbol{\delta}(t)$ between the trajectories as:

$$\lambda_{\max} = \lim_{\substack{\delta(0)\to 0 \\ t\to\infty}} \frac{1}{t} \log\left(\frac{|\boldsymbol{\delta}(t)|}{|\boldsymbol{\delta}(0)|}\right) \tag{145}$$

We can see that if $\lambda_{\max} < 0$, the absolute distance between trajectories $|\delta(t)|$ decays over time, indicating the system converges at the same stable steady steady. If $\lambda_{\max} > 0$, the trajectories diverge over time, indicating the system is unstable. We use this definition to calculate each community's maximum Lyapunov exponent using the algorithm from [13], summarised by Hrothgar (2015) and Koehler (2024).

##### Algorithm for estimating $\lambda_{\max}$

1. For the CRM, extract the final species and resource abundances from the end of simulations (detailed in Appendix D.1). For the eLV, extract the final species abundances. This is the "original trajectory" at $t = 0$.
2. Initialise a "perturbed trajectory". Perturb the original abundances by some small amount $\boldsymbol{\delta}(0)$ i.e., the Euclidean distance between the original and perturbed trajectory is $\boldsymbol{\delta}(0)$.
   - We set $\boldsymbol{\delta}(0)$ to $10^{-6}$.
3. Simulate the dynamics of the original and perturbed trajectory, computing the normalised (log) Euclidean distance between them $\log(\boldsymbol{\delta}(t))$ at each time step. Continue to simulate dynamics until $\log(\boldsymbol{\delta}(t))$ becomes approximately constant within some tolerance.
   - We found simulating dynamics for $t = 1000$ was sufficient to achieve this.
4. To estimate the maximum Lyapunov exponent $\lambda_{\max}$, fit a line to the log distance between trajectories $\log(\boldsymbol{\delta}(t))$ in the region where it varies with time. The best-fit slope is the estimated maximum Lyapunov exponent $\lambda_{\max}$.

#### E.4 Numerically solving the self-consistency equations

The self-consistency equations describing the consumer and resource abundance distributions — eq.s C.5.3 — cannot be solved analytically. Instead we needed to solve them numerically for each set of model parameter distributions. Please see `https://github.com/JamilaRowlandChandler/CRM-Resource-diversity-vs-Stability`. to see how this numerical routine is run.

To obtain solutions to our self consistency equations, we combined a non-linear least squares algorithm (`scipy.optimize.least_squares`) to find the local solution for the equations, and a basin-hopping routine (`scipy.optimize.basinhopping`) so that the solver traverses a large area of parameter space without getting trapped at a local minima. This routine has a long runtime, but can solve the self-consistency equations from any initial conditions within the bounds of said equations. (e.g., $v^{(N)} < 0$, $0 < \phi_N < 1$, etc.)

The loss function $\mathcal{L}(\text{SCE})$ minimised was the sum of the squared difference between the estimated value of each self consistency equation and the value calculated from the self-consistency equations listed in (C.5.3).

$$\mathcal{L}(\text{SCE}) = \sum_{p \in \text{SCE}} (p_{i,\text{estimated}} - p_{i,\text{calculated}})^2, \text{ where SCE is the set } \left\{\phi_N, \langle N \rangle, \langle N^2 \rangle, \phi_R, \langle R \rangle, \langle R^2 \rangle, v^{(N)}, \chi^{(R)}\right\}. \tag{146}$$

##### E.4.1 Solving for the stability boundary

To solve for stability boundary when varying some parameter $x$, we add the stability threshold to the loss function, $(\rho_{c,g}^2 - S^*/M^*) - 0$:

$$\mathcal{L}(\text{SCE}, x) = \sum_{p \in \text{SCE}} (p_{i,\text{estimated}} - (p_{i,\text{calculated}})^2 + (\rho_{c,g}^2 - S^*/M^*) - 0). \tag{147}$$

We also slightly modify our solving routine to improve runtime. We first choose a set of self-consistency equations solved by our global routine that are close to the stability boundary. We determine how close each set is to the stability boundary by calculating $\rho_{c,g}^2 - S^*/M^*$, for each set. Then, we locally solve the loss function using *scipy.optimize.least_ squares*.

### F Extended information

#### F.1 The geometric intuition for our model's stability condition

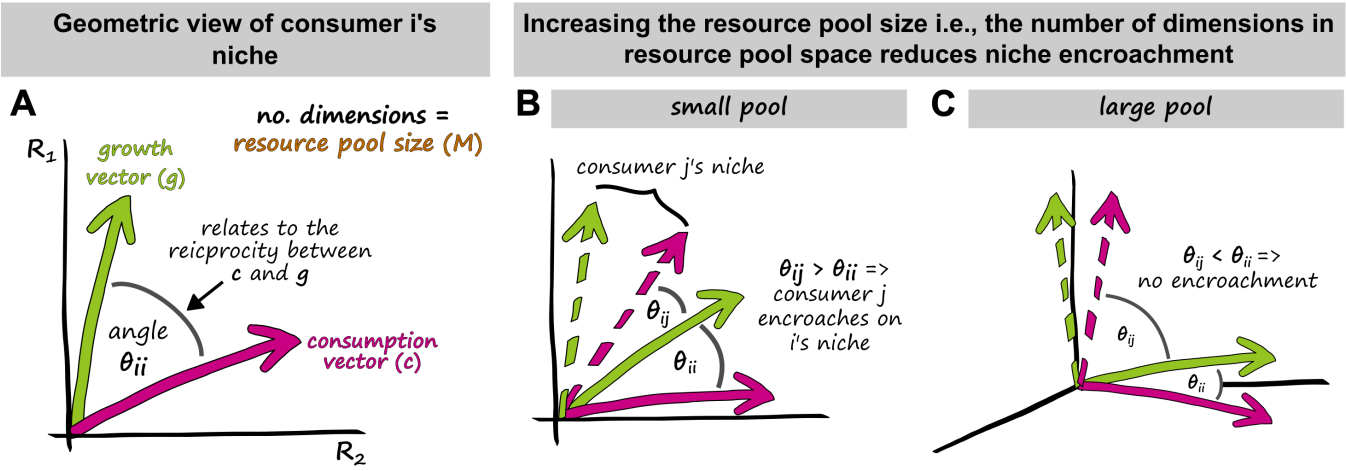

Figure S7: Geometric intuition for our model's stability condition **A** Geometric view of consumer $i$'s niche. In a resource space of $M$ dimensions (where $M$ is the resource pool size), the growth and consumption rates of consumer $i$ can be expressed as a growth $\mathbf{g}$ and consumption $\mathbf{c}$ vector. **B** When the resource pool size ($M$) is small, the likelihood that consumer $j$ will encroach on $i$'s niche is high. When $M$ is small, the reciprocity between consumer-resource interactions is low. This increases the chance that consumer $i$ has a more similar growth vector to consumer $j$'s consumption vector than its own ($\theta_{ij} < \theta_{ii}$), so $j$ encroaches on $i$'s niche. **C** Increasing the resource pool size/the dimensions of resource space increases reciprocity faster than the $\sqrt{\text{packing ratio}}$. This decreases the chance that consumer $i$ has a more similar growth vector to consumer $j$'s consumption vector than its own ($\theta_{ij} > \theta_{ii}$), so $j$ does not encroach on $i$.

In the main text, we discussed our biological interpretation of the stability condition: "interaction reciprocity sets a maximum bound on the number of species that can pack into a community without encroaching on each other's niches." We also showed that increasing the resource pool size $M$ generates species-rich and stable communities by increasing reciprocity faster than the $\sqrt{\text{species packing ratio}}$. Here, we provide a geometric intuition for these stability relationships, building on the work of Tilman [14], and Liu and colleagues [15].

Tilman [14] originally developed a geometric intuition for understanding when a community with two species and two resources would be stable. We will refer to the two species as consumer $i$ and $j$. He described the growth and consumption rates of a consumer $i$ at steady state as vectors ($\mathbf{g_i}$ and $\mathbf{c_i}$ respectively) projected onto resource space:

$$\mathbf{g_i} = N_i \begin{bmatrix} g_{i1}R_1 \\ \vdots \\ g_{iM}R_M \end{bmatrix}, \quad \mathbf{c_i} = N_i \begin{bmatrix} c_{i1}R_1 \\ \vdots \\ c_{iM}R_M \end{bmatrix}, \tag{148}$$

where the $M$ is the number of dimensions in resource space, or the resource pool size.

The angle between the growth and consumption vector measures how well they project onto each other, which is determined by the correlation between growth and consumption coefficients i.e., the interaction reciprocity. High reciprocity corresponds to a small angle (similar vectors), while low reciprocity corresponds to a large angle (Fig. S7 A).

Tilman then addressed the question: if consumers $i$ and $j$ coexist at steady state, when is this steady state stable? He found that the steady state would be stable when each consumer's growth vector aligns more closely with its own consumption vector than with other consumers' consumption vectors. In other words, when the angle between

$\mathbf{g}_i$ and $\mathbf{c}_i$, $\theta_{ii}$, is less than the angle between $\mathbf{g}_i$ and $\mathbf{c}_j$, $\theta_{ij}$. The steady state is unstable if $\mathbf{g}_i$ aligns more closely with $\mathbf{c}_j$ than $\mathbf{c}_i$ i.e., when $\theta_{ij} > \theta_{ii}$ (Fig. S7 B). This has a somewhat intuitive interpretation. Consumer $j$ consumes resources at rates that better match consumer $i$'s growth requirements (set by $\mathbf{g}_i$), enabling $j$ to displace $i$ from its own niche and destabilise the steady state. In other words, consumer $j$ **encroaches** on consumer $i$'s niche. Liu and colleagues [15] demonstrated that this geometric intuition extends to complex communities.

This geometric framework clarifies how increasing the resource pool size stabilises communities through its effects on interaction reciprocity and the species packing ratio. When the resource pool size is small, reciprocity is lower than the $\sqrt{\text{packing ratio}}$ (main text, fig. 3E). Low reciprocity causes poor alignment between a consumer's growth and consumption vectors in resource space. Because the $\sqrt{\text{packing ratio}}$ exceeds reciprocity, there is a decent chance that consumers will have growth vectors that align more closely with other consumers' consumption vectors than with their own; i.e., $\theta_{ij} < \theta_{ii}$ (Fig. S7 B). Therefore, consumers will likely encroach on each other's niches and destabilise community dynamics. Increasing the resource pool size increases both reciprocity and the $\sqrt{\text{packing ratio}}$, but reciprocity increases faster (main text, fig. 3E). High reciprocity causes a consumer's growth and consumption vector to align closely in resource space, creating more "room" for additional consumers to pack into the community without their growth vectors overlapping with each other's consumption vectors more than their own; i.e., $\theta_{ij} > \theta_{ii}$ (Fig. S7 C). Therefore, although more species coexist (higher $\sqrt{\text{packing ratio}}$) in larger resource spaces, they should not encroach on each other's niches (due to the higher reciprocity). Hence, large resource pools generate stable, species-rich communities.

#### F.2 Other stability transitions

Pictured here are the stability transitions induced by other parameters not discussed in the main text: mean yield conversion, mean and variance in consumer death rate, and mean and variance in intrinsic resource growth rate. To obtain these stability diagrams, we numerically-solved the self-consistency equations for each set of parameter distributions, then calculated how far communities are from the stability threshold (measured as interaction reciprocity $-\sqrt{\text{packing ratio}}$. These stability transitions are depicted in Fig. S8.

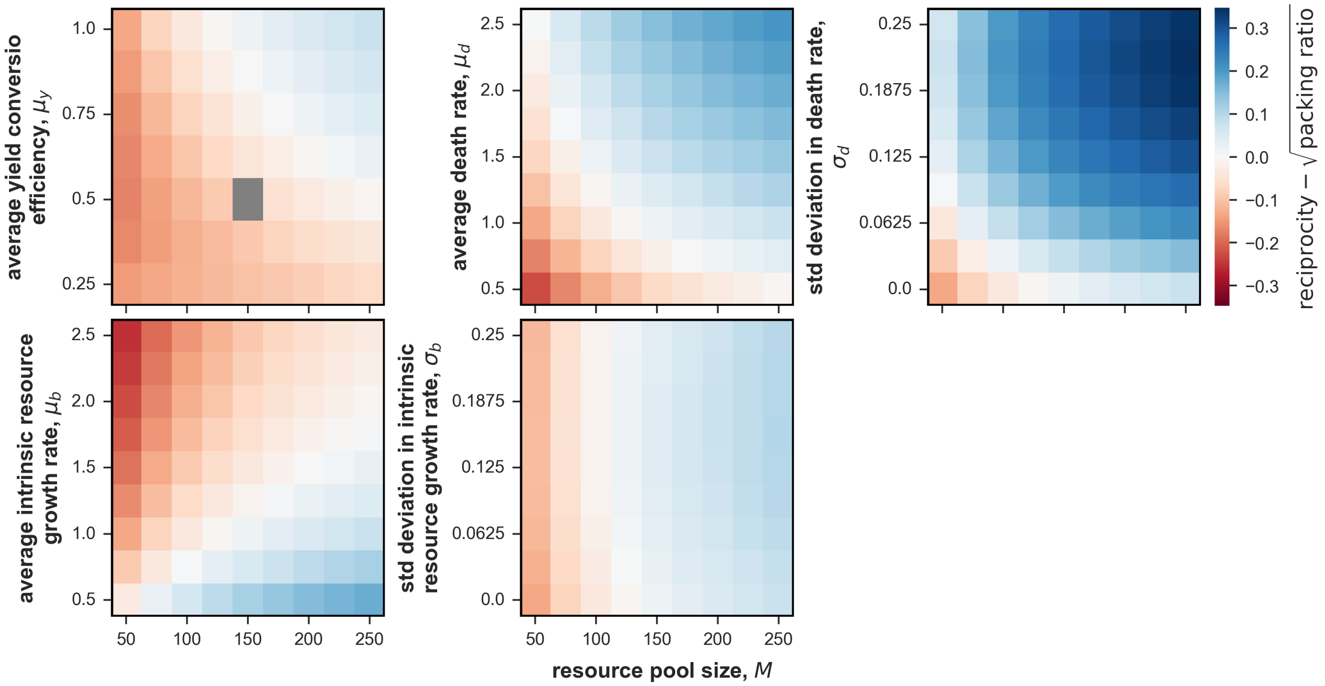

Figure S8: Additional stability transitions obtained by changing other parameters in the model, which were not detailed in the main text. Red cells indicate unstable regions, blue cells indicate stable regions while white cells demarcate the stability transition. Grey boxes indicate parameter combinations where our solver failed to converge.

#### F.3 The effects of the resource pool size on the self-consistency equations

Here, we plot the effect of the resource pool size ($M$) and the average total consumption coefficient ($\mu_c$) on the species and resource survival fractions ($\phi_N$ and $\phi_R$), the average species and resource abundances ($\langle N \rangle$ and $\langle R \rangle$), the fluctuations in species and resource abundances ($\langle N^2 \rangle$ and $\langle R^2 \rangle$), and the average susceptibilities of a species to a perturbation in its death rate and a resource to a perturbation on its intrinsic supply rate ($v^{(N)}$ and $\chi^{(R)}$) (Fig., S9).

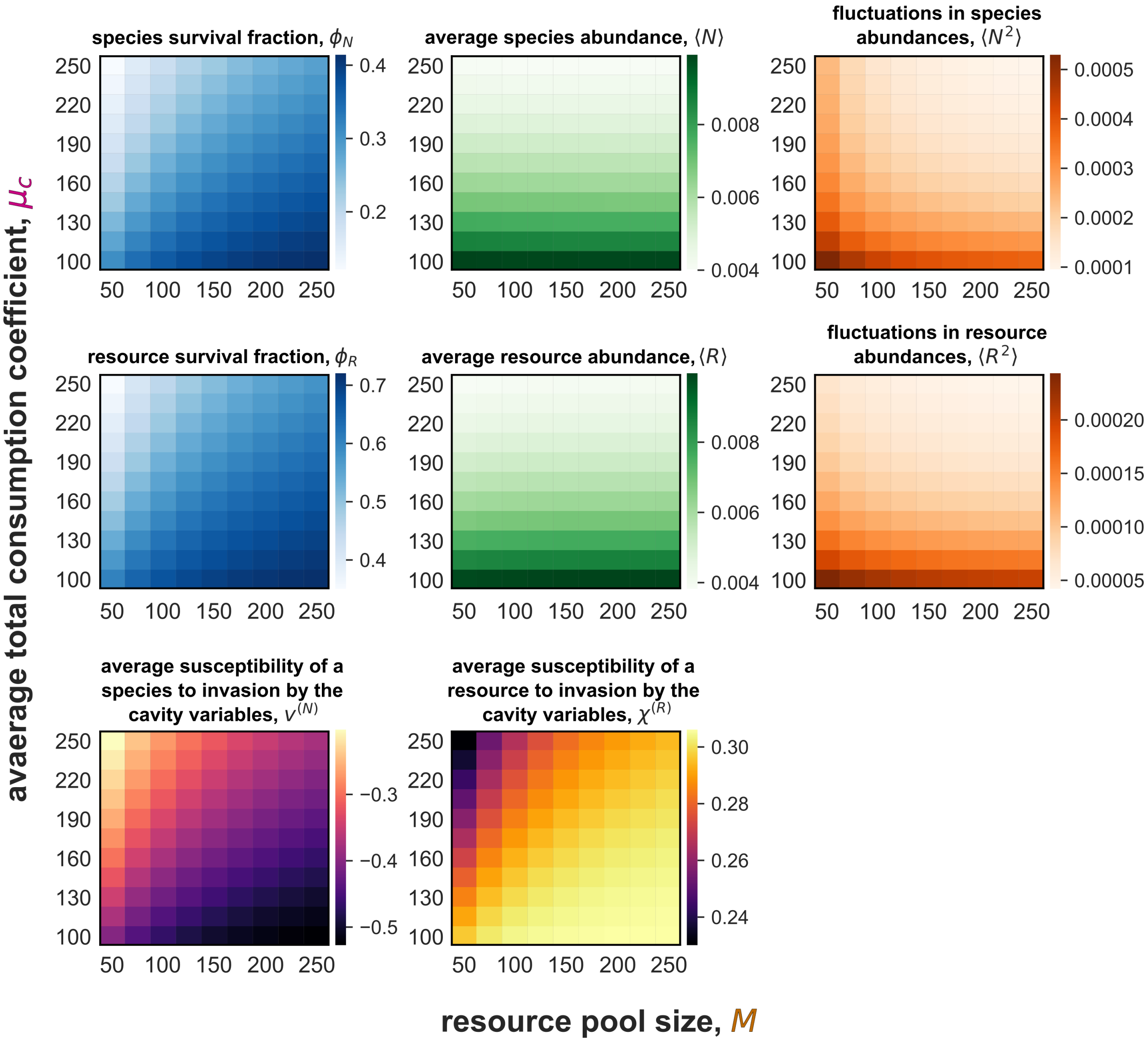

Figure S9: The effect of resource pool size and the average total consumption coefficient on the self-consistency equations. These values are obtained through numerically solving the analytical expressions for the self-consistency equations. See "Numerical methods" for how this is done.

#### F.4 The robustness of the positive diversity-stability relationship to changes in model formulation

##### F.4.1 Adding self-inhibition to match May's assumptions does not change our positive resource diversity-stability relationship

While our consumer-resource model reduces to the gLV under fast resource dynamics, it still differs from May's framework. Namely, self-inhibition and inter-consumer interactions are not independent, but arise from the same underlying mechanisms. Since May's stability criterion depends on the difference between inter-species interactions and self-inhibition, we modified our model to include direct consumer self-inhibition independent of resource-mediated interactions:

$$\frac{dN_i}{dt} = N_i \left( \sum_{\alpha=1}^{M} y_{i\alpha} c_{i\alpha} R_\alpha - d_i - \underbrace{A_{ii} N_i}_{\substack{\text{direct} \\ \text{self-inhibition}}} \right) + \zeta, \tag{149}$$

where $A_{ii} = 0.1$ for all species. This enabled us to more directly compare our model with May's framework to identify the primary drivers of stability. We hypothesised that our positive resource diversity-stability relationship would persist despite this addition, as explicit resource dynamics should continue to govern stability through consumer-resource interaction reciprocity and species packing ratio. As expected, adding direct self-inhibition did not alter the diversity-stability relationship (Fig. SI S10), confirming that resource dynamics remain the dominant factor controlling community stability in our model.

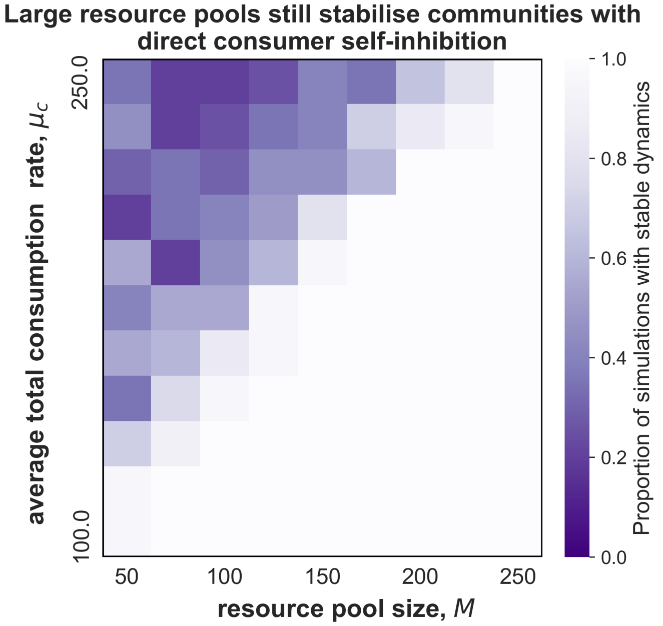

Figure S10: Simulated dynamics for the consumer-resource model with direct consumer self-inhibition

##### F.4.2 Large resource pools still stabilise communities under fixed total resource flux

Our model scales per-resource consumption rates with resource pool size ($c_{i\alpha} \propto 1/M$). This assumption contributes to the positive resource diversity-stability relationship observed in this study by making interaction reciprocity to depend on $M$. This scaling attempts to capture metabolism of substitutable resources. Such resources feed into common metabolic pathways (e.g., respiration), so total resource uptake saturates at $\mu_c \times$ total resource abundance regardless of resource diversity. However, the model formulation we present in the main text does not fully control resource flux because intrinsic resource growth rates ($b_\alpha$) remain constant. To test whether it is only the effect of increasing the pool of substitutable resources that stabilises communities, we also scaled the intrinsic resource growth rates $b_\alpha$ (from the consumers' perspective, the resource supply rates) by the resource pool size $M$, as $b_\alpha \propto 1/M$. We found that our positive diversity-stability relationship persisted, showing that our proposed biological mechanism can stabilise communities (Fig. SI S11).

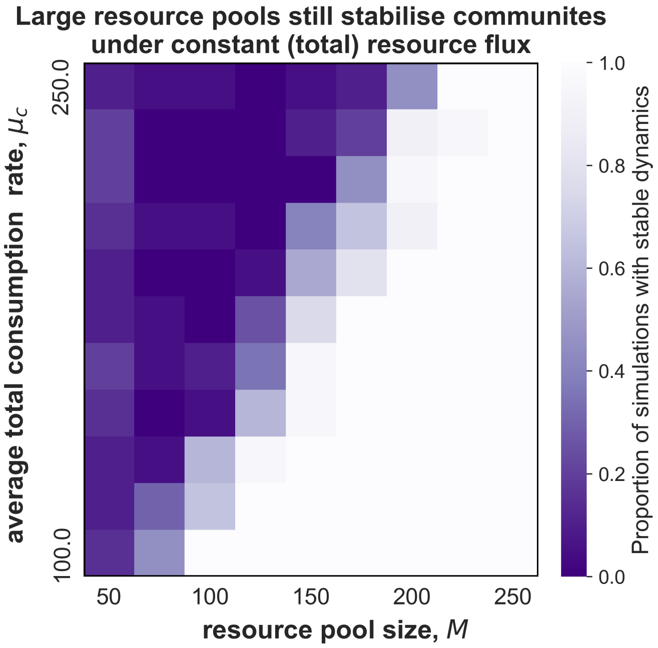

Figure S11: Simulated dynamics for the consumer-resource model with fixed total resource flux

### References

[1] Robert M. May. "Will a Large Complex System be Stable?" en. In: *Nature* 238.5364 (Aug. 1972), pp. 413–414. DOI: 10.1038/238413a0.

[2] Robert M. May. "Qualitative Stability in Model Ecosystems". en. In: *Ecology* 54.3 (May 1973), pp. 638–641. DOI: 10.2307/1935352.

[3] Guy Bunin. "Ecological communities with Lotka-Volterra dynamics". en. In: *Physical Review E* 95.4 (Apr. 2017), p. 042414. DOI: 10.1103/PhysRevE.95.042414.

[4] Emmy Blumenthal, Jason W. Rocks, and Pankaj Mehta. "Phase Transition to Chaos in Complex Ecosystems with Nonreciprocal Species-Resource Interactions". en. In: *Physical Review Letters* 132.12 (Mar. 2024), p. 127401. DOI: 10.1103/PhysRevLett.132.127401.

[5] Madhu Advani, Guy Bunin, and Pankaj Mehta. "Statistical physics of community ecology: a cavity solution to MacArthur's consumer resource model". eng. In: *Journal of Statistical Mechanics (Online)* 2018 (Mar. 2018), p. 033406. DOI: 10.1088/1742-5468/aab04e.

[6] Matthieu Barbier and Jean-Francois Arnoldi. *The cavity method for community ecology.* en. June 2017. DOI: 10.1101/147728.

[7] Itay Dalmedigos and Guy Bunin. “Dynamical persistence in high-diversity resource-consumer communities”. en. In: *PLOS Computational Biology* 16.10 (Oct. 2020). Ed. by James OâDwyer. Publisher: Public Library of Science (PLoS), e1008189. DOI: 10.1371/journal.pcbi.1008189.

[8] Wenping Cui, Robert Marsland, and Pankaj Mehta. “Effect of Resource Dynamics on Species Packing in Diverse Ecosystems”. en. In: *Physical Review Letters* 125.4 (July 2020), p. 048101. DOI: 10.1103/PhysRevLett.125.048101.

[9] Wenping Cui, Robert Marsland, and Pankaj Mehta. “Diverse communities behave like typical random ecosystems”. en. In: *Physical Review E* 104.3 (Sept. 2021), p. 034416. DOI: 10.1103/PhysRevE.104.034416.

[10] Emil Mallmin, Arne Traulsen, and Silvia De Monte. “Chaotic turnover of rare and abundant species in a strongly interacting model community”. en. In: *Proceedings of the National Academy of Sciences* 121.11 (Mar. 2024), e2312822121. DOI: 10.1073/pnas.2312822121.

[11] Stefano Allesina and Si Tang. “Stability criteria for complex ecosystems”. en. In: *Nature* 483.7388 (Mar. 2012), pp. 205–208. DOI: 10.1038/nature10832.

[12] Steven H. Strogatz. *Nonlinear Dynamics and Chaos.* en. 0th ed. CRC Press, May 2018. DOI: 10.1201/9780429492563.

[13] RÃŒdiger Seydel. *Practical Bifurcation and Stability Analysis.* en. Vol. 5. Interdisciplinary Applied Mathematics. New York, NY: Springer New York, 2010. DOI: 10.1007/978-1-4419-1740-9.

[14] David Tilman. *Resource Competition and Community Structure.* Princeton University Press, 1982. DOI: 10.2307/j.ctvx5wb72.

[15] Yizhou Liu et al. “Complex Ecosystems Lose Stability When Resource Consumption Is Out of Niche”. en. In: *Physical Review X* 15.1 (Jan. 2025), p. 011003. DOI: 10.1103/PhysRevX.15.011003.
